# Discovery of first-in-class inhibitors of the TRF1-TIN2 protein-protein interaction by fragment screening

**DOI:** 10.1101/2025.02.14.638354

**Authors:** Giacomo Casale, Manjuan Liu, Yann-Vaï Le Bihan, Oviya Inian, Ellie Stammers, John Caldwell, Rob L. M. van Montfort, Ian Collins, Sebastian Guettler

## Abstract

TRF1 is a subunit of the shelterin complex that binds to and protects the linear ends of chromosomes known as telomeres. Both genetic deletion and chemical inhibition of TRF1 have been shown to block the growth of lung carcinoma, glioblastoma, and renal cell carcinoma in mice without affecting mouse survival or tissue function, making TRF1 a potential therapeutic target in cancer^1–3^. Here, we report the discovery of a series of fragment hits that bind at the interface between the TRFH domain of TRF1 (TRF1_TRFH_) and a peptide of TIN2 (TIN2_TBM_), an interaction essential for the recruitment of TRF1 to shelterin, using X-ray crystallography (XChem) and ligand-observed NMR (LO-NMR) fragment screening. We discovered a first-in-class inhibitor of the TRF1-TIN2 interaction (compound **40**) that binds to TRF1_TRFH_ with a K_D_ of 29 μM (95% CI: 20 – 41 μM), displaces a TIN2 probe with an IC_50_ of 67 ± 28 μM, and expels TRF1 from purified shelterin. Aided by a novel crystal system of TRF1_TRFH_, we characterised fragments binding in a hotspot at the TRF1-TIN2 interface which will serve as a starting point for the structure-guided development of potent inhibitors of TRF1 protein-protein interactions to disrupt shelterin complex assembly.

## Introduction

Telomeres mark the linear ends of chromosomal DNA and are central to genomic integrity and control of the proliferative potential of the cell^4^. Human telomeres are characterised by several hundred kilobase pairs of tandem TTAGGG repeats, terminating in a shorter, single-stranded (ss) extension of the G-rich strand 50-400 nucleotides in length^4^. The resemblance of telomeres with DNA breaks requires the suppression of several DNA damage response (DDR) pathways that otherwise target double-strand (ds) DNA breaks and ssDNA, which make telomeres particularly susceptible to erroneous activation of the DDR and cell cycle arrest^4^.

The shelterin complex binds to both the double-stranded and single-stranded stretches of the telomere, responsible for the protection and length maintenance of telomeres^5^. Shelterin consists of six subunits: the two dsDNA-binders TRF1 and TRF2 (telomeric repeat-binding factor 1/2), the ssDNA-binder POT1 that forms a subcomplex with TPP1, the TRF2-associated RAP1 subunit and the central scaffolding subunit TIN2 that binds to TPP1, TRF1 and TRF2, thereby bridging the dsDNA- and ssDNA-binding modules (Fig. 1a)^6^.

**Figure 1:**
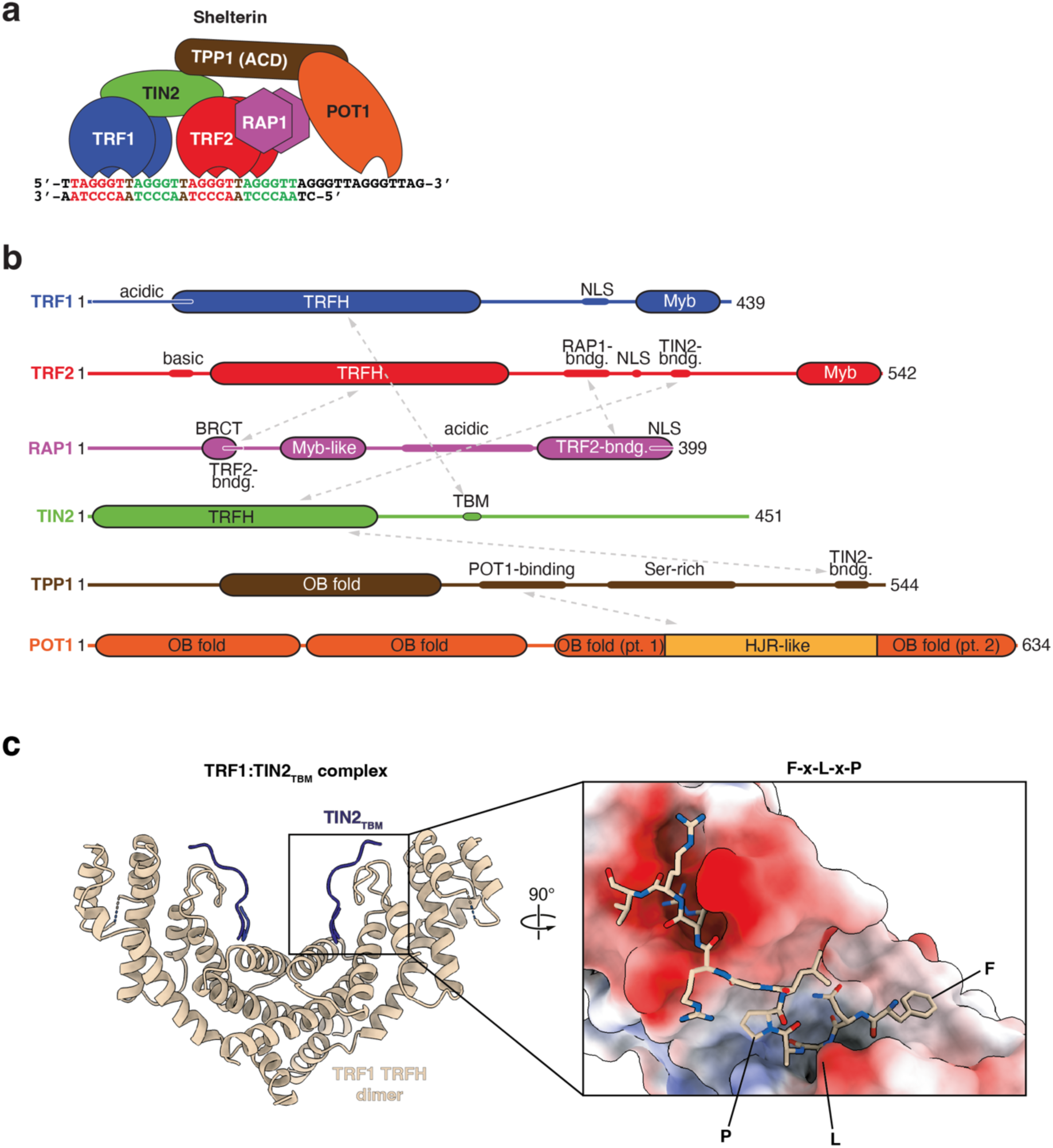
Shelterin subunits and domains. **(a)** Schematic representation of DNA-bound shelterin. **(b)** Domain architecture of shelterin subunits, with known inter-subunit interactions indicated by grey arrows. **(c)** Structural representation of the complex between TRF1_TRFH_ and TIN2_TBM_ (PDB 3BQO)^13^. Left-hand side panel is a cartoon representation of the overall complex, whilst the right-hand side panel is a close-up view, where TRF1_TRFH_ is shown in surface representation and coloured by electrostatic potential (red, negative; blue, positive). TIN2_TBM_ is shown in stick representation, and key residues of the F-X-L-X-P are highlighted.

In collaboration with telomerase, shelterin plays a central role in homeostatically maintaining telomere length^7^. Restored telomere length homeostasis is a hallmark of cancer initiation and progression, which makes shelterin a promising therapeutic target^8,9^. Whilst indirect inhibitors have been developed to weaken the integrity of the shelterin complex, the direct modulation of protein interactions within shelterin has been largely unexplored^10,11^.

TRF1 (Fig. 1a, b) binds to double-stranded telomeric repeats and mediates telomere protection, replication and extension^4,12^. It is recruited to shelterin through a domain:peptide interaction between its TRF homology (TRFH) domain, also responsible for dimerisation of TRF1 (TRF1_TRFH_), and a short linear motif of TIN2, known as its TRFH-binding motif or TBM (TIN2_TBM_; Fig. 1c)^13^. Versions of this TBM are also found in other telomere-associated proteins that are recruited to telomeres through a similar domain-peptide interaction with TRF1 characterised by the consensus motif F-X-L-X-P, such as the telomerase inhibitor Pin2/TRF1-interacting protein (PinX1)^6,14^, although it remains unknown how PinX1 binding to TRF1_TRFH_ can occur in tandem with TIN2_TBM_ binding to the same site.

Both genetic ablation and chemical inhibition of TRF1 telomeric localisation can block the growth of p53-null *K-Ras^G12V^*-induced lung carcinomas and PDGFA/B-induced glioblastomas in a telomere-length independent manner, remarkably without affecting mouse survival or tissue function^1–3^. Whilst these observations suggest that TRF1 may be a suitable therapeutic target, they also raise questions regarding the function of TRF1 in telomere length maintenance and protection. Garcia-Beccaria and coworkers identified two small molecules from a phenotypic screen that decreased TRF1 nuclear localisation, although it is not known whether the small molecules directly bind to TRF1^1^. To date, no small molecule exists that is known to inhibit the TRF1-TIN2 interaction. A chemical probe to directly modulate the incorporation of TRF1 into shelterin would be a valuable tool to interrogate telomere maintenance mechanisms and a potential starting point for the development of novel anti-cancer therapeutics targeting the shelterin complex. Moreover, such a small molecule could serve as a tool compound to interrogate the function of the ADP-ribosyltransferase tankyrase, which is proposed to expel TRF1 from shelterin, thereby promoting telomere extension^15–18^.

We therefore set out to discover and characterise small molecule fragments that bind to the TRFH domain of TRF1, aiming to disrupt the TRF1-TIN2 interaction and evict TRF1 from shelterin. Fragment-based drug discovery (FBDD) is a powerful tool to develop potent inhibitors of protein-protein interactions similar to that between TRF1_TRFH_ and TIN2_TBM_^19–21^. By screening small fragment-sized molecules (typically 120 – 250 Da), chemical space can be effectively sampled using a substantially smaller number of compounds than during typical high-throughput screens (HTS) with bigger, lead-like molecules. Whilst the resulting fragment hits would be expected to bind with a weaker affinity than HTS hits due to their smaller size, they can show a comparatively better ligand efficiency^22^. Fragment hits can then be grown, merged, or linked to develop high-affinity lead compounds with drug-like properties^20^. This has been demonstrated in the development of several clinical candidates and approved drugs^23–26^.

Here we used a combination of both LO-NMR and X-ray crystallographic fragment screening techniques to identify a first-in-class series of fragment hits that bind to TRF1_TRFH_ and inhibit the interaction between TRF1_TRFH_ and TIN2_TBM_ with low-micromolar potency as measured by a fluorescence polarisation (FP) competition assay. The hit matter characterised herein provides a firm basis for the development of potent, cell-active inhibitors of the TRF1-TIN2 interaction.

## Results and Discussion

### Primary fragment screening by X-ray crystallography

Due to its broad affinity detection range (nanomolar to millimolar) and unique ability to initiate rational drug design^27^, we carried out a crystallographic fragment screen against TRF1_TRFH_ at the XChem facility at Diamond Light Source^27^. Crystals of TRF1_TRFH_ with space group P3_1_21 were grown following previously published conditions^28^. They routinely diffracted to 2.0 – 2.2 Å, an improvement on the published resolution of 2.8 Å. In preparation for XChem screening, we tested the DMSO tolerance of the crystals and found that diffraction consistency could be maintained in the presence of 10% DMSO (data not shown).

For crystallographic fragment screening, we soaked 983 fragments from three fragment libraries (the DSiP fragment library^29^, the EUbOPEN DSiP extension library, and the FragLite library^30^) as singletons into TRF1_TRFH_ crystal drops. Of the 983 fragment-soaked crystals screened, 696 full datasets could be collected that were of sufficient quality for automated fragment detection analysis using the Pan-Dataset Density Analysis (PanDDA) algorithm^31^. We then inspected the PanDDA events (Fig. 2a) sequentially within PanDDA maps in Coot^32^ and identified 40 fragment hits across various sites, with 31 of those being located within the TIN2_TBM_ pocket. These data were extracted from PanDDA, and structures were corrected and refined in a more classical manner using Buster^33^ and Coot^32^. The resulting models and Sigma-A weighted 2mFo-DFc electron density maps were inspected to assess if the ligands were still observable before refining ligand occupancy with Buster (Supplementary Fig. 1). This process resulted in the high-confidence assignment of 19 of the 31 fragments (Fig. 2b). The binding poses of fragments that could be unambiguously assigned in conventional maps are also shown in Supplementary Table 2.

**Figure 2:**
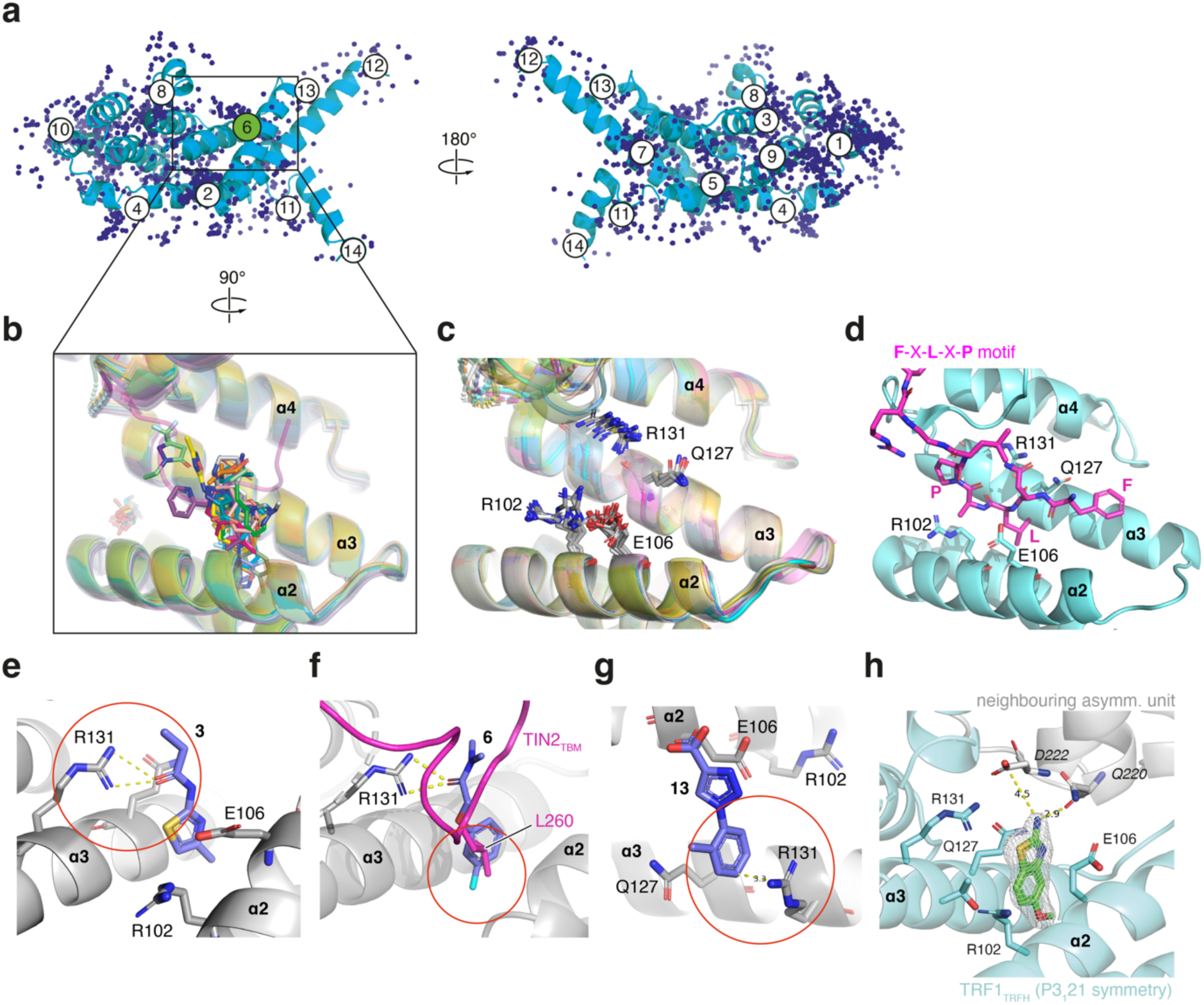
TRF1_TRFH_ Fragment screen by X-ray crystallography. **(a)** Cartoon representation of a TRF1_TRFH_ monomer with 1286 PanDDA events superimposed as blue spheres. Each circled number represents a PanDDA ligand binding site. The TIN2_TBM_-binding site, site 6, is highlighted in green. **(b)** Cartoon representation of 19 refined and superimposed TRF1_TRFH_ structures with hit fragments bound in the TIN2_TBM_ binding site. **(c)** The same structures as in b, but without the bound fragment hits, showing the relative positions of four key residues involved in fragment binding (R102, E106, Q127, R131). **(d)** Cartoon representation of the TRF1_TRFH_-TIN2_TBM_ crystal structure (PDB 3BQO)^13^ with the four residues involved in fragment binding shown as blue sticks and the TIN2_TBM_ shown as magenta sticks. **(e)** Example of an H-bond between R131 of TRF1_TRFH_ and an amide group of a hit fragment (**3**). **(f)** Example of a hit fragment (**6**) with a halide group buried in the leucine pocket of TRF1_TRFH_, with the TIN2_TBM_ peptide (PDB 3BQO)^13^ superimposed as a cartoon and L260 shown in stick representation. **(g)** Example of a cation-pi interaction between R131 of TRF1_TRFH_ and an aryl group of a hit fragment (**13**). **(h)** Crystal structure of the XChem hit fragment **5** bound to TRF1_TRFH_, with the neighbouring asymmetric unit shown in grey.

### Structural analysis of XChem fragment hits

Four key residues implicated in the binding of TIN2_TBM_ to TRF1_TRFH_ were also commonly involved in ligand binding, namely R102, E106, Q127, and R131 (Fig. 2c-h). Whilst the side chains of R131 and Q127 in helix ɑ3 assumed relatively fixed positions across the 19 ligand-bound structures, the side chains of E106 and R102 in the antiparallel helix ɑ2 showed more conformational heterogeneity (Fig. 2c). The range of positions adopted by E106 and R102 allowed TRF1_TRFH_ to accommodate a range of ligand binding modes, such as aromatic groups either parallel (e.g. Fig. 2e) or perpendicular to the helices ɑ2 and ɑ3 (Fig. 2g).

Interestingly, most ligand interactions with TRF1_TRFH_ (12/19 ligands) involved the side chains of R131 or Q127, whilst only three fragments formed direct interactions with the more flexible R102 and E106 residues. Ten out of the 19 fragment hits contained an amide or sulfonamide group that formed a hydrogen bond with either R131 or Q127 (Fig. 2e). In the hydrophobic pocket normally occupied by the leucine sidechain of the TIN2_TBM_ peptide (L260), a halide group was found in seven structures (Fig. 2f). Cation-pi interactions were also observed in five complexes between an aromatic ring of a fragment and either R102 or R131 (Fig. 2g).

### Orthogonal validation of XChem fragment hits

We next used a LO-NMR CPMG assay to orthogonally validate XChem hits in solution, aiming to eliminate false positives and ranking hit matter for follow-up studies. Compounds were tested as singletons and prepared in two samples containing either compound alone (sample 1) or compound + TRF1_TRFH_ (sample 2). Seven of the 19 XChem hits were shown to bind to the protein, but with only a modest average peak intensity reduction of 20 – 37% in the CPMG assay (Supplementary Table 2), suggesting that the fragments likely bound with weak (millimolar) affinities, at the very limit of the CPMG assay sensitivity.

Fragment **5** (Fig. 2h), which showed strong electron density after refinement and a CPMG peak reduction of 25%, was chosen for initial structure-activity relationship studies using commercially available analogues. However, small modifications to both the amino group and the methoxy group at position 5 of the benzothiazole core did not result in any significant increase or decrease in binding activity as measured by the CPMG assay, and no reliable LO-NMR K_D_ data could be obtained for the analogues (Supplementary Table 1). Attempts to soak TRF1_TRFH_ crystals with the commercially available analogues of the XChem hit **5** were also largely unsuccessful. Only one slightly smaller analogue (**22**) could be soaked into TRF1_TRFH_ crystals, albeit with an ambiguous binding mode. Closer inspection of the TIN2_TBM_ binding site revealed that a TRF1 molecule from the neighbouring asymmetric unit partially occluded the peptide binding site and participated in the stabilisation of several XChem hits. In the case of compound **5**, we observed a hydrogen bond between the amino group of the fragment and a backbone carbonyl in the neighbouring protein chain (Fig. 2h). This suggested that some of the XChem hits may have arisen from non-native, i.e., crystal-dependent, binding events. Additionally, the pocket normally occupied by the conserved phenylalanine residue of the TIN2 F-X-L-X-P motif was fully obstructed by the crystal contact with an adjacent asymmetric unit, further limiting the possibility of obtaining XChem hits outside of the leucine pocket.

### Primary fragment screening by LO-NMR

Whilst the XChem screen highlighted the possibility of obtaining hit matter in the TIN2_TBM_ leucine pocket of TRF1_TRFH_, it also raised questions about the tractability of the fragment hits obtained using the available crystal system, especially given the crystal contacts and poor hit confirmation rate by the orthogonal CPMG assay. It is known that different screening assays often identify different hits in fragment screening, and using a combination of biophysical and biochemical assays can help eliminate false positives and confirm strong hits^34,35^. We therefore set out to carry out an orthogonal fragment screen against TRF1_TRFH_ using relaxation-edited (CPMG) LO-NMR.

A total of 1097 compounds from the ICR biophysical fragment library^36^ were screened in pools of four structurally dissimilar molecules with non-overlapping resonances in the aromatic region of the ^1^H spectrum (5.5 – 9.5 ppm). The same fragment cocktails were also screened in the presence of TIN2_TBM_, a short peptide previously shown to bind to TRF1_TRFH_ with a dissociation constant of 0.3 μM, to allow for an initial assessment of binding site specificity (Fig. 3a)^13^. We defined a TIN2 site-specific hit arbitrarily as a compound showing an average peak intensity reduction of >35% in the presence of TRF1_TRFH_, and an average recovery of peak intensity of >20% in the presence of both TRF1_TRFH_ and TIN2_TBM_. Fragments with less than 20% peak intensity recovery in the presence of TRF1_TRFH_ and TIN2_TBM_ were classified as non-site-specific hits.

**Figure 3:**
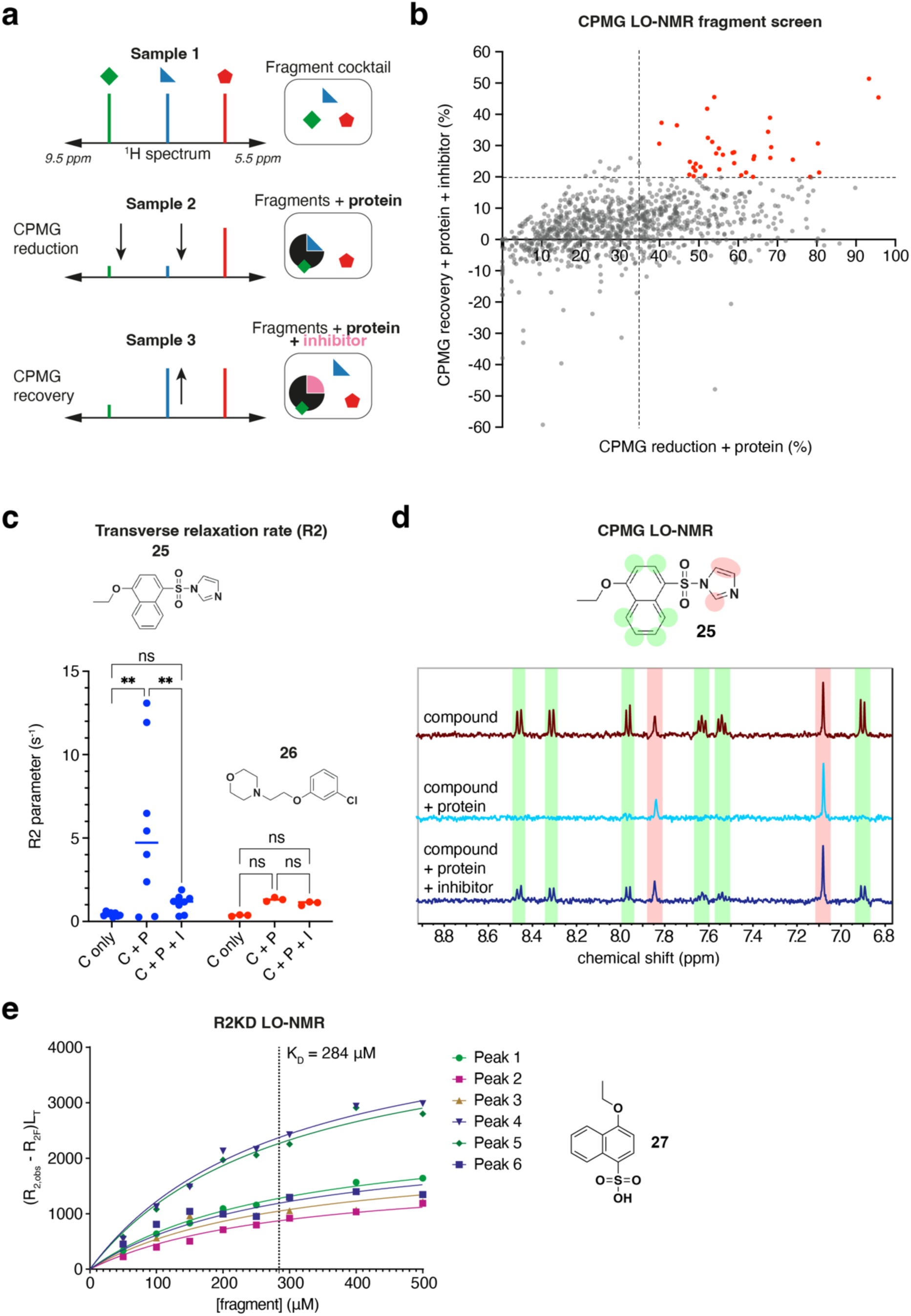
Identification and characterisation of TRF1_TRFH_ hit matter by LO-NMR. **(a)** Principle of the CPMG fragment screen. **(b)** Plot showing fragment hits from the CPMG NMR fragment screen, with primary hits classified as potentially TIN2 site-specific highlighted in red. **(c)** Measured R_2_ values from a single experiment for each proton peak of two fragment hits **25** and **26**. C, compound only; C + P, compound plus protein (TRF1_TRFH_); C + P + I, compound plus protein plus inhibitor (TIN2_TBM_ peptide). **(d)** CPMG spectra for compound **25** highlighting the lack of CPMG reduction for the imidazole group protons. **(e)** LO-NMR R2KD data for compound **27** from six aromatic proton peak relaxation rates (n = 1).

Of the 1092 fragments present in the CPMG-edited spectra, 702 were classed as non-binders, 266 as non-site-specific hits and 34 as site-specific hits (Fig. 3b). We then screened the 34 site-specific hits again as singletons to ensure that the observed binding was not influenced by interactions between fragments in the cocktail. However, only three fragments were validated as site-specific hits when tested as singletons, with 20 being reclassed as non-site-specific hits and eight as non-binders. One of the three site-specific hits later failed quality control, leaving only two specific hits (**25**, **26**, Fig. 3c) from the NMR fragment screen.

### Hit validation by transverse relaxation rate (R2) analysis

To further characterise the two site-specific hits from the NMR fragment screen, we measured the transverse relaxation rate of the fragments with and without TRF1_TRFH_. The transverse relaxation rates R2 of nuclei are heavily influenced by the rate of molecular tumbling, and hence by the size of the molecule. Fragments in solution tend to tumble rapidly, resulting in small R2 values (e.g. 0.5 – 2 s^-1^), whilst fragments bound to large macromolecules such as proteins tumble slowly, resulting in larger R2 values (e.g. 20 – 100 s^-1^)^37^.

By measuring the R2 values of fragments with and without protein present in the sample, a strong indication of the extent of protein binding can be obtained. Only one of the fragments (**25**) showed a significant increase in R2 upon addition of TRF1_TRFH_, indicative of protein binding (Fig. 3c). This increase in R2 was reversed upon addition of the TIN2_TBM_ competitor. Interestingly, the relaxation data also showed a stark contrast in relaxation rates between the protons attached to the naphthyl core of fragment **25** and those belonging to the imidazole group (Fig. 3c,d), indicating that the imidazole group did not bind to the protein. This suggested that the initial compound had likely degraded through hydrolysis to a sulfonic acid derivative **27** with the separate imidazole leaving group present in the mixture, a hypothesis that was confirmed by LC-MS analysis of the original compound stock (Supplementary Fig. 2). Nevertheless, given the strong binding activity of the fragment derivative **27** in the CPMG assay, a pure stock of **27** was synthesised and put forward for further characterisation using a quantitative LO-NMR assay to determine the dissociation constant.

### Measuring binding affinity by R2KD

Recent work by Liu and colleagues introduced a novel LO-NMR method that can be used to quantitatively determine the dissociation constant (K_D_) of a ligand for a protein from measured R2 values of ligand nuclei at a range of ligand concentrations^37^. The method, known as the R2KD assay, was used to characterise binding of NMR fragment screen hit **27** to TRF1_TRFH_ and revealed a K_D_ value of 284 μM (95% confidence intervals (CI): 230 – 534 μM, Fig. 3e).

### Orthogonal validation of NMR fragment hit 27

Encouraged by the relatively high binding affinity of compound **27** compared to expectations for fragments, we synthesised a series of analogues to explore the effect of modifying both the sulfonic acid and the ethoxy substituents on binding affinity (Table 2). Replacing the sulfonic acid at position R_1_ with a sulfonamide group (**33**) resulted in a drastic decrease in solubility of the fragment with a concomitant reduction of binding to TRF1_TRFH_ by CPMG, whilst the dimethyl sulfonamide **35** or the even bigger benzyl sulfonamide **36** showed an even poorer solubility and a loss of TRF1_TRFH_ binding (Table 2). The sulfonic acid replacement by a carboxylic acid (**34**) maintained a good solubility but led to a substantial loss of binding activity in the CPMG assay (Table 2). Similarly, substituting the ethoxy group at position R_2_ with the smaller methoxy (**28**) or hydroxy (**29**) groups did not hinder solubility but did reduce binding to TRF1_TRFH_ in the CPMG assay. However, substituting the ethoxy with larger aliphatic ether groups was better tolerated both in terms of TRF1_TRFH_ binding and aqueous solubility (**30**, **31**, **32**). The largest increase in binding was observed upon substitution of the aliphatic ether by a benzyl ether group (**32**), resulting in a five-fold increase in affinity as measured by the R2KD assay.

**Table 2:** SAR analysis of NMR screen hit analogues. LO-NMR CPMG values were determined from the average peak intensities of ≥4 ^1^H NMR peaks in the chemical shift region 5.5 – 9.5 of the ^1^H proton spectrum, from a single experiment. R2KD values are presented as a global KD value shared across ≥4 ^1^H NMR peaks from a single experiment with 95% confidence interval (CI) (profile likelihood) values shown.

| No. | R<sub>1</sub> | R<sub>2</sub> | Kin. Sol. (μM) | CPMG reduction (%) | CPMG recovery (%) | K<sub>D</sub> (μM) |
| --- | --- | --- | --- | --- | --- | --- |
| 27 |  |  | 810 | 96 | 49 | 284<br>(230 – 354) |
| 28 |  |  | 610 | 87 | 22 | - |
| 29 |  |  | 840 | - | - | >1000 |
| 30 |  |  | 860 | 83 | 4 | >1000 |
| 31 |  |  | 820 | 93 | 52 | 732<br>(560 – 995) |
| 32 |  |  | 850 | 100 | 29 | 60<br>(45 – 82) |
| 33 |  |  | 50 | 79 | 1 | - |
| 34 |  |  | 895 | 28 | 20 | - |
| 35 |  |  | <50 | - | - | - |
| 36 |  |  | 0 | - | - | - |

We then attempted to soak the NMR hit (**27**) and its analogues (**28 - 34**) into the TRF1_TRFH_ crystals to validate the binding mode of the series. However, the crystals disintegrated upon addition of the fragments at a range of concentrations, suggesting a ligand-induced disruption of the crystal packing. Given the close proximity of the neighbouring TRF1_TRFH_ molecule in the crystal packing to the TIN2_TBM_ binding site (Fig. 2h), we hypothesised that binding of the NMR hits and derivatives competed with crystal contacts, thereby degenerating the crystals. Thus, whilst we could obtain hits from both the NMR and XChem fragment screens, the cross-platform validation was generally limited, which prevented the rational design of larger, more potent analogues. We therefore turned our efforts towards finding a new crystal system that could accommodate larger ligands in the TIN2_TBM_ binding site, and where the binding mode of ligands would not be influenced by a neighbouring TRF1_TRFH_ molecule in the crystal packing arrangement.

### Discovery of a new TRF1_TRFH_ crystal system

To find a different crystal form of TRF1_TRFH_, we carried out a series of commercially available crystallisation screens and identified conditions that generated crystals in the P4_1_2_1_2 space group, different from the original P3_1_21 crystal system. These crystals diffracted consistently to 1.5 – 1.7 Å in their apo form. An initial inspection of the TIN2_TBM_ site showed that the closest neighbouring TRF1_TRFH_ molecule in this new crystal was further away than in the previous crystal system, this time exposing both the TIN2 leucine and phenylalanine binding pockets to the solvent (Fig. 4a,b; see also Fig. 4e,f). Furthermore, the loop region adjacent to the leucine pocket that typically accommodates three arginine residues of TIN2 was well-resolved and closely resembled that of the published TRF1-TIN2 co-crystal structure^13^; this was in contrast to the previous crystal system, in which F142 formed a cation-pi interaction with the sidechain of R131 (Fig. 4c,d). The new crystals were shown to tolerate DMSO concentrations of up to 10% without significant effects on resolution. We were therefore more confident that the new P4_1_2_1_2 crystal system would be better suited than the previous system for the identification and confirmation of TRF1-TIN2 PPI inhibitors.

**Figure 4:**
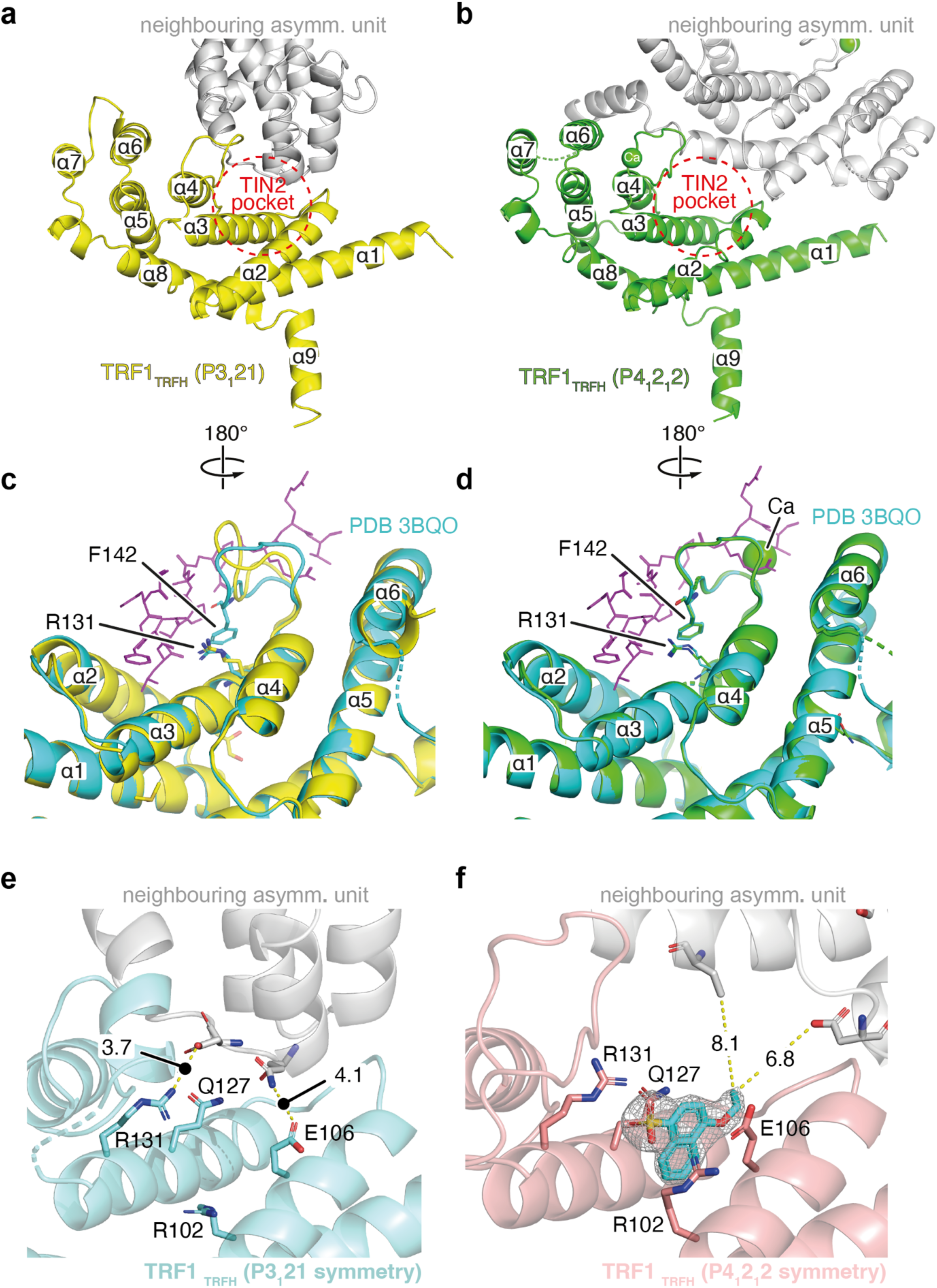
Comparison of two TRF1_TRFH_ crystal systems. (a, **b)** Cartoon representations of the P3_1_21 TRF1_TRFH_ monomer (yellow, a) and the P4_1_2_1_2 TRF1_TRFH_ monomer (green, b), with the neighbouring asymmetric units shown in grey. The TIN2 binding site is indicated by a red dotted circle. **(c, d)** Superimposed cartoon representations of TRF1-TIN2 complexes in turquoise and magenta (PDB: 3BQO) and the P3_1_21 TRF1_TRFH_ structure in yellow (c) versus the P4_1_2_1_2 TRF1_TRFH_ structure in green (d). **(e)** Structural representation of the vacant TIN2_TBM_-binding pocket from P3_1_21 TRF1_TRFH_ crystals. **(f)** Structural representation of NMR hit compound **27** bound to TRF1_TRFH_ in P4_1_2_1_2 TRF1_TRFH_ crystals with the 2mF_o_-DF_c_ electron density map for the ligand superimposed with a contour level of 1.0 RMSD. The distances in Å to the nearest residues of the neighbouring asymmetric unit (coloured in grey) are annotated as dashed yellow lines.

Most promisingly, we could successfully soak in and solve crystal structures with the NMR hit **27** and several of its analogues to fully characterise the binding modes within this fragment hit series. As shown in Fig. 4f, the naphthyl core of **27** is found deep in the leucine pocket of the TIN2_TBM_ site, with the negatively charged sulfonic acid group forming strong electrostatic interactions with the positively charged side chains of R102 and R131 on either side of the pocket. In the case of compound **32,** the benzyl group can be seen extending towards the TIN2 phenylalanine binding pocket where it sits almost perpendicular to the naphthyl core (Supplementary Fig. 3).

### Structure-guided optimisation of NMR screen hit

The successful soaking of NMR hit analogues **30 – 32** into TRF1_TRFH_ crystals validated their site-specific binding and opened up an opportunity to design more analogues *in silico.* Superimposing the published TRF1-TIN2 structure onto the crystal structure with compound **32** (Supplementary Fig. 3) highlighted the possibility of growing the compound towards the TIN2 phenylalanine binding pocket. We then used template-based docking^38^ to assess and rank various modifications of the benzyl group. The crystal structure of **32** bound to TRF1_TRFH_ was used as a template to dock a library of analogues in the target site with small substituents added to the benzyl group. The binding pose of the naphthylsulfonic acid core of the analogues was constrained to the leucine pocket, imitating the binding mode of **32**, whilst the substituted benzyl group was left unconstrained in the phenylalanine pocket. This docking exercise guided us to synthesise compounds **37 – 44**. As predicted *in silico*, the addition of relatively small, lipophilic groups to the meta position of the ring maintained or slightly improved affinity when measured by the R2KD assay (Table 3). However, some analogues with larger groups added at the meta position of the benzyl group, such as compounds **42** and **44**, could not be reliably measured by the R2KD assay due to poor signal-to-noise at ligand concentrations of 100 μM and above.

**Table 3.** SAR analysis of the benzoxyl group. . FP IC_50_ data are presented as the mean ± standard error (SEM) from 3 independent experiments done in technical duplicate. R2KD values are presented as a global K_D_ value shared across ≥4 ^1^H NMR peaks from a single experiment with 95% confidence interval (CI) (profile likelihood) values shown.

| No. | R | Kin. Sol.<br>(μM) | IC<sub>50</sub> (μM) | K<sub>D</sub> (μM) |
| --- | --- | --- | --- | --- |
| 32 |  | 850 | 152<br>(± 21) | 61<br>(45 – 82) |
| 37 |  | 460 | 200<br>(± 76) | 83<br>(59 – 117) |
| 38 |  | 570 | 146<br>(± 69) | 53<br>(31 – 90) |
| 39 |  | 320 | 166<br>(± 87) | 105<br>(58 – 201) |
| 40 |  | 320 | 67<br>(± 28) | 29<br>(20 – 41) |
| 41 |  | 590 | 23<br>(± 10) | 46<br>(28 – 73) |
| 42 |  | 260 | 80<br>(± 91) | - |
| 43 |  | 500 | 197<br>(± 79) | 144<br>(104 – 203) |
| 44 |  | 160 | 117<br>(± 53) | - |
| 60 |  | 715 | >1000 | - |

To complement the biophysical R2KD assay, we developed a fluorescence polarisation (FP) competition assay based on a TIN2_TBM_ peptide labelled with an N-terminal 5(6)- carboxyfluorescein (6-FAM) fluorophore. An initial titration of TRF1_TRFH_ against a fixed probe concentration of 25 nM revealed a K_D_ value of 0.4 ± 0.1 μM (Supplementary Fig. 4), which aligned closely to the published K_D_ of 0.3 μM derived from ITC^13^. The NMR hit analogues were then titrated against constant concentrations of TRF1_TRFH_ and TIN2-FAM (0.6 μM and 25 nM, respectively). As shown in Table 3, the IC_50_ values obtained correlated well with the R2KD affinity values, and highlighted compounds **40** and **41** as the most potent analogues for which we could obtain values in both assay formats (Fig. 5).

**Figure 5:**
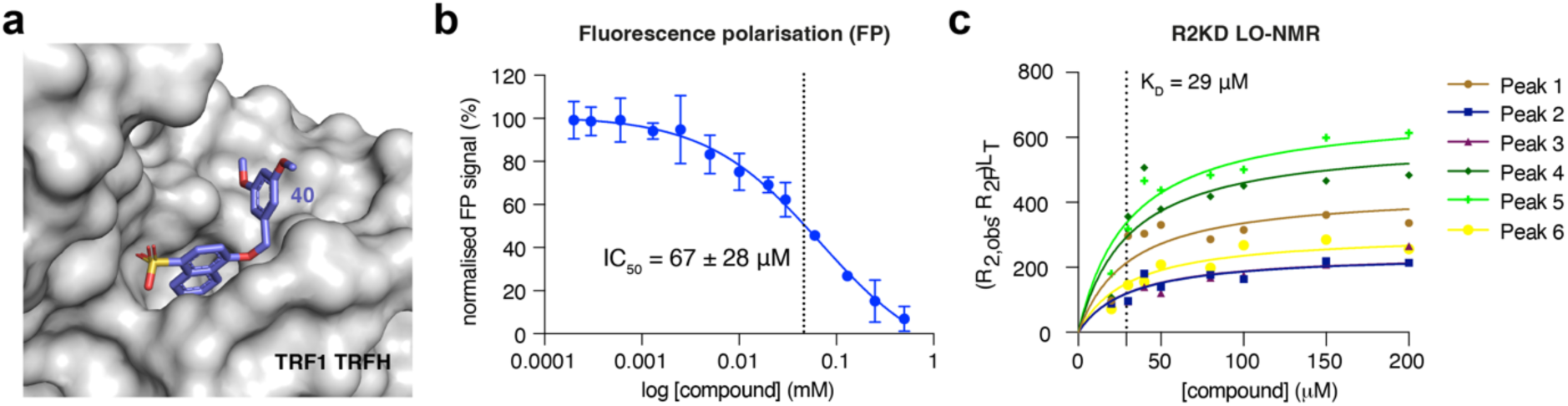
Biochemical and biophysical characterisation of compound 40. **(a)** Surface representation of TRF1_TRFH_ with bound compound **40** in stick representation. **(b)** Data from the FP competition assay showing TIN2-FAM displacement by compound **40** with an average IC_50_ value of 67 ± 28 μM derived from three independent experiments done in technical duplicate. **(c)** LO-NMR R2KD data showing a K_D_ of 29 μM (95% CI: 20 – 41 μM) derived across six aromatic proton peak relaxation rates (n = 1).

### Compound 40 evicts TRF1 from shelterin complex

We developed a mass-photometry based assay to evaluate the ability of compound **40** to disrupt the TRF1_TRFH_:TIN2_TBM_ interaction and subsequently evict TRF1 from shelterin. Purified shelterin, obtained by recombinant multi-protein expression in insect cells (see Materials and Methods), was incubated with increasing concentrations of compound **40**, and the resulting mass distribution was analysed by mass photometry. We observed an increase in a peak of 100 kDa (Fig. 6), corresponding to the full-length TRF1 dimer (Supplementary Fig. 5), with increasing concentrations of compound **40**, consistent with eviction of TRF1 from shelterin Quantification (as percentage of the 100-kDa TRF1 mass photometry peak relative to the total sample) demonstrated a concentration-dependent increase in eviction of TRF1, reaching a plateau at 250 μM of compound **40** (Fig. 6). Incubation of shelterin with an inactive compound (**60**) at the highest concentration of compound **40** tested (300 μM) did not show any increase in the TRF1 peak (Supplementary Fig. 5). Additionally, the mass photometry peak of shelterin shifted and reduced in size by approximately 100 kDa upon the addition of >250 μM of compound **40**, indicating the removal of TRF1 from the complex (Fig. 6).

**Figure 6:**
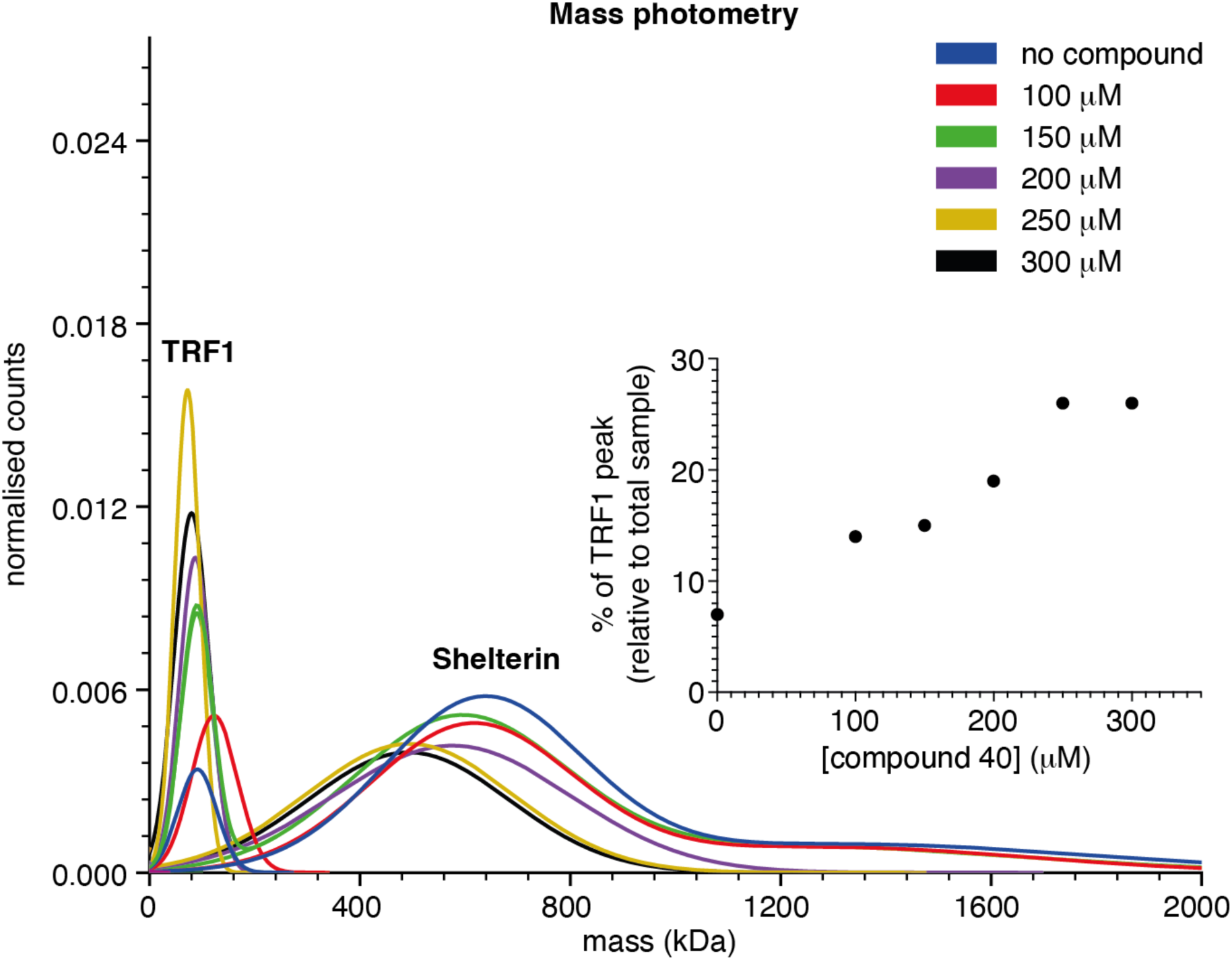
Characterisation of full-length TRF1 eviction from the shelterin complex by mass photometry. Mass photometry distribution represented as normalized Gaussian fits for shelterin at increasing concentrations of compound **40** showing eviction of TRF1 (n=1). The inset shows TRF1 eviction plotted as a percentage of TRF1 peak (of the total sample) vs compound concentration.

### Second crystallographic fragment screen using the P4_1_2_1_2 TRF1_TRFH_ crystal system

The results presented in Table 3 demonstrated the tractability of the NMR hit series and the possibility of obtaining more potent tool compounds through structure-guided optimisation. With a more exposed TIN2_TBM_ binding site and a native loop conformation adjacent to the site, the new P4_1_2_1_2 crystal system presented an opportunity to obtain more diverse, tractable fragment hits across the TIN2_TBM_ site, which could facilitate the NMR hit optimisation process through fragment growing, linking, and merging, and provide new hit matter. In light of this, we carried out a second XChem fragment screen against the new crystal system of TRF1_TRFH_.

A total of 1075 crystals were soaked with fragments from two libraries (DSi Poised library^29^ and ‘Probing Fragment All’ library), following the same screening pipeline as for the first XChem screen. PanDDA analysis of the 981 processable datasets resulted in 50 hits being identified across different sites of the protein, 28 of which were found in the TIN2_TBM_ binding site (Fig. 7a). After refinement of the 28 PanDDA hits, 19 could be confidently assigned in the conventional electron density maps (Fig. 7b; Supplementary Table 3). Four of the 19 refined hits were common to both XChem screens, although the binding pose of two of the hits had changed substantially (Table 4; Supplementary Fig. 6).

**Figure 7:**
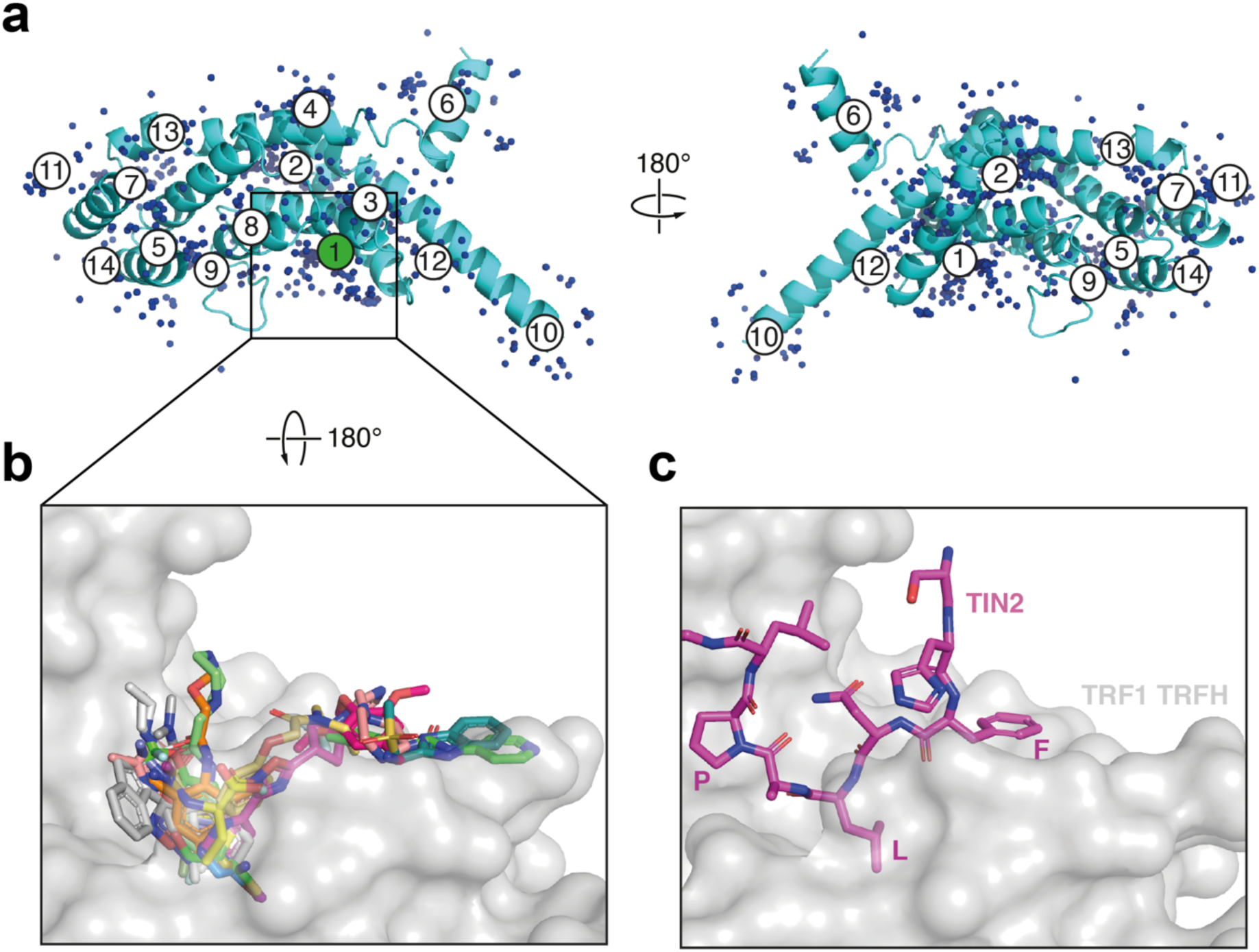
XChem Screen with P4_1_2_1_2 TRF1_TRFH_ Crystals. **(a)** Cartoon representation of TRF1_TRFH_ with 674 PanDDA events superimposed as blue spheres. Each circled number represents a PanDDA ligand binding site. The TIN2_TBM_-binding site, site 1, is highlighted in green. **(b)** Overlay of 19 XChem hits, shown in stick representation, bound in the TRF1_TRFH_ TIN2 pocket, shown in cartoon representation. **(c)** View of the TRF1_TRFH_ TIN2 pocket as in panel b, with TIN2_TBM_ in stick representation (PDB: 3BQO).

**Table 4.**
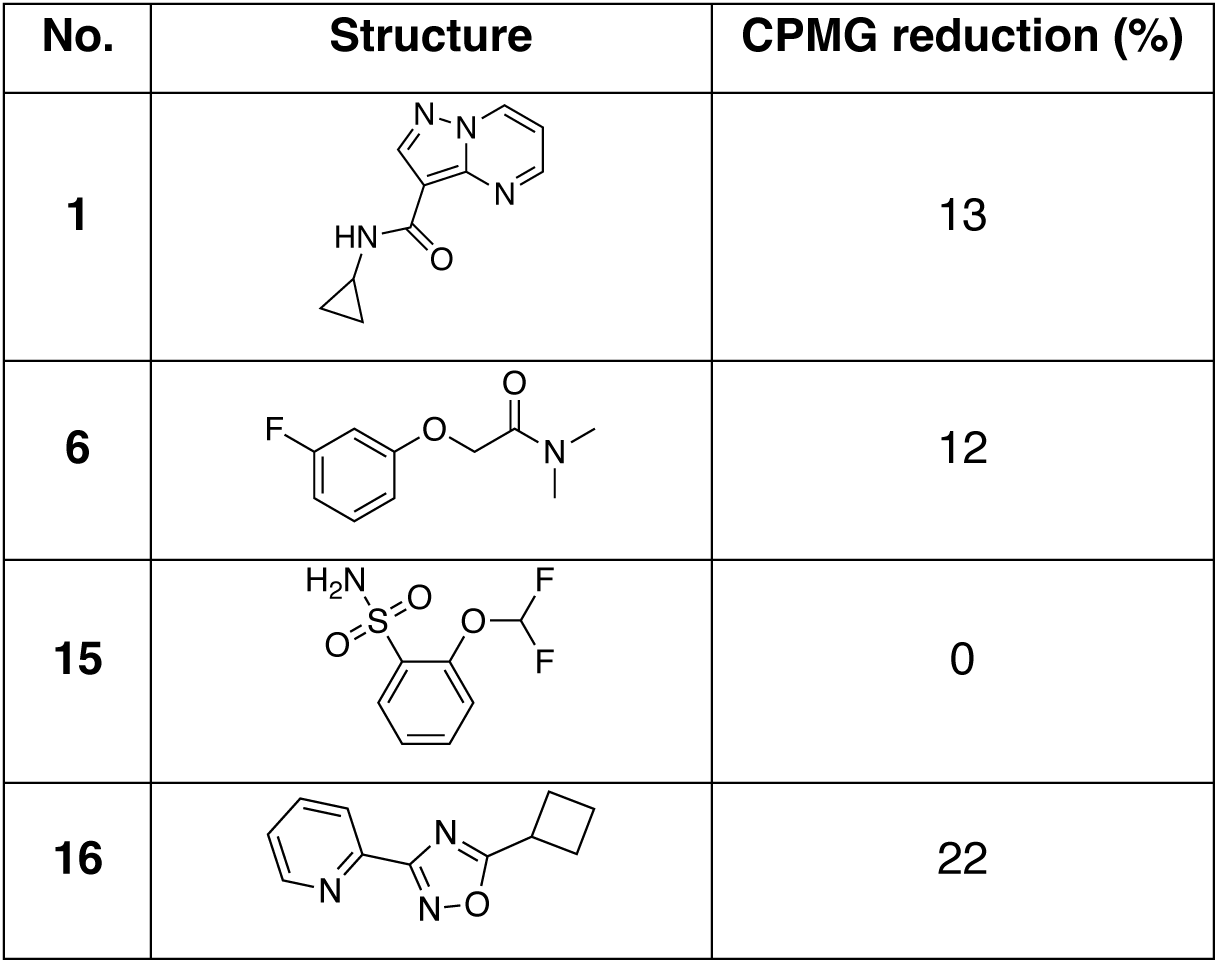
Fragment hits common to both XChem screens. . LO-NMR CPMG values were determined from the average peak intensities of ≥4 ^1^H NMR peaks in the chemical shift region 5.5 – 9.5 of the ^1^H proton spectrum, from a single experiment.

| No. | Structure | CPMG reduction (%) |
| --- | --- | --- |
| 1 |  | 13 |
| 6 |  | 12 |
| 15 |  | 0 |
| 16 |  | 22 |

We then carried out the CPMG assay, aiming to confirm the commercially available fragment hits from the second XChem screen. We excluded the four fragments we had already tested previously (**1, 6, 15, 16)** and one fragment that possessed no aromatic protons (**59**). Only six new fragments showed a peak reduction of greater than 20% in the CPMG assay (Supplementary Table 2), a similar degree of hit validation to the first XChem screen. However, unlike the hits obtained from the P3_1_21 XChem screen, the hits from the P4_1_2_1_2 XChem screen spanned a larger surface area of the TIN2_TBM_ binding site (Fig. 7b,c), and most did not appear to form any interactions with TRF1_TRFH_ molecules from neighbouring asymmetric units. This diversity of hit matter covering the leucine pocket and the phenylalanine pocket of the TIN2_TBM_ peptide will provide a firm starting point for efforts aimed at linking and merging hits.

### Conclusions and Discussion

Here we report the discovery of a set of hit fragments that bind to the TIN2_TBM_ binding site of the TRFH homodimerisation domain of TRF1. Using a combination of X-ray crystallography and LO-NMR methods, we identified fragments binding in both the leucine pocket and the phenylalanine binding pockets of the TIN2 binding site, two sites crucial for TIN2_TBM_ engagement with TRF1_TRFH_ through the F-X-L-X-P motif^13^. The cluster of fragments found to bind in the leucine pocket across both TRF1_TRFH_ XChem screens strongly validates the ligandability of this site and the possibility of developing potent inhibitors of the TRF1-TIN2 protein-protein interaction.

From a CPMG-based LO-NMR fragment screen, we identified one naphthalene sulfonic acid fragment hit that binds to TRF1_TRFH_ with three-digit micromolar affinity as measured by the R2KD assay. Through an initial exploration of the structure-activity relationship (SAR), we found that modifications of the ethoxy group were well tolerated, resulting in an increased affinity, whilst substitution of the sulfonic acid moiety was less well tolerated, mostly by negatively affecting the aqueous solubility of the fragments. The discovery and optimisation of a new crystal system of TRF1_TRFH_ enabled routine soaking of hits into the TIN2_TBM_ binding site. We show that the TRF1_TRFH_ ligand structures can serve as a basis for *in silico* design of more potent analogues through template-based docking. An FP competition assay was also established to biochemically validate displacement of the TIN2 peptide by the compounds. We further showed that compound **40**, identified as the most potent analogue among the compounds tested, evicted full-length TRF1 from the shelterin complex by disrupting the TRF1_TRFH_:TIN2_TBM_ interaction.

Whilst the analogues synthesised and presented in Table 3 do not represent a full-scale medicinal chemistry effort, they nevertheless highlight a proof-of-concept approach to develop high-affinity inhibitors of the TRF1-TIN2 interaction. Further SAR studies will also be required to investigate the selectivity of the inhibitors for TRF1_TRFH_ over the paralogous TRFH domain of TRF2 (TRF2_TRFH_). We have set up a robust assay cascade to enable full-scale lead optimisation and overcome some limitations in the current lead compounds. For example, the sulfonic acid component provides a substantial barrier to cell permeability and will need to be modified or replaced prior to cell-based experiments. Interestingly, several fragment hits originating from our second XChem screen possess alternative groups to the sulfonic acid that interact with R131 and R102, which opens up opportunities to link and merge hit matter found across the TIN2 leucine and phenylalanine binding pockets. Future work will therefore focus on combining XChem hit matter with the main NMR hit series through structure-guided merging and linking of fragments to design more potent, drug-like, and cell-permeable tool compounds.

## Materials and Methods

### Protein expression and purification

Purification of TRF1_TRFH_ domains was carried out as described previously with some modifications^13,28^. In brief, a pET-His_6_-MBP-Asn_10_-TEV (1C) expression vector (gift from Dr Scott Gradia, UC Berkeley, via Addgene; #29654) with human TRF1_TRFH_ cDNA (encoding residues 47-268, UniProt identifier P54274-1, codon-optimised for *E. coli* and synthesised by GenScript) downstream of an N-terminal His_6_-MBP tag was used to transform *E. coli* BL21-CodonPlus(DE3)-RIL (Stratagene) for protein expression.

A single clone of transformed *E. coli* cells was grown in Terrific Broth to exponential phase before induction with 0.5 mM IPTG and overnight incubation at 16 °C. Next day, cells were harvested by centrifugation and the pellets flash-frozen and stored at -80 °C. The cells were resuspended in lysis buffer (50 mM Tris pH 8.0, 500 mM NaCl, 10 mM β-mercaptoethanol, 0.5 mM AeBSF, 10 μL DNase, 1x Complete protease inhibitor tablet [Roche]). The resuspended cells were lysed by sonication and then centrifuged to remove cellular debris. The resulting cell lysate was filtered through a 0.2-μm filter and loaded onto a 5-mL HisTrap HP column (GE Healthcare/Cytiva) equilibrated in wash buffer (50 mM Tris-HCl pH 8.0, 500 mM NaCl, 1 mM TCEP, 25 mM imidazole). Once the lysate had been loaded, the column was washed with 20 column volumes (CV) wash buffer, and the His-tagged TRFH domain was eluted using a linear gradient from 0 to 100% elution buffer (50 mM Tris pH 8.0, 500 mM NaCl, 1 mM TCEP, 300 mM imidazole) over 25 CV.

The eluted protein was dialysed overnight in the presence of His-tagged TEV protease to cleave the His_6_-MBP tag. The dialysis buffer (50 mM Tris-HCl pH 8.0, 500 mM NaCl, 1 mM TCEP) was exchanged twice the next morning after 3-h intervals. To remove the cleaved tag and TEV protease, the dialysed protein was loaded onto an equilibrated 5-mL HisTrap HP column, and the flowthrough was collected. Step-wise elutions of 2.5%, 5% and 7.5% elution buffer were performed to remove any cleaved target protein bound tag-independently to the affinity column. Fractions containing the TRFH domain as confirmed by SDS-PAGE were pooled and concentrated by centrifugation on a Vivaspin Turbo 15 centrifugal concentrator (10 kDa MWCO) to a final volume of 5 mL. The resulting protein was centrifuged to remove any aggregates and loaded onto a Superdex 200 16/600 size-exclusion column equilibrated in SEC buffer for crystallisation (10 mM Tris pH 8.0, 200 mM KCl, 0.5 mM PMSF, 1 mM TCEP)^28^ or NMR (50 mM mono/dibasic sodium phosphate buffer pH 8.0, 150 mM NaCl, 1 mM TCEP). The purified TRF1_TRFH_ domain was then concentrated by centrifugation on a Vivaspin Turbo 15 centrifugal concentrator (10 kDa MWCO) to either ∼3 mg/mL for NMR studies or ∼30 mg/mL for crystallisation before flash-freezing and storage at -80°C (Supplementary Fig. 7). Shelterin complex was recombinantly expressed and purified from *Spodoptera frugiperda* (Sf9) insect cells. Briefly, the Shelterin genes TRF1, TRF2, RAP1, TPP1, POT1 and (StrepII)_2_- TEV-tagged TIN2 were codon-optimised for *E. coli* and synthesised by GenScript. Genes were cloned using nicking cloning^39^ into a single expression construct in the pACEBac1 vector. Baculoviruses were generated using the MultiBac system^40–42^ with the bacmids made using DH10^EMBacY^ cells (Geneva Biotech). Around 1.2 - 1.6 L of Sf9 cells were grown in suspension at 27 °C and 130 rpm, to a density of 1 x 10^6^ cells/mL and infected with 1:500 v/v of the baculovirus to the culture volume. The cultures were incubated for 72 h at 27 °C until the YFP signal was detected in >80% of cells and subsequently harvested by centrifugation, flash-frozen and stored at -80 °C. The cells were resuspended in lysis buffer (50 mM HEPES-NaOH pH 8.0, 300 mM NaCl, 10% glycerol, 1 mM MgCl_2_, 5 mM β-mercaptoethanol 4 μg/ml Avidin, 500 U Pierce^TM^ Universal nuclease for cell lysis, 1 mM AeBSF, 1 Roche cOmplete^TM^ EDTA-free protease inhibitor tablet per 100 mL of lysis buffer), lysed by sonication and clarified by centrifugation at ∼35,000 x g for 45 min. The supernatant was filtered using a 0.45-μm filter before being applied to a pre-equilibrated 5-mL StrepTrap column (Cytiva). The column was extensively washed with the wash buffer (50 mM HEPES-NaOH pH 8.0, 300 mM NaCl, 10% glycerol, 1 mM TCEP), and elution was performed with wash buffer containing 10 mM desthiobiotin. The affinity-purified Shelterin was then subjected to multiple rounds of size-exclusion chromatography (SEC) using the Superose 6 Increase 10/300 column equilibrated with the SEC buffer (50 mM HEPES pH 8.0, 300 mM NaCl, 5% glycerol, 1mM TCEP) to further purify and polish the complex. SEC fractions were flash-frozen and stored at -80 °C until further analysis (Supplementary Fig. 8). (StrepII)_2_-tagged TRF1 was expressed and purified from approximately 800 mL of Sf9 cells under the same conditions as outlined for the full Shelterin complex.

### Crystallisation of TRF1_TRFH_

TRF1_TRFH_ crystals in the P3_1_21 space group were grown by sitting drop vapour diffusion in SWISSCI 3-drop plates using similar crystallisation conditions to those described previously^28^. The total drop volume was 300 nL, consisting of 150 nL TRF1_TRFH_ (31.2 mg/mL) and 150 nL reservoir solution (1% PEG8000, 100 mM MES pH 6.0, 1 - 12 mM Mg(OAc)_2_, 10 - 15% glycerol), whilst the reservoir volume was 35 µL. Crystallisation plates were set up using an SPT Labtech mosquito Xtal2 liquid handler. Cuboid-shaped crystals grew to their maximum size (300 μm across) within 24 h. Crystals were cryoprotected through the addition of 180 nL 40% glycerol to the crystal drop prior to harvesting, giving a final glycerol concentration of 28.8 – 35.5% (accounting for the glycerol already in the crystallisation conditions). The crystals were then flash-frozen in liquid nitrogen, and data were collected at the Diamond Light Source beamlines I04 and I04-1 and processed automatically using the Xia2 Dials pipeline^43,44^. The TRF1_TRFH_ structure was solved by molecular replacement in Phaser MR^45^ within CCP4^46^ using the previously published TRF1_TRFH_ structure as a search model (PDB: 1H6O)^28^, after having first stripped the search model of solvent molecules and ligands. Structure refinement was done using Refmac^47^ and Buster^33^, and model correction and building was done using Coot 0.9.6^32^.

TRF1_TRFH_ crystals in the P4_1_2_1_2 space group were first identified from a PEGRx HT crystallisation screen (Hampton Research) using the sitting drop vapour diffusion method in SWISSCI 3-drop plates. Crystal drops were prepared by adding 150 nL of 28.6 mg/mL TRF1_TRFH_ to 150 nL of reservoir solution (100 mM MES pH 6.0, 35 – 45% PEG200, 50 mM CaCl_2_), against 35 μL reservoir, and reached their maximum size (200 – 300 μm across) after 5 d. Crystals were cryoprotected by addition of ethylene glycol at a final concentration of 10% prior to harvesting. The crystals were then flash-frozen in liquid nitrogen, and data were collected at the Diamond Light Source beamlines I04 and I04-1 and processed automatically using the Xia2 pipeline^43^. The TRF1_TRFH_ structure was solved by molecular replacement in Phaser MR^45^ using the in-house refined P3_1_21 TRF1_TRFH_ structure as a search model (solvent molecules and ligands stripped). Structure refinement was done using Refmac^47^ and Buster^33^ and model building was done in Coot 0.9.6^32^.

### Fragment screening by X-ray crystallography

#### Screening with the P3_1_21 system

Crystal plates were set up as described above three days prior to the fragment screen. Crystal soaking and harvesting were done at the XChem facility, Diamond Light Source. Sub-wells were inspected to identify crystal drops that were suitable for fragment soaking, i.e. isolated cuboid crystals of >150 μm.

The DSi-Poised library (768 fragment, Enamine)^29^, the EUbOPEN DSiP extension library (108 fragments) and the FragLite library (31 fragments)^30^ were screened against TRF1_TRFH_ crystals. Compounds at 500 mM in DMSO stocks were dispensed into selected crystal drops as singletons using an Echo 550 liquid handler to give a final concentration of 50 mM (10% DMSO final concentration). After incubation for 1 h at 20 °C, 180 nL glycerol (40% stock, final drop concentration 28.8 – 32.5%) were added to each drop immediately prior to crystal harvesting and flash-freezing in liquid nitrogen.

Data were collected on the I04-1 beamline at Diamond Light Source. Data were processed using the Xia2-Dials pipeline^43,44^ and were then loaded into XChemExplorer (XCE)^48^ for data analysis. The XChem data processing pipeline was followed as described previously^27^. The refined apo-TRF1_TRFH_ structure was used as a reference structure to build a ground-state model that was then used for PanDDA analysis^31^. PanDDA Inspect was used to sequentially assess each event in Coot^32^, and ligands were added to the TRF1_TRFH_ model if density could be confidently assigned in real space. Datasets with ligands from the XCE analysis were then taken and refined manually using Refmac^47^ and Buster^33^ to assess ligand density in conventional 2F_o_-F_c_ and F_o_-F_c_ maps. Ligand restraints were generated with Grade2^49^.

#### Screening with the P4_1_2_1_2 system

Crystals in the P4_1_2_1_2 space group were grown as described above. Crystals in a total of 1075 drops were soaked with compounds from both the DSi Poised fragment library (768 fragments)^29^ and the “Probing fragment all” library (238) at the XChem facility, Diamond Light Source. Crystal drops were soaked using an Echo 550 liquid handler to give a final compound concentration of 50 mM and a final DMSO concentration of 10%. After 1 h, crystals were cryoprotected by addition of 10% ethylene glycol to the drops before crystal harvesting and flash-freezing in liquid nitrogen. Data were collected on the I04-1 beamline and analysed on XCE using the same method as for the P3_1_21 crystals. A summary of data collection and refinement statistics for datasets of fragment complexes is shown in Supplementary Table 3.

#### Fragment solubility assay

Aqueous solubility samples were prepared by adding 3.6 μL compound (50 mM stock in 100% DMSO-d6) to 176.4 μL NMR buffer in an Eppendorf tube. The samples were incubated overnight and transferred to a 3-mm NMR tube (Bruker, Part No. Z112272). 1H NMR spectra were recorded, with the DMSO and water signals dampened. 1 mM caffeine (Sigma, C1778 in PBS (pH 7.4) with 1% DMSO-d6) was used as a standard to quantify the ligand signals and determine the concentration of samples.

NMR data was collected on a Bruker Avance Neo 600-MHz spectrometer equipped with a 5 mm TCI-CryoProbe. The ^1^H spectrum was referenced to the internal deuterated solvent. The operating frequency for ^1^H was 600 MHz. All NMR data were acquired at the temperature of 298 K. All data were acquired and processed using Bruker Topspin 4.0. The quantitative ^1^H-NMR spectrum was acquired using a Bruker standard 1D lc1pngppsf2 pulse sequence with 32 scans. The sweep width was 6.2 ppm with O1P set to 8.8 ppm, and the FID contained 16k time-domain data points. Relaxation delay was set to 20 sec. Water signal was suppressed.

#### Fragment screening using the CPMG NMR assay

The fragment screen by LO-NMR was carried out using conditions described previously^50^, with some modifications. Fragments were dispensed in triplicate into a 384-well plate using an Echo 550 liquid handler to give a final ligand concentration of 250 μM (total sample volume of 40 μL). 40 μL of TRF1_TRFH_ protein (in NMR buffer; 20 μM final protein concentration in the assay) were added to the ‘protein’ samples; 40 μL of TRF1_TRFH_ protein and TIN2_TBM_ peptide (21-mer, Ac-RHFNLAPLGRRRVQSQWASTR-CONH_2_; 40 μM final concentration in the assay) were both added to the ‘competitor’ samples; 40 μL NMR buffer were added to the ‘compound-only’ samples. Solutions were transferred to 1.7-mm NMR tubes using a Bruker SamplePro Tube robotics system. ^1^H NMR spectra were recorded on a Bruker AVIII 500 MHz spectrometer at 298 K, equipped with a 1.7 mm TXI probe, with samples in 1.7-mm SampleJET NMR tubes (Bruker). DMSO and water signals were dampened. A relaxation spin filter was applied at 400 ms. Data were processed using Bruker Topspin 4.0. Lines were broadened with LB = 3.0, and the baseline was corrected between 6.0–10.0 ppm.

The data were then analysed using the MNova Screen software. Only peaks between 5.5 and 9.5 ppm were considered. Peaks with a height of <5% maximum peak height within the region of interest were considered noise, and the minimum matched peak level was set at >51%. The relative peak intensity change (I) was calculated for all peaks in the 5.5–9.5 ppm region, for each compound, as described previously^50^. The average integral for all peaks between 6.0-10.0 ppm was calculated, and the differences between compound-only, compound-plus-protein, and compound-plus-protein-plus-competitor samples were compared. The net recovery was calculated by subtracting the reduction in peak integral of the compound-plus-protein samples from the compound-plus-protein-plus-competitor samples.

Fragments that showed both a peak reduction of at least 35% in the presence of TRF1_TRFH_ and a net percentage recovery of peak intensity by at least 20% in the presence of both TRF1_TRFH_ and TIN2_TBM_ were classed as specific hits by MNova. For hit validation, relaxation-edited LO-NMR experiments were repeated as described above on single compounds instead of fragment cocktails.

#### R2KD assay

The R2KD assay was carried out as described by Liu *et al*^37^. A Bruker SamplePro liquid handler was used to prepare 10 samples: 8 samples containing an increasing ligand concentration from 50-500 μM whilst maintaining a constant TRF1_TRFH_ concentration (2 μM final); and two samples containing ligand only at 200 μM and 400 μM. All ten samples were prepared from stock solutions of ligand, DMSO, protein and buffer alone using the Bruker SamplePro liquid handler prior to transfer to 3-mm NMR tubes.

NMR data were collected at 298 K on a Bruker Avance Neo 600-MHz spectrometer using the Bruker Topspin 4.0 software. T_2_ relaxation experiments were performed using the standard Bruker CPMG pulse program with Excitation Sculpting incorporated to suppress water signal. The spin-echo period (delay-180°-delay) was set to 1 ms (d20 = 500 μs), and the relaxation delay (d1) was set to 10 s. The sudo-2D experiment contained nine slices with the spin echo period repeated at the following times: 4, 20, 100, 200, 300, 500, 700, 1000, and 2000 ms. T_2_ relaxation data was processed using the MestReNova 14.1 data analysis module to obtain integrals of individual ^1^H-NMR signals, which were used to calculate R_2_ with the equation I=I_0_e^(-tR2)^. GraphPad Prism 9.0.0 was used to fit R_2_ values of individual compound peaks to obtain an overall (shared) K_D_ value from all peaks of the compound.

#### Fluorescence polarisation (FP) assay

A TIN2_TBM_ peptide was obtained from Pepceuticals with a 5(6)-carboxyfluorescein (FAM) fluorophore added to the N-terminus for fluorescence studies. To determine the assay window for the FP assay, the TIN2-FAM probe was prepared as a 16-μM stock in aqueous buffer and titrated from 0-400 nM against either a constant concentration of TRF1_TRFH_ (0.5 µM in FP buffer) or FP buffer alone (25 mM HEPES-NaOH pH 7.5, 100 mM NaCl, 1 mM TCEP, 0.05% w/v CHAPS). Samples were prepared to a final volume of 10 μL in Packard Proxiplate 384 F plus plates; plates were incubated in darkness at room temperature for 1 h prior to measuring fluorescence intensities and calculating FP values, using a BMG Labtech PHERAstar FSX plate reader with an FP 485 520 520 optic module. A settling time of 0.3 s was used, and the number of flashes was set to 200 for FP measurements. For each plate, a target mP of 35 was set and the gain adjusted on the TIN2-FAM probe control well minus protein. An optimal assay window of 40 mP was observed at a probe concentration of 25 nM (Supplementary Fig. 4a).

To determine the K_D_ of the TRF1_TRFH_:TIN2-FAM interaction, an 11-point serial dilution of TRF1_TRFH_ was made from 20-0.02 μM, with each well backfilled to 5 μL with FP buffer. Each well was then made up to a final volume of 10 μL with 2x TIN2-FAM (50 nM) in FP buffer to give a final concentration of 25 nM. The same sample volume and measurement settings were used as described above.

For the FP competition assay, the unlabelled compounds (from 50-mM DMSO stocks) were first dispensed in duplicate into wells of a Proxiplate 384 F Plus plate using an Echo 550 acoustic liquid handler to give a 13-point concentration range from 0.2-500 μM. Wells were then backfilled to 250 nL with DMSO before the addition of 5 μL of 2X TRF1_TRFH_ (0.6 μM final concentration) and 4.75 μL 2x TIN2-FAM (25 nM final concentration). Plates were spun and sealed prior to incubation for 1 h in darkness. FP measurements were then carried out as described above.

#### Mass Photometry

Mass Photometry (MP) experiments were performed using a Refeyn OneMP instrument. Microscope coverslip and silicone gaskets (cut to 3 x 3 wells each) were cleaned sequentially with ultrafiltered distilled (UF) water, isopropanol and UF water before being dried with pressurised air. The cleaned coverslip and gasket were assembled on the microscope.

Contrast-to-mass calibration was generated using a native protein marker (NativeMark unstained protein standard, Thermo Scientific) with approximately 350-fold dilution in the MP buffer (50 mM HEPES-NaOH pH 8.0, 300 mM NaCl, filtered through a 0.2-μm filter). Standards of sizes 66, 146, 480 and 1048 kDa were used to fit the calibration curve with an R2 value of 99% and an error of 7.2%.

Experiments were set up using SEC-purified shelterin at a final concentration of 1.25 μM and incubated with increasing concentrations of compound **40**, in the range 100, 150, 200, 250 and 300 μM, and incubated for 1 h at room temperature. Controls were set up using comparable reaction conditions with shelterin and 300 μM of the inactive compound **60** or TRF1 only (without ligand). To prevent shelterin from breaking down into smaller subunits and to keep the complex intact during MP measurements, the samples were crosslinked after co-incubation with compounds, using 0.1% glutaraldehyde for 25 min at room temperature and quenched with Tris-HCl pH 8.0 for 5 min (Supplementary Fig. 8). Samples were rapidly diluted to a final protein concentration of 50 nM in a 15-μL buffer droplet in the well.

Movies of 60 s duration and 6000 frames were collected using Refeyn AcquireMP software (v2024) in a regular-sized (10.8 μm × 2.9 μm) field of view. Data were processed and analysed using Refeyn DiscoverMP software (v2024 R2). Gaussian fits for the mass distribution histograms, mass measurements and percentage abundance (of peak counts to total counts) were determined for each peak observed.

#### *In silico* fragment docking

Computational work was carried out using the MOE software package (Chemical Computing Group)^38^. Combinatorial libraries were generated using the QuaSAR application with 4-hydroxynaphthalene-1-sulfonic acid as a template and a series of commercially available benzyl bromide analogues as possible modifications of the benzyl group of compound **32**. The generated compound coordinates were then rebuilt in 3D, and the protonation state of each compound was set to the dominant state at pH 7.4.

The refined TRF1_TRFH_ model with compound **32** bound was chosen as a model to dock against. The naphthalene-1-sulfonic acid core was selected as a template for docking, and the receptor mobility was set to rigid. The computationally generated compounds were then docked into the target site and ranked according to the docking score. The top-scoring compounds were then synthesised as described in the Chemistry supplementary information.

## Supporting information

Chemistry Supplementary Information

## Acknowledgements

We thank Lauren Knightley, Beth Jago, Meirion Richards, and Amin Mirza of the Structural Chemistry team for their expertise and assistance, Swen Hoelder for his support, Stephen Hearnshaw for assistance with crystal optimisation, and Chris Richardson for crystallography computing support. We thank the staff of the Diamond Light Source (Didcot, UK), particularly the staff of the XChem facility and beamline I04-1, for their support with data collection. This work was funded by The Institute of Cancer Research (ICR), including an ICR PhD Studentship supporting G.C., the Wellcome Trust through a Wellcome Trust Investigator Award (214311/Z/18/Z) to S.G., and the Lister Institute of Preventive Medicine through a Lister Institute Research Prize Fellowship to S.G.

## Data availability

Atomic coordinates and structure factors for the crystal structures of the TRF1_TRFH_ domain in complex with the compounds discussed herein have been deposited in the Protein Data Bank (www.wwpdb.org) and can be accessed using the PDB accession numbers provided in Supplementary Table 3. The authors will release the atomic coordinates and experimental data upon article publication. Other data associated with this study are available upon request to the corresponding author.

## Author contributions

S.G. and I.C. conceived the project. S.G, I.C, Y.-V.L.-B. and J.C. supervised the project. G.C. and O.I. expressed and purified TRF1_TRFH_. G.C. and Y.-V.L.-B. optimised crystallisation and carried out crystallographic fragment screening, supervised by R.L.M.V.-M. G.C. and M.L. performed NMR assays. G.C. performed fluorescence polarisation. O.I. expressed and purified shelterin and full-length TRF1 and performed mass photometry. G.C. carried out *in silico* fragment docking. J.C. and I.C. designed fragment analogues. E.S. and J.C. synthesised compounds. G.C. wrote the manuscript with support from S.G. and I.C. All authors reviewed and approved the manuscript.

## Additional information

**Supplementary Information** comprising supplementary tables and figures

**Chemistry Supplementary Information** comprising compound sources, synthesis and validation details

