## Supplementary Information for "Discovery of first-in-class inhibitors of the TRF1-TIN2 protein-protein interaction by fragment screening"

**Supplementary Table 1: Analysis of commercially available analogues of XChem screen hit 1.**

| Compound no. | Structure | Aq. Sol. (μM) | CPMG reduction (%)[a] | CPMG recovery (%)[b] | Crystal structure |
|---|---|---|---|---|---|
| **20** | 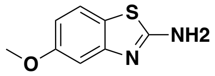 | 580 | 25 | 0 | Yes |
| **21** | 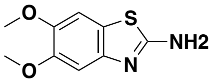 | 190 | 14 | 12 | No |
| **22** | 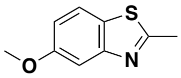 | 790 | 18 | 9 | Yes |
| **23** | 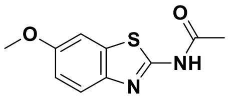 | 150 | 32 | 10 | No |
| **24** | 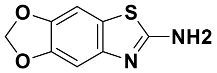 | 400 | 28 | 11 | No |

[a]The reduction in CPMG peak intensity upon addition of protein (n = 1)
[b]The net recovery of peak intensity upon addition of the $TIN2_{TBM}$ peptide to the compound-plus-protein sample (n = 1)

### Supplementary Table 2: Analysis of fragment hits from the crystallographic screens carried out against $TRF1_{TRFH}$.

**P3 XChem screen**

| Number: | Dataset no. | Structure | Crystal structure | Resolution (Å) | Buster ligand occupancy (%)* | Solubility (µM) | CPMG reduction (%) |
|---|---|---|---|---|---|---|---|
| **1** | TRF1-x0077 | 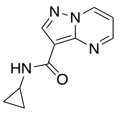 | 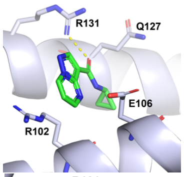  | 2.23 | 76 | 600 | 13 |
| **2** | TRF1-x0197 | 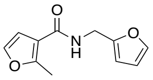 | 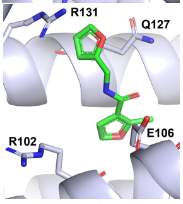  | 2.23 | 74 | 555 | 0 |
| **3** | TRF1-x0276 | 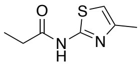 | 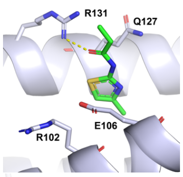  | 2.38 | 70 | 670 | 12 |
| **4** | TRF1-x0284 | 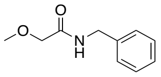 | 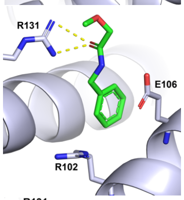  | 2.19 | 83 | 600 | 0 |
| **5** | TRF1-x0289 | 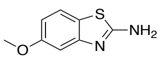 | 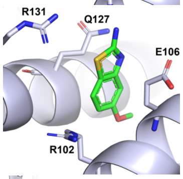  | 1.9 | 77 | 580 | 25 |
| **6** | TRF1-x0323 | 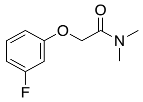 | 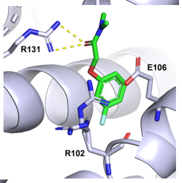  | 2.05 | 89 | 720 | 0 |
| **7** | TRF1-x0325 | 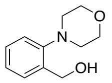 | 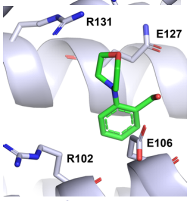  | 2.29 | 88 | 730 | 0 |
| **8** | TRF1-x0348 | 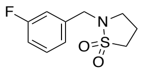 | 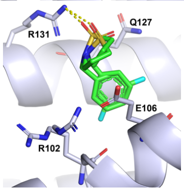  | 2.17 | 34/65 | 660 | 0 |
| **9** | TRF1-x0394 | 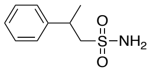 | 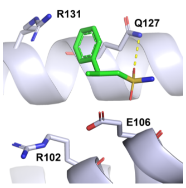  | 2.15 | 73 | 690 | 21 |

| | | | | | | | |
|---|---|---|---|---|---|---|---|
| **10** | TRF1-x0439 | 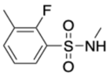 | 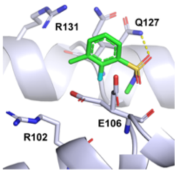 | 2.04 | 84 | 680 | 20 |
| **11** | TRF1-x0560 | 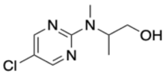 | 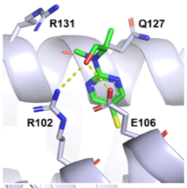 | 2.06 | 93 | 820 | 6 |
| **12** | TRF1-x0586 | 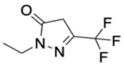 | 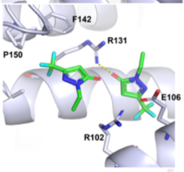 | 1.87 | 76/81 | 660 | 1 |
| **13** | TRF1-x0648 | 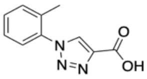 | 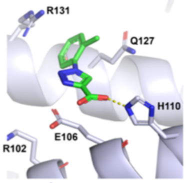 | 2.16 | 72 | 660 | 23 |
| **14** | TRF1-x0656 | 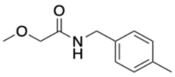 | 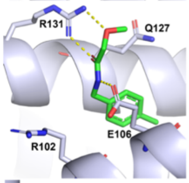 | 2.02 | 87 | 660 | 0 |
| **15** | TRF1-x0674 | 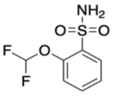 | 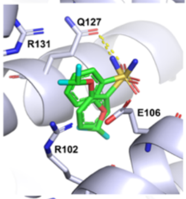 | 2.48 | 53/47 | 570 | 12 |
| **16** | TRF1-x0713 | 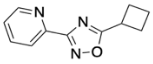 | 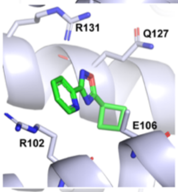 | 2.12 | 69 | 630 | 22 |
| **17** | TRF1-x0715 | 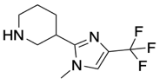 | 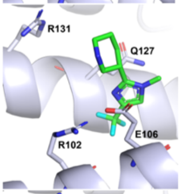 | 2.44 | 79 | - | 0 |
| **18** | TRF1-x0741 | 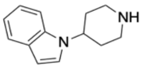 | 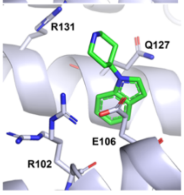 | 2.24 | 88 | - | 31 |
| **19** | TRF1-x0845 | 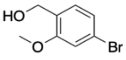 | 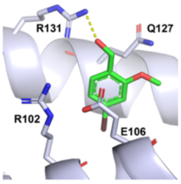 | 2.23 | 92 | - | 37 |

*Two values are shown in cases where two ligand conformations are observed in the binding site

**P4 XChem screen**

| Number: | Dataset no. | Structure | Crystal structure | Resolution (Å) | Buster ligand occupancy (%) | Solubility (μM) | CPMG reduction (%) |
|---|---|---|---|---|---|---|---|
| **1** | TRF1_2-x0021 | 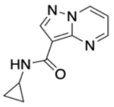 | 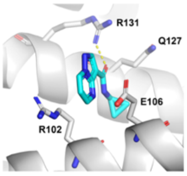 | 1.87 | 89 | 600 | 13 |
| **6** | TRF1_2-x0612 | 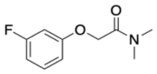 | 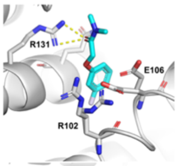 | 1.55 | 43 | 725 | 12 |
| **15** | TRF1_2-x0076 | 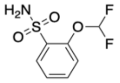 | 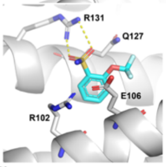 | 1.70 | 46 | 570 | 0 |
| **16** | TRF1_2-x0425 | 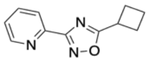 | 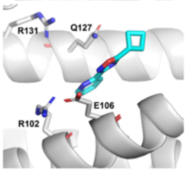 | 2.08 | 95 | 630 | 22 |
| **45** | TRF1_2-x0009 | 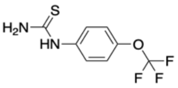 | 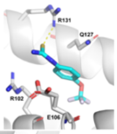 | 2.32 | 71 | 650 | |
| **46** | TRF1_2-x0025 | 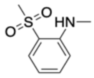 | 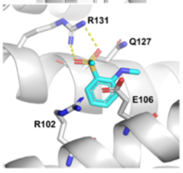 | 1.93 | 53 | 610 | 36 |
| **47** | TRF1_2-x0116 | 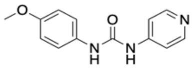 | 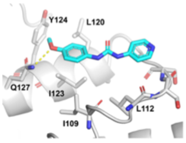 | 1.96 | 80 | >1000 | 56 |
| **48** | TRF1_2-x0195 | 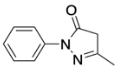 | 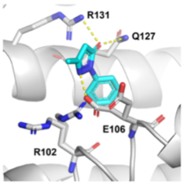 | 1.93 | 47 | 670 | 0 |
| **49** | TRF1_2-x0238 | 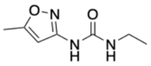 | 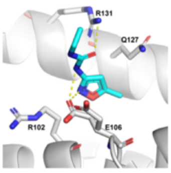 | 2.20 | 98 | 870 | 24 |

| | | | | | | | |
|---|---|---|---|---|---|---|---|
| **50** | TRF1_2-x0292 | 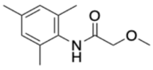 | 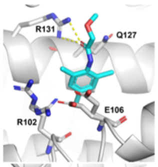  | 1.70 | 88 | 660 | 23 |
| **51** | TRF1_2-x0312 | 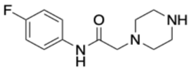 | 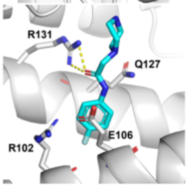  | 2.07 | 86 | 570 | 11 |
| **52** | TRF1_2-x0522 | 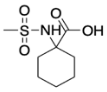 | 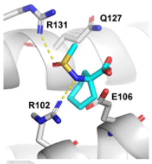  | 2.08 | 92 | 0 | - |
| **53** | TRF1_2-x0572 | 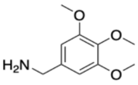 | 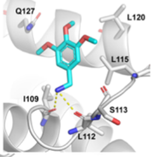  | 1.69 | 67 | 575 | 11 |
| **54** | TRF1_2-x0592 | 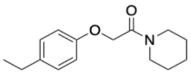 | 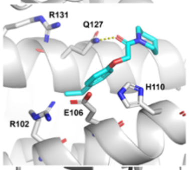  | 1.93 | 62 | 485 | 21 |
| **55** | TRF1_2-x0656 | 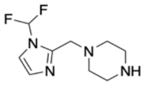 | 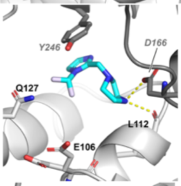  | 1.60 | 48 | 485 | 0 |
| **56** | TRF1_2-x0866 | 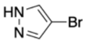 | 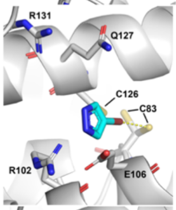  | 2.17 | 39 | | 14 |
| **57** | TRF1_2-x0869 | 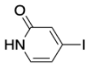 | 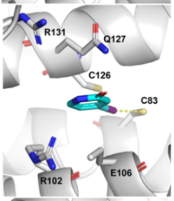  | 2.50 | 48 | >1000 | 25 |
| **58** | TRF1_2-x0874 | 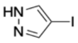 | 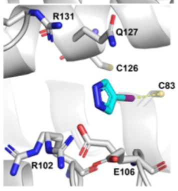  | 2.10 | 66 | - | 9 |
| **59** | TRF1_2-x1043 | 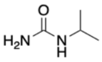 | 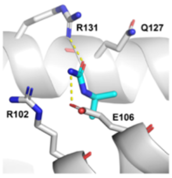  | 1.73 | 100 | - | - |

**Supplementary Table 3: X-ray crystallography data collection and refinement statistics.** Fragment hits from the first crystallographic fragment screen using the $P3_121$ $TRF1_{TRFH}$ crystal system are numbered **1 – 19.**Three fragment hits developed from the LO-NMR fragment screen and soaked into $TRF1_{TRFH}$ crystals in the $P4_12_12$ space group are numbered **27**, **32**, and **40**, respectively. Fragment hits from the second crystallographic screen using the $P4_12_12$ $TRF1_{TRFH}$ crystal system are numbered **45 – 59.**

| **Compound_ID** | **1** | **2** | **3** | **4** | **5** | **6** | **7** | **8** | **9** | **10** |
|---|---|---|---|---|---|---|---|---|---|---|
| **PDB_ID** | **9HF6** | **9HHK** | **9HF7** | **9HF8** | **9HF9** | **9HFA** | **9HFB** | **9HFC** | **9HFD** | **9HFE** |
| **Dataset_ID** | TRF1-x0077 | TRF1-x0197 | TRF1-x0276 | TRF1-x0284 | TRF1-x0289 | TRF1-x0323 | TRF1-x0325 | TRF1-x0348 | TRF1-x0394 | TRF1-x0439 |
| **Crystal** | | | | | | | | | | |
| Space group | P 31 2 1 | P 31 2 1 | P 31 2 1 | P 31 2 1 | P 31 2 1 | P 31 2 1 | P 31 2 1 | P 31 2 1 | P 31 2 1 | P 31 2 1 |
| Unit cell dimensions (a/b/c in Å) | 84.71, 84.71, 91.44 | 85.09, 85.09, 91.45 | 85.01, 85.01, 91.29 | 85.3, 85.3, 91.55 | 85.61, 85.61, 90.83 | 85.54, 85.54, 91.35 | 85.25, 85.25, 91.19 | 84.69, 84.69, 91.51 | 85.37, 85.37, 91.64 | 85.37, 85.37, 91.26 |
| Unit cell angles (α/β/γ in °) | 90, 90, 120 | 90, 90, 120 | 90, 90, 120 | 90, 90, 120 | 90, 90, 120 | 90, 90, 120 | 90, 90, 120 | 90, 90, 120 | 90, 90, 120 | 90, 90, 120 |
| **Data collection and processing** | | | | | | | | | | |
| Beamline | i04-1 | i04-1 | i04-1 | i04-1 | i04-1 | i04-1 | i04-1 | i04-1 | i04-1 | i04-1 |
| Wavelength (Å) | 0.9179 | 0.9179 | 0.9179 | 0.9179 | 0.9179 | 0.9179 | 0.9179 | 0.9179 | 0.9179 | 0.9179 |
| Integration program | XDS | XDS | XDS | XDS | XDS | XDS | XDS | XDS | XDS | XDS |
| Reduction program | AIMLESS | AIMLESS | AIMLESS | AIMLESS | AIMLESS | AIMLESS | AIMLESS | AIMLESS | AIMLESS | AIMLESS |
| Resolution range | 42.35 - 2.23 | 73.69 - 2.23 | 57.3 - 2.23 | 57.49 - 2.19 | 45.39 (1.90) | 45.68 - 2.05 | 57.38 - 2.29 | 57.23 - 2.17 | 45.82 - 2.15 | 45.63 - 2.04 |
| Number of unique reflections [a] | 18969 (1729) | 19097 (1727) | 19057 (1734) | 20300 (1729) | 30774 (1941) | 24733 (1885) | 17716 (1714) | 20560 (1764) | 21466 (1815) | 24971 (1904) |
| Completeness [a] | 100 (100) | 100 (100) | 100 (100) | 100 (100) | 100 (100) | 100 (100) | 100 (100) | 100 (100) | 100 (100) | 100 (100) |
| Redundancy [a] | 9.7 (9.7) | 9.9 (9.9) | 9.4 (9.4) | 9.8 (9.3) | 9.8 (8.7) | 9.6 (9.9) | 9.6 (10.1) | 9.8 (9.4) | 9.7 (9.5) | 9.8 (10.1) |
| $R_{merge}$ (%) [a] | 0.069 (2.494) | 0.084 (2.608) | 0.147 (5.052) | 0.066 (2.636) | 0.071 (2.838) | 0.078 (2.500) | 0.095 (2.586) | 0.064 (2.405) | 0.102 (2.490) | 0.062 (2.595) |
| I/σ(I) [a] | 14.6 (0.7) | 11.2 (1.0) | 6.0 (0.3) | 14.0 (0.7) | 12.8 (0.6) | 13.3 (0.7) | 11.6 (1.0) | 14.1 (0.6) | 8.9 (0.7) | 16.1 (0.7) |
| $CC_{1/2}$ [a, b] | 1 (0.817) | 0.998 (0.816) | 0.998 (0.860) | 0.999 (0.789) | 0.999 (0.785) | 1.000 (0.767) | 0.999 (0.758) | 0.976 (0.918) | 0.998 (0.944) | 1.000 (0.652) |
| **Refinement** | | | | | | | | | | |
| Programme | BUSTER | BUSTER | BUSTER | BUSTER | BUSTER | BUSTER | BUSTER | BUSTER | BUSTER | BUSTER |
| $R_{work}$ (%) | 0.2291 | 0.2325 | 0.2342 | 0.2279 | 0.2131 | 0.2251 | 0.2292 | 0.2366 | 0.2293 | 0.2203 |
| $R_{free}$ (%) | 0.2483 | 0.2458 | 0.2614 | 0.255 | 0.2279 | 0.2456 | 0.2652 | 0.2432 | 0.2446 | 0.2471 |
| Number of residues | 193 | 202 | 198 | 197 | 198 | 192 | 193 | 202 | 196 | 194 |
| Number of water molecules | 57 | 43 | 53 | 42 | 124 | 90 | 57 | 50 | 64 | 90 |
| Average B-factor ($Å^2$) | 84.2 | 84.11 | 84.34 | 82.58 | 59.09 | 63.42 | 76.47 | 88.99 | 74.09 | 67.48 |
| Ramachandran favoured (%) | 99.46 | 98.47 | 98.94 | 99.47 | 98.95 | 98.92 | 100 | 98.98 | 99.46 | 99.46 |
| Ramachandran outliers (%) | 0 | 0 | 0 | 0 | 0 | 0 | 0 | 0 | 0 | 0 |
| RMSD bonds (Å) | 0.012 | 0.0111 | 0.0121 | 0.012 | 0.0135 | 0.0133 | 0.0116 | 0.0124 | 0.0173 | 0.0119 |
| RMSD angles (°) | 1.367 | 1.313 | 1.317 | 1.31 | 1.336 | 1.33 | 1.334 | 1.322 | 1.392 | 1.3 |

[a]Values between brackets are for the highest resolution shell.

| Compound_ID | 11 | 12 | 13 | 14 | 15 | 16 | 17 | 18 | 19 |
|---|---|---|---|---|---|---|---|---|---|
| **PDB_ID** | **9HFF** | **9HCL** | **9HLT** | **9HFG** | **9HLR** | **9HFH** | **9HFI** | **9HLP** | **9HLU** |
| **Dataset_ID** | TRF1-x0560 | TRF1-x0586 | TRF1-x0648 | TRF1-x0656 | TRF1-x0674 | TRF1-x0713 | TRF1-x0715 | TRF1-x0741 | TRF1-x0845 |
| **Crystal** | | | | | | | | | |
| Space group | P 31 2 1 | P 31 2 1 | P 31 2 1 | P 31 2 1 | P 31 2 1 | P 31 2 1 | P 31 2 1 | P 31 2 1 | P 31 2 1 |
| Unit cell dimensions (a/b/c in Å) | 86.46, 86.46, 91.79 | 85.92, 85.92, 91.11 | 85.61, 85.61, 91.03 | 85.79, 85.79, 91.42 | 85.02, 85.02, 91.62 | 84.91, 84.91, 91.28 | 84.87, 84.87, 92.11 | 85.2, 85.2, 91.1 | 85.82, 85.82, 91.16 |
| Unit cell angles (α/β/γ in °) | 90, 90, 120 | 90, 90, 120 | 90, 90, 120 | 90, 90, 120 | 90, 90, 120 | 90, 90, 120 | 90, 90, 120 | 90, 90, 120 | 90, 90, 120 |
| **Data collection and processing** | | | | | | | | | |
| Beamline | i04-1 | i04-1 | i04-1 | i04-1 | i04-1 | i04-1 | i04-1 | i04-1 | i04-1 |
| Wavelength (Å) | 0.9179 | 0.9179 | 0.9179 | 0.9179 | 0.9179 | 0.9179 | 0.9179 | 0.9179 | 0.9179 |
| Integration program | XDS | XDS | XDS | XDS | XDS | XDS | XDS | XDS | XDS |
| Reduction program | AIMLESS | AIMLESS | AIMLESS | AIMLESS | AIMLESS | AIMLESS | AIMLESS | AIMLESS | AIMLESS |
| Resolution range | 45.9 - 2.06 | 74.41 - 1.87 | 57.49 - 2.16 | 45.71 - 2.02 | 57.39 - 2.48 | 73.54 - 2.12 | 57.45 - 2.44 | 57.34 - 2.24 | 57.60 - 2.23 |
| Number of unique reflections [a] | 25010 (1906) | 32610 (2043) | 21166 (1799) | 26013 (1892) | 13996 (1552) | 22083 (1796) | 14711 (1612) | 18844 (1703) | 19399 (1763) |
| Completeness [a] | 100 (100) | 100 (100) | 100 (100) | 100 (100) | 100 (100) | 100 (100) | 100 (100) | 100 (100) | 100 (100) |
| Redundancy [a] | 9.6 (9.9) | 9.8 (8.3) | 9.9 (9.5) | 9.7 (10.1) | 9.6 (10.1) | 9.7 (9.8) | 9.9 (10.3) | 9.9 (10.1) | 9.7 (9.8) |
| $R_{merge}$ (%) [a] | 0.092 (1.901) | 0.050 (1.791) | 0.067 (1.920) | 0.076 (2.583) | 0.100 (1.643) | 0.073 (2.569) | 0.051 (2.370) | 0.060 (2.592) | 0.084 (1.889) |
| I/σ(I) [a] | 9.8 (0.8) | 19.7 (0.8) | 13.3 (0.8) | 13.6 (0.8) | 8.5 (0.7) | 12.4 (0.8) | 17.3 (0.7) | 14.7 (0.9) | 12.2 (0.7) |
| $CC_{1/2}$ [a, b] | 0.999 (0.794) | 1.000 (0.728) | 0.948 (0.963) | 1.000 (0.702) | 0.997 (0.855) | 1.000 (0.960) | 1.000 (0.695) | 0.999 (0.858) | 0.998 (0.878 |
| **Refinement** | | | | | | | | | |
| Programme | BUSTER | BUSTER | BUSTER | BUSTER | BUSTER | BUSTER | BUSTER | BUSTER | BUSTER |
| $R_{work}$ (%) | 0.2238 | 0.2044 | 0.237 | 0.2282 | 0.2347 | 0.2286 | 0.2434 | 0.2356 | 0.2425 |
| $R_{free}$ (%) | 0.2612 | 0.2203 | 0.2605 | 0.2525 | 0.2689 | 0.2628 | 0.2758 | 0.2551 | 0.2794 |
| Number of residues | 201 | 199 | 197 | 192 | 202 | 195 | 197 | 196 | 190 |
| Number of water molecules | 99 | 193 | 65 | 134 | 28 | 72 | 21 | 60 | 55 |
| Average B-factor (Å$^2$) | 68.44 | 54.24 | 79.67 | 62.15 | 101.15 | 74.79 | 108.81 | 90.98 | 79.37 |
| Ramachandran favoured (%) | 98.46 | 98.96 | 99.47 | 98.91 | 98.48 | 99.46 | 98.95 | 97.88 | 97.27 |
| Ramachandran outliers (%) | 0 | 0 | 0 | 0 | 0 | 0 | 0 | 0 | 0 |
| RMSD bonds (Å) | 0.0121 | 0.0126 | 0.0121 | 0.0132 | 0.0101 | 0.012 | 0.0125 | 0.0118 | 0.0119 |
| RMSD angles (°) | 1.344 | 1.337 | 1.3 | 1.32 | 1.231 | 1.313 | 1.345 | 1.345 | 1.372 |

[a]Values between brackets are for the highest resolution shell.

| Compound_ID | 27 | 32 | 40 | 45 | 1 | 46 | 15 | 47 | 48 | 49 | 50 |
|---|---|---|---|---|---|---|---|---|---|---|---|
| **PDB_ID** | **9HD3** | **9HF4** | **9HD2** | **9HCM** | **9HD9** | **9HCN** | **9HCP** | **9HCQ** | **9HCR** | **9HCS** | **9HCT** |
| **Dataset_ID** | - | - | - | TRF1_2-x0009 | TRF1_2-x0021 | TRF1_2-x0025 | TRF1_2-x0076 | TRF1_2-x0116 | TRF1_2-x0195 | TRF1_2-x0195 | TRF1_2-x0292 |
| **Crystal** | | | | | | | | | | | |
| Space group | P 41 21 2 | P 41 21 2 | P 41 21 2 | P 41 21 2 | P 41 21 2 | P 41 21 2 | P 41 21 2 | P 41 21 2 | P 41 21 2 | P 41 21 2 | P 41 21 2 |
| Unit cell dimensions (a/b/c in Å) | 51.82, 51.82, 145.72 | 51.73, 51.73, 145.24 | 52.40 52.40 146.34 | 52.18, 52.18, 146.29 | 51.87, 51.87, 146.94 | 52.03, 52.03, 147.38 | 51.87, 51.87, 147.22 | 51.66, 51.66, 146.53 | 51.96, 51.96, 146.99 | 51.85, 51.85, 147.25 | 52.03, 52.03, 147.33 |
| Unit cell angles (α/β/γ in °) | 90, 90, 90 | 90, 90, 90 | 90, 90, 90 | 90, 90, 90 | 90, 90, 90 | 90, 90, 90 | 90, 90, 90 | 90, 90, 90 | 90, 90, 90 | 90, 90, 90 | 90, 90, 90 |
| **Data collection and processing** | | | | | | | | | | | |
| Beamline | i03 | i04 | i04-1 | i04-1 | i04-1 | i04-1 | i04-1 | i04-1 | i04-1 | i04-1 | i04-1 |
| Wavelength (Å) | 0.9763 | 0.9537 | 0.9212 | 0.9212 | 0.9212 | 0.9212 | 0.9212 | 0.9212 | 0.9212 | 0.9212 | 0.9212 |
| Integration program | XDS | XDS | XDS | XDS | XDS | XDS | XDS | XDS | XDS | XDS | XDS |
| Reduction program | AIMLESS | AIMLESS | AIMLESS | AIMLESS | AIMLESS | AIMLESS | AIMLESS | AIMLESS | AIMLESS | AIMLESS | AIMLESS |
| Resolution range | 42.23 - 2.19 | 48.73 - 2.17 | 49.33 - 2.07 | 49.15 - 2.32 | 51.87 - 1.87 | 49.13 - 1.93 | 49.07 - 1.70 | 51.66 - 1.96 | 48.99 - 1.93 | 49.08 - 2.20 | 49.11 - 1.70 |
| Number of unique reflections [a] | 10947 (916) | 11161 (929) | 13216 (1003) | 9421 (885) | 17475 (1103) | 16077 (1034) | 23100 (1211) | 15090 (1029) | 15976 (1023) | 10903 (911) | 23251 (1210) |
| Completeness [a] | 100 (100) | 100 (100) | 100 (100) | 100 (100) | 100 (100) | 100 (100) | 100 (100) | 100 (100) | 100 (100) | 99.8 (99.9) | 100 (100) |
| Redundancy [a] | 25.4 (25.0) | 24.2 (25.5) | 14.1 (14.7) | 13.9 (15.0) | 13.5 (11.0) | 13.8 (12.0) | 12.9 (9.2) | 14.1 (13.6) | 13.1 (11.3) | 13.9 (13.6) | 13.1 (9.4) |
| $R_{merge}$ (%) [a] | 0.094 (2.820) | 0.072 (2.602) | 0.121 (2.255) | 0.167 (2.612) | 0.183 (1.576) | 0.145 (2.949) | 0.085 (1.900) | 0.226 (2.854) | 0.205 (2.789) | 0.301 (2.427) | 0.064 (2.443) |
| I/σ(I) [a] | 19.0 (1.6) | 20.1 (1.4) | 8.8 (0.8) | 10.5 (3.2) | 7.6 (0.9) | 11.2 (1.8) | 13.4 (0.9) | 8.0 (2.4) | 8.8 (2.5) | 13.0 (5.7) | 15.9 (0.9) |
| $CC_{1/2}$ [a] | 0.999 (0.833) | 1.000 (0.927) | 0.997 (0.855) | 0.992 (0.977) | 0.986 (0.839) | 0.996 (0.845) | 0.997 (0.713) | 0.987 (0.945) | 0.969 (0.853) | 0.961 (0.446) | 0.999 (0.708) |
| **Refinement** | | | | | | | | | | | |
| Program | BUSTER | BUSTER | BUSTER | BUSTER | BUSTER | BUSTER | BUSTER | BUSTER | BUSTER | BUSTER | BUSTER |
| $R_{work}$ (%) | 23.95 | 23.19 | 22.29 | 23.01 | 22.82 | 20.52 | 20.04 | 20.24 | 19.56 | 20.51 | 20.29 |
| $R_{free}$ (%) | 27.38 | 25.75 | 26.78 | 31.2 | 26.99 | 22.74 | 24.44 | 24.53 | 21.68 | 24.23 | 24.04 |
| Number of residues | 206 | 204 | 205 | 205 | 204 | 205 | 205 | 204 | 204 | 204 | 204 |
| Number of water molecules | 26 | 22 | 48 | 48 | 79 | 99 | 129 | 100 | 134 | 100 | 117 |
| Average B-factor (Å$^2$) | 67.4 | 75.88 | 66.3 | 63.8 | 40.07 | 43.31 | 39.3 | 41.64 | 39.7 | 34.18 | 46.32 |
| Ramachandran favoured (%) | 99.5 | 96.98 | 99.5 | 99.5 | 97.5 | 98 | 98 | 98.49 | 97.99 | 97.5 | 97.5 |
| Ramachandran outliers (%) | 0 | 0 | 0 | 0 | 0 | 0 | 0 | 0 | 0 | 0 | 0 |
| RMSD bonds (Å) | 0.013 | 0.012 | 0.013 | 0.011 | 0.0131 | 0.0124 | 0.0135 | 0.013 | 0.0126 | 0.0119 | 0.0135 |
| RMSD angles (°) | 1.359 | 1.407 | 1.351 | 1.305 | 1.259 | 1.281 | 1.297 | 1.302 | 1.286 | 1.317 | 1.334 |

[a]Values in parentheses are for the highest resolution shell.

| Compound_ID | 51 | 16 | 52 | 53 | 54 | 6 | 55 | 56 | 57 | 58 | 59 |
|---|---|---|---|---|---|---|---|---|---|---|---|
| **PDB_ID** | **9HCU** | **9HCV** | **9HCW** | **9HCX** | **9HCY** | **9HCZ** | **9HD0** | **-** | **9HLQ** | **9HDA** | **9HD1** |
| **Dataset_ID** | TRF1_2-x0312 | TRF1_2-x0425 | TRF1_2-x0522 | TRF1_2-x0572 | TRF1_2-x0592 | TRF1_2-x0612 | TRF1_2-x0656 | TRF1_2-x0866 | TRF1_2-x0869 | TRF1_2-x0874 | TRF1_2-x1043 |
| **Crystal** | | | | | | | | | | | |
| Space group | P 41 21 2 | P 41 21 2 | P 41 21 2 | P 41 21 2 | P 41 21 2 | P 41 21 2 | P 41 21 2 | | P 41 21 2 | P 41 21 2 | P 41 21 2 |
| Unit cell dimensions (a/b/c in Å) | 52.03, 52.03, 146.94 | 52.44, 52.44, 146.11 | 52.35, 52.35, 147.07 | 51.66, 51.66, 146.62 | 51.92, 51.92, 147.23 | 51.88, 51.88, 147.22 | 51.77, 51.77, 146.79 | 52.07, 52.07, 146.42 | 51.97, 51.97, 145.88 | 52.27, 52.27, 146.05 | 51.65, 51.65, 147.18 |
| Unit cell angles (α/β/γ in °) | 90, 90, 90 | 90, 90, 90 | 90, 90, 90 | 90, 90, 90 | 90, 90, 90 | 90, 90, 90 | 90, 90, 90 | 90, 90, 90 | 90, 90, 90 | 90, 90, 90 | 90, 90, 90 |
| **Data collection and processing** | | | | | | | | | | | |
| Beamline | i04-1 | i04-1 | i04-1 | i04-1 | i04-1 | i04-1 | i04-1 | i04-1 | i04-1 | i04-1 | i04-1 |
| Wavelength (Å) | 0.9212 | 0.9212 | 0.9212 | 0.9212 | 0.9212 | 0.9212 | 0.9212 | 0.9212 | 0.9212 | 0.9212 | 0.9212 |
| Integration program | XDS | XDS | XDS | XDS | XDS | XDS | XDS | XDS | XDS | XDS | XDS |
| Reduction program | AIMLESS | AIMLESS | AIMLESS | AIMLESS | AIMLESS | AIMLESS | AIMLESS | AIMLESS | AIMLESS | AIMLESS | AIMLESS |
| Resolution range | 49.05 - 2.07 | 52.44 - 2.08 | 49.32 - 2.08 | 48.87 - 1.69 | 49.08 - 1.93 | 51.88 - 1.55 | 51.77 - 1.60 | 49.06 - 2.17 | 51.97 - 2.50 | 52.27 - 2.10 | 51.65 - 1.73 |
| Number of unique reflections [a] | 13101 (979) | 13036 (968) | 13072 (977) | 22736 (1085) | 15989 (1021) | 30074 (1369) | 27393 (1344) | 10873 (527) | 7528 (817) | 12580 (989) | 21763 (1144) |
| Completeness [a] | 100 (99.9) | 100 (100) | 100 (100) | 98.5 (94) | 100 (100) | 99.5 (94.2) | 100 (100) | 95.4 (100) | 100 (100) | 100 (100) | 100 (100) |
| Redundancy [a] | 12.6 (12.1) | 14.2 (14.8) | 14.2 (14.8) | 12.1 (7.7) | 13.7 (12.6) | 11.5 (4.1) | 12.0 (5.3) | 9.9 (10) | 13.3 (13.4) | 14.2 (14.7) | 13.3 (9.8) |
| $R_{merge}$ (%) [a] | 0.281 (2.779) | 0.111 (2.938) | 0.080 (2.411) | 0.200 (2.134) | 0.142 (2.409) | 0.068 (1.202) | 0.074 (1.298) | 0.248 (6.329) | 0.202 (3.003) | 0.106 (2.796) | 0.109 (2.729) |
| I/σ(I) [a] | 5.8 (1.5) | 10.9 (1.3) | 14.2 (1.4) | 6.5 (0.9) | 9.5 (1.1) | 15.1 (0.9) | 16.0 (1.4) | 8.8 (0.5) | 6.8 (1.2) | 13.9 (1.8) | 13.9 (1.4) |
| $CC_{1/2}$ [a, b] | 0.988 (0.837) | 0.998 (0.867) | 0.994 (0.777) | 0.989 (0.621) | 0.989 (0.685) | 0.997 (0.773) | 0.998 (0.701) | 1.0 (0.4) | 0.989 (0.645) | 0.995 (0.810) | 0.998 (0.491) |
| **Refinement** | | | | | | | | | | | |
| Program | BUSTER | BUSTER | BUSTER | BUSTER | BUSTER | BUSTER | BUSTER | BUSTER | BUSTER | BUSTER | BUSTER |
| $R_{work}$ (%) | 22.62 | 22.9 | 24.21 | 19.52 | 20.1 | 19.85 | 19.76 | 23.22 | 26.26 | 23.17 | 19.37 |
| $R_{free}$ (%) | 25.99 | 26.84 | 25.53 | 24.71 | 22.28 | 21.43 | 22.44 | 28.91 | 29.74 | 28.29 | 22.66 |
| Number of residues | 204 | 204 | 204 | 204 | 204 | 204 | 206 | 204 | 204 | 206 | 204 |
| Number of water molecules | 78 | 49 | 34 | 133 | 126 | 153 | 127 | 58 | 13 | 42 | 143 |
| Average B-factor (Å$^2$) | 40.77 | 66.5 | 68.35 | 30.69 | 38.26 | 36.62 | 33.7 | 47.93 | 89.62 | 69.43 | 36.15 |
| Ramachandran favoured (%) | 97.99 | 99.5 | 97.49 | 98.49 | 97.99 | 97.99 | 98.51 | 97.99 | 97.41 | 97.03 | 99 |
| Ramachandran outliers (%) | 0 | 0 | 0 | 0 | 0 | 0 | 0 | 0 | 0 | 0.5 | 0 |
| RMSD bonds (Å) | 0.0122 | 0.0121 | 0.0119 | 0.0135 | 0.0124 | 0.0142 | 0.0138 | 0.012 | 0.0113 | 0.0122 | 0.0135 |
| RMSD angles (°) | 1.335 | 1.31 | 1.354 | 1.362 | 1.316 | 1.362 | 1.445 | 1.421 | 1.345 | 1.384 | 1.324 |

[a]Values in parentheses are for the highest resolution shell.

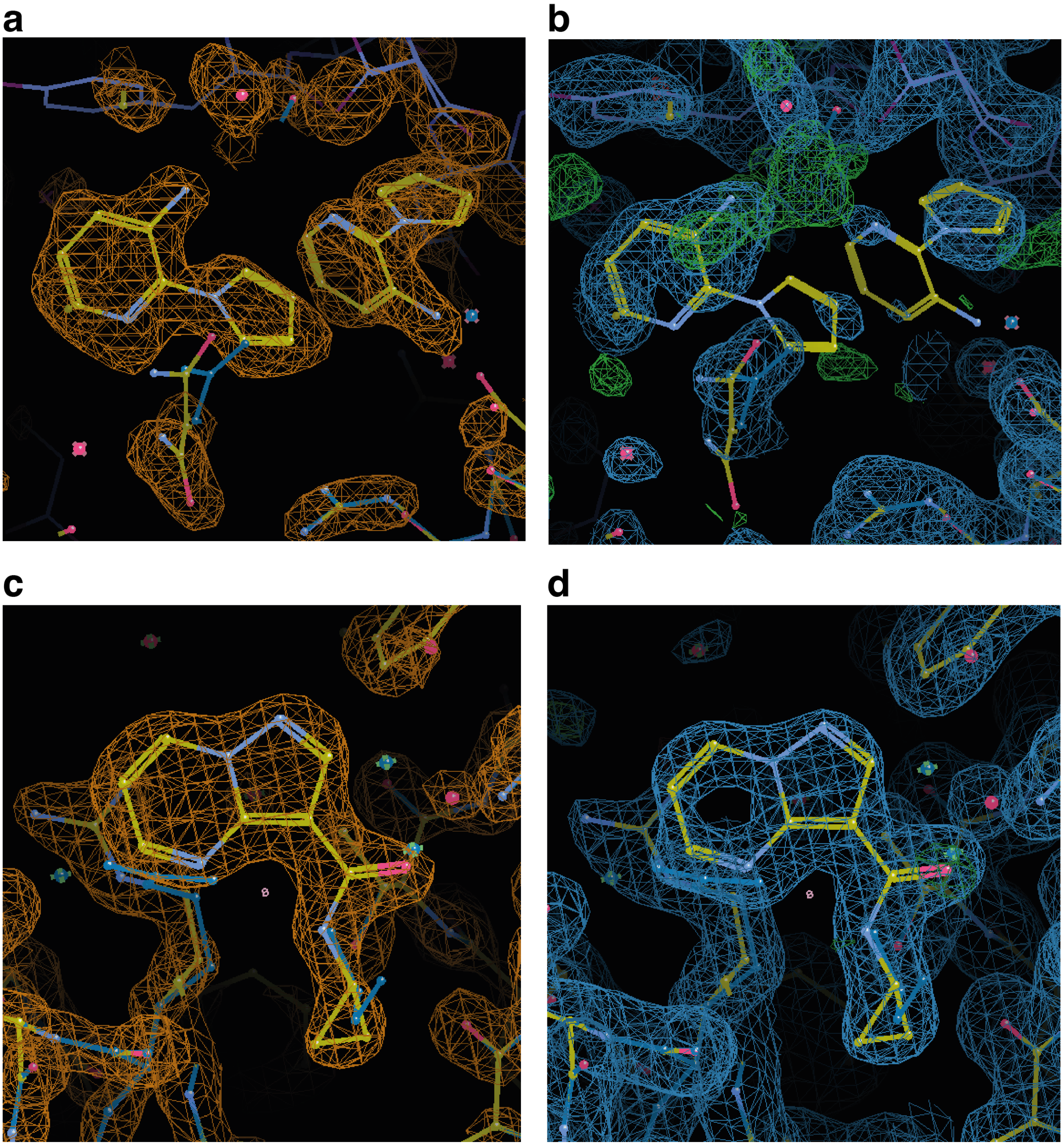

**Supplementary Figure 1: Examples of PANDDA maps versus conventional 2mF$_o$-DF$_c$ electron density maps. (a)** PANDDA map of dataset TRF1_2-x0455 showing two fragments bound to the TRF1$_{TRFH}$ crystal compared to **(b)** 2mF$_o$-DF$_c$ map after refinement with Buster. Given the poorly defined density, this hit was not taken forward. **(c)** PANDDA map of dataset TRF1_2-x0021 showing compound **1** bound to TRF1$_{TRFH}$ compared to **(d)** 2mF$_o$-DF$_c$ maps after refinement with Buster. All PANDDA maps are contoured at 2.40 RMSD level, and 2mF$_o$-DF$_c$ maps are contoured at 1.00 RMSD level.

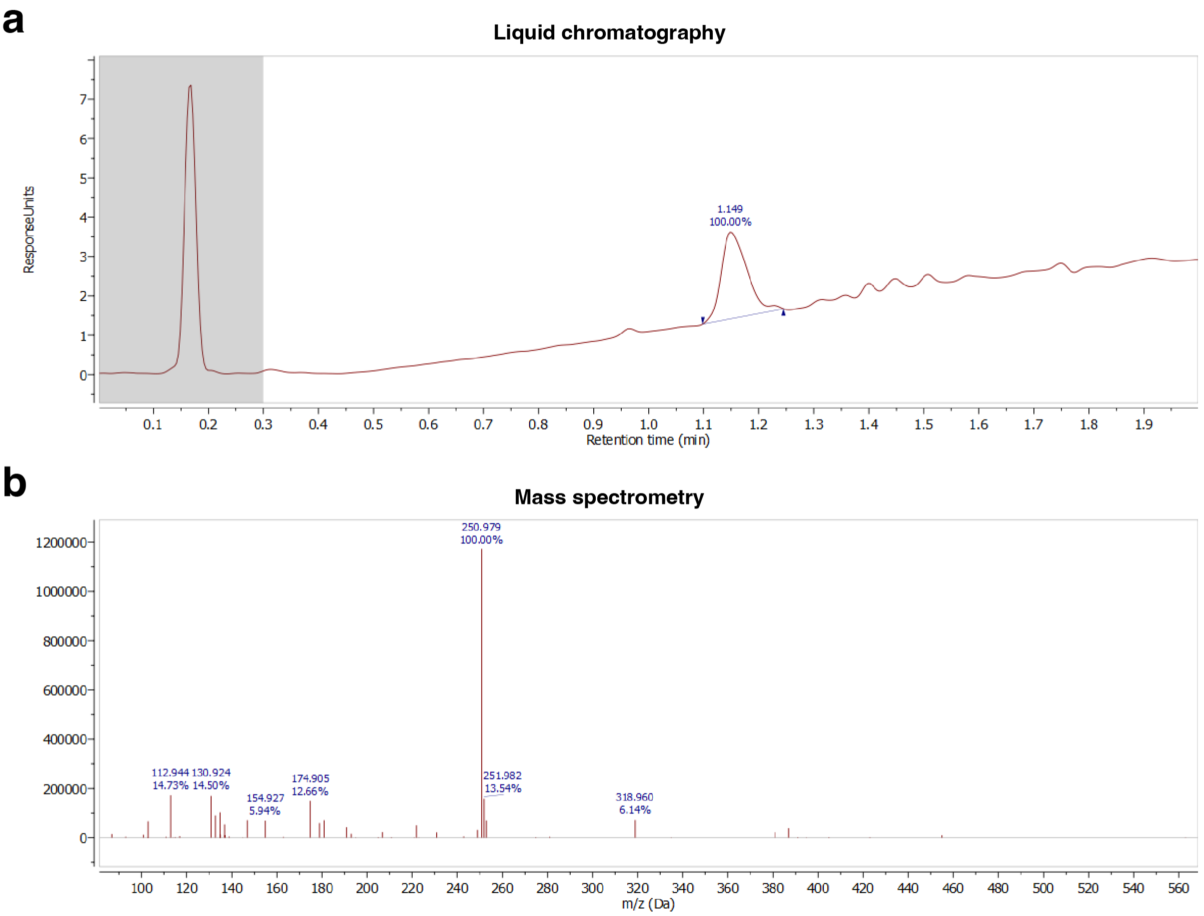

**Supplementary Figure 2: Negative mode LC-MS analysis of compound 25.** The LC-UV trace suggested a major compound eluting at 1.149 min **(a)** with an m/z value of 250.98 **(b)**. This value aligned with the molecular weight of compound **27** (251 g $mol^{-1}$), confirming that compound **25** in the original screening DMSO stock had degraded into compound **27**.

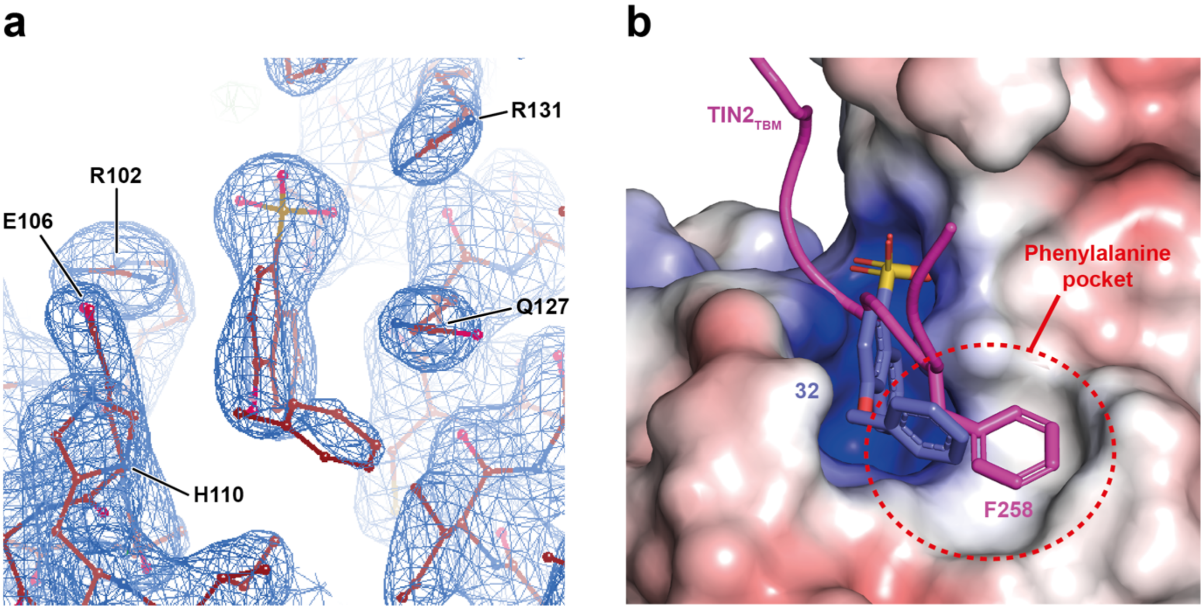

**Supplementary Figure 3: Binding mode of compound 32. (a)** Sigma-A weighted 2mF$_o$-DF$_c$ electron density map of compound **32** bound to TRF1$_{TRFH}$ (from P4$_1$2$_1$2 crystals) contoured at a level of 1.0. **(b)** Phenylalanine of the TIN2$_{TBM}$ peptide (L258) from the TRF1:TIN2 co-crystal structure (PDB: 3BQO) is shown in stick representation, coloured in magenta, superimposed onto the TRF1$_{TRFH}$-**32** structure, with compound **32** shown in stick representation in blue.

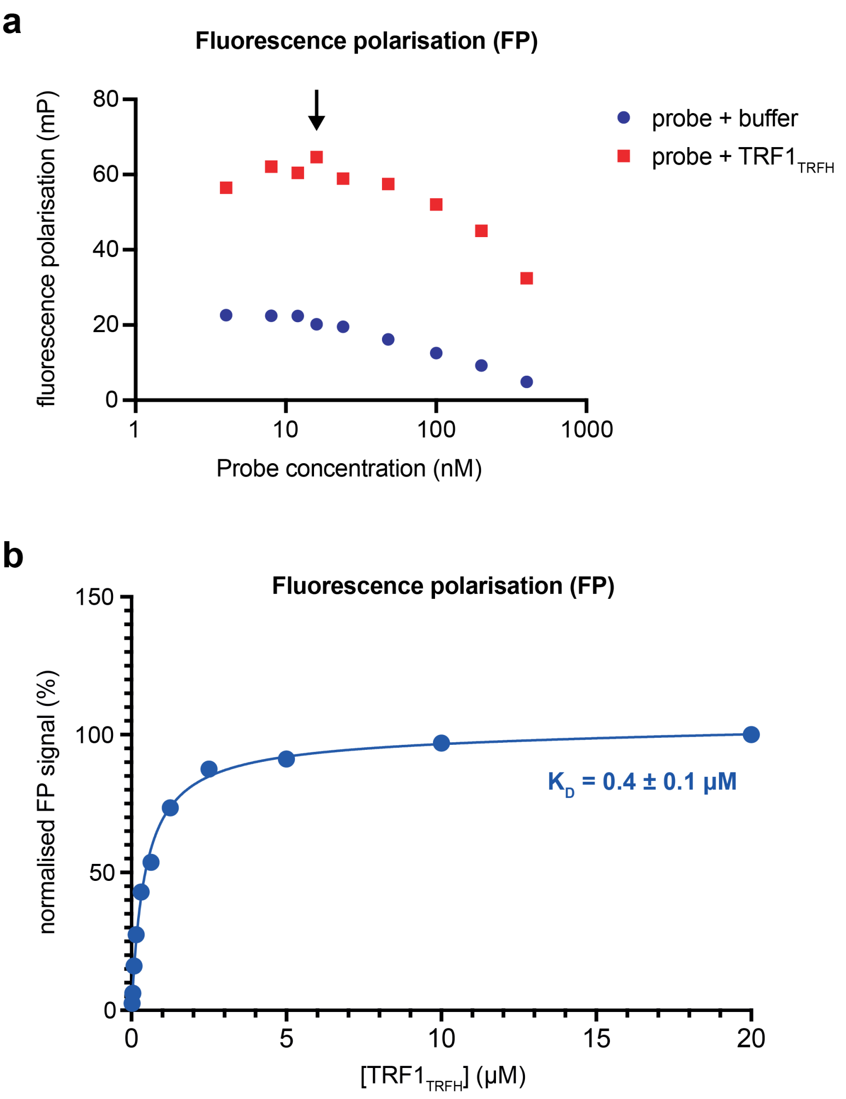

**Supplementary Figure 4: Establishing conditions for an FP competition assay. (a)** TIN2-FAM probe titration. The fluorescence polarisation signal was determined as a function of probe concentration both in the presence and absence of $TRF1_{TRFH}$. The black arrow indicates the probe concentration chosen for further experiments. **(b)** The FP signal of the TIN2-FAM probe as a function of $TRF1_{TRFH}$ domain concentration. Each data point is a mean value from two technical replicates of a single experiment. The calculated $K_D$ with standard error of the fit is indicated.

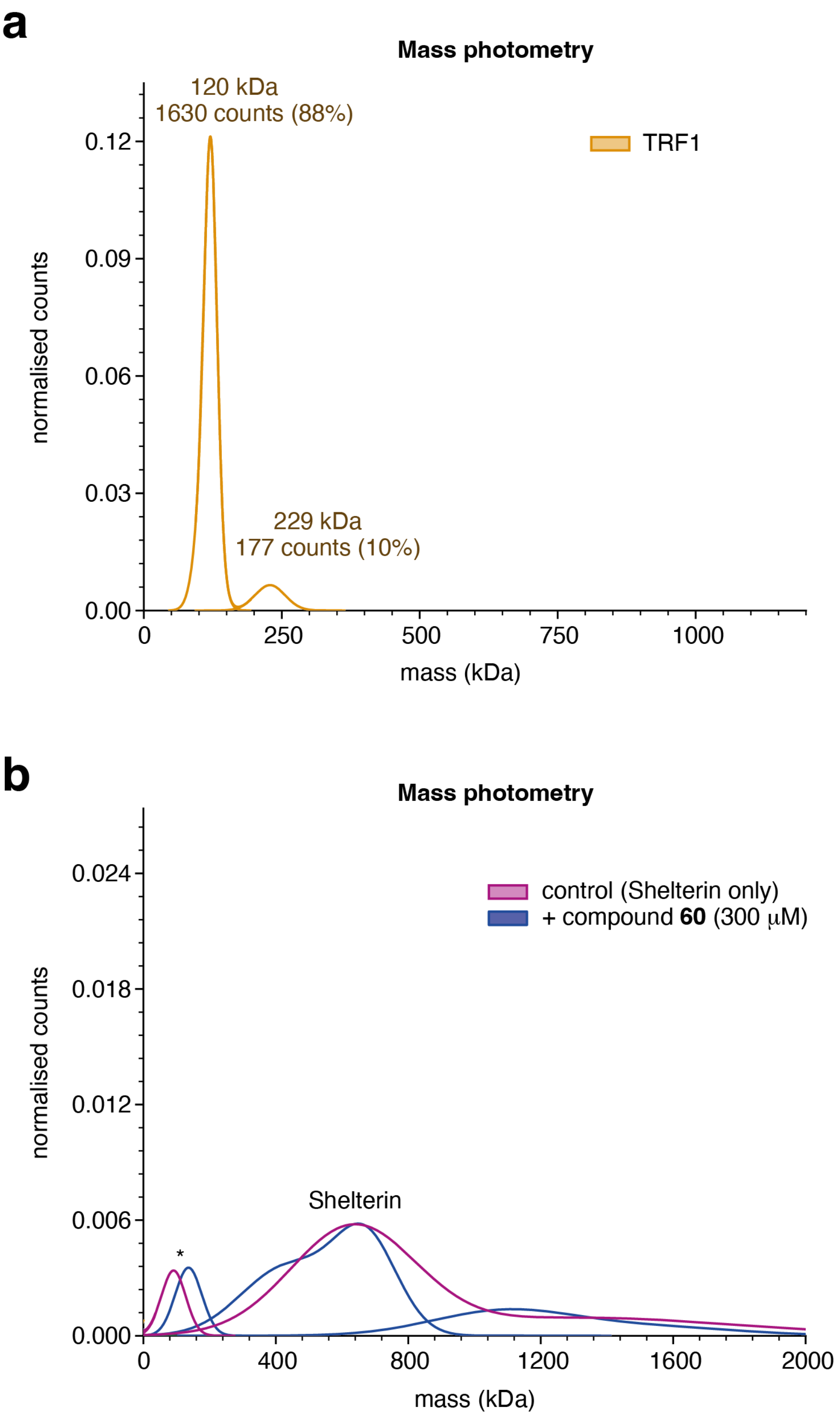

**Supplementary Figure 5: Controls for mass photometry assay.** Mass photometry Gaussian fits for **(a)** the full-length TRF1 sample and **(b)** shelterin with or without 300 µM of the inactive compound **60,** showing no eviction of TRF1 with the inactive compound**.** Asterisk (*) indicates a lower-molecular-weight contaminant in the Shelterin-only sample. Gaussian fits also indicate the presence of higher-molecular-weight species in the sample.

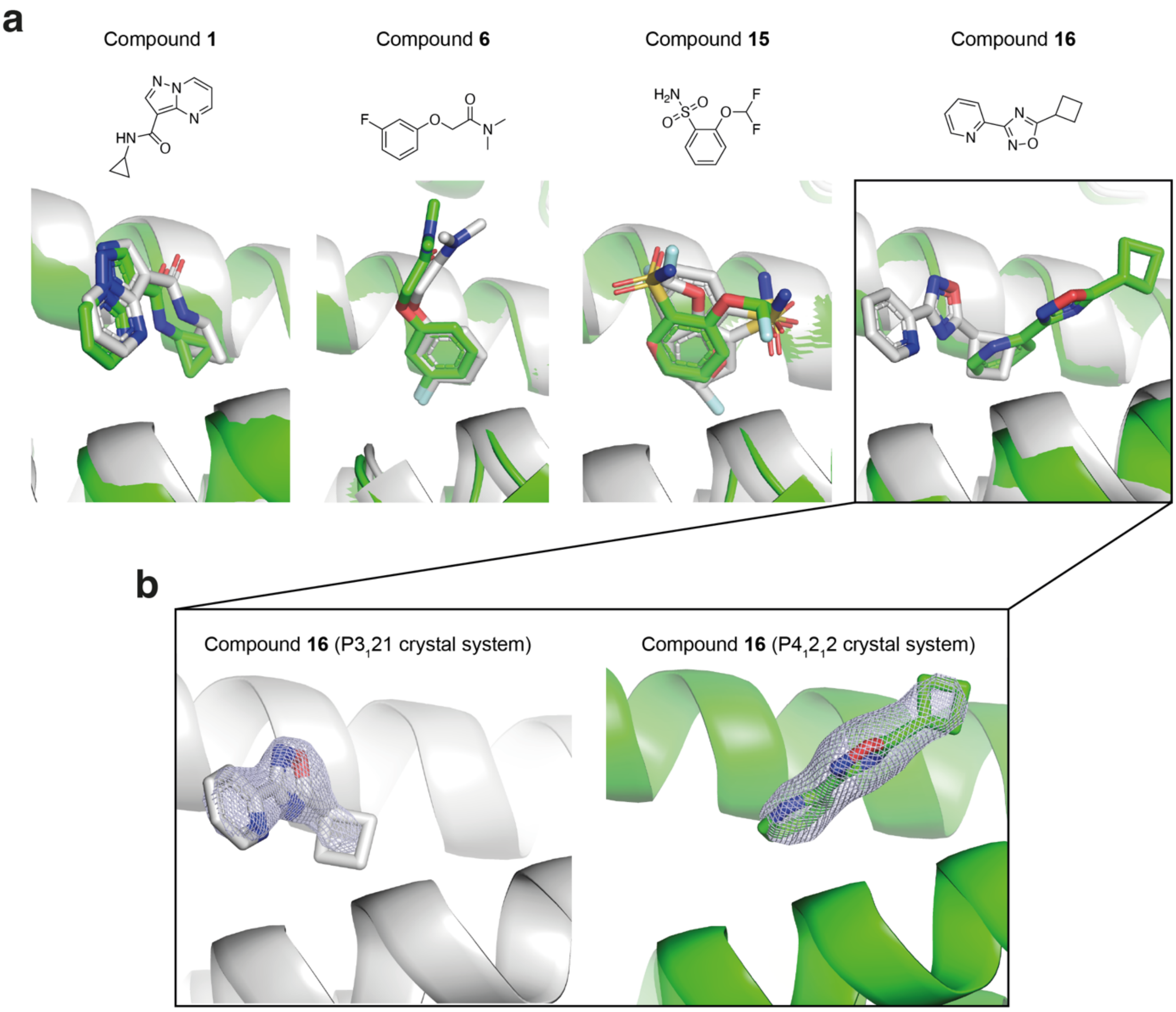

**Supplementary Figure 6: Fragment hits common to both XChem screens. (a)** Refined $TRF1_{TRFH}$:ligand structures showing the protein backbone in cartoon representation and bound ligands as sticks (coloured by heteroatom). Protein:ligand structural representations from $P3_121$ crystals are coloured in grey; those from $P4_12_12$ crystals are coloured in green. (**b**) Compound **16** bound to $TRF1_{TRFH}$ as per structures determined from $P3_121$ crystals and $P4_12_12$ crystals, in grey and green, respectively, with the Sigma-A weighted $2mF_o$-$DF_c$ electron density maps of ligands contoured at a level of 0.8.

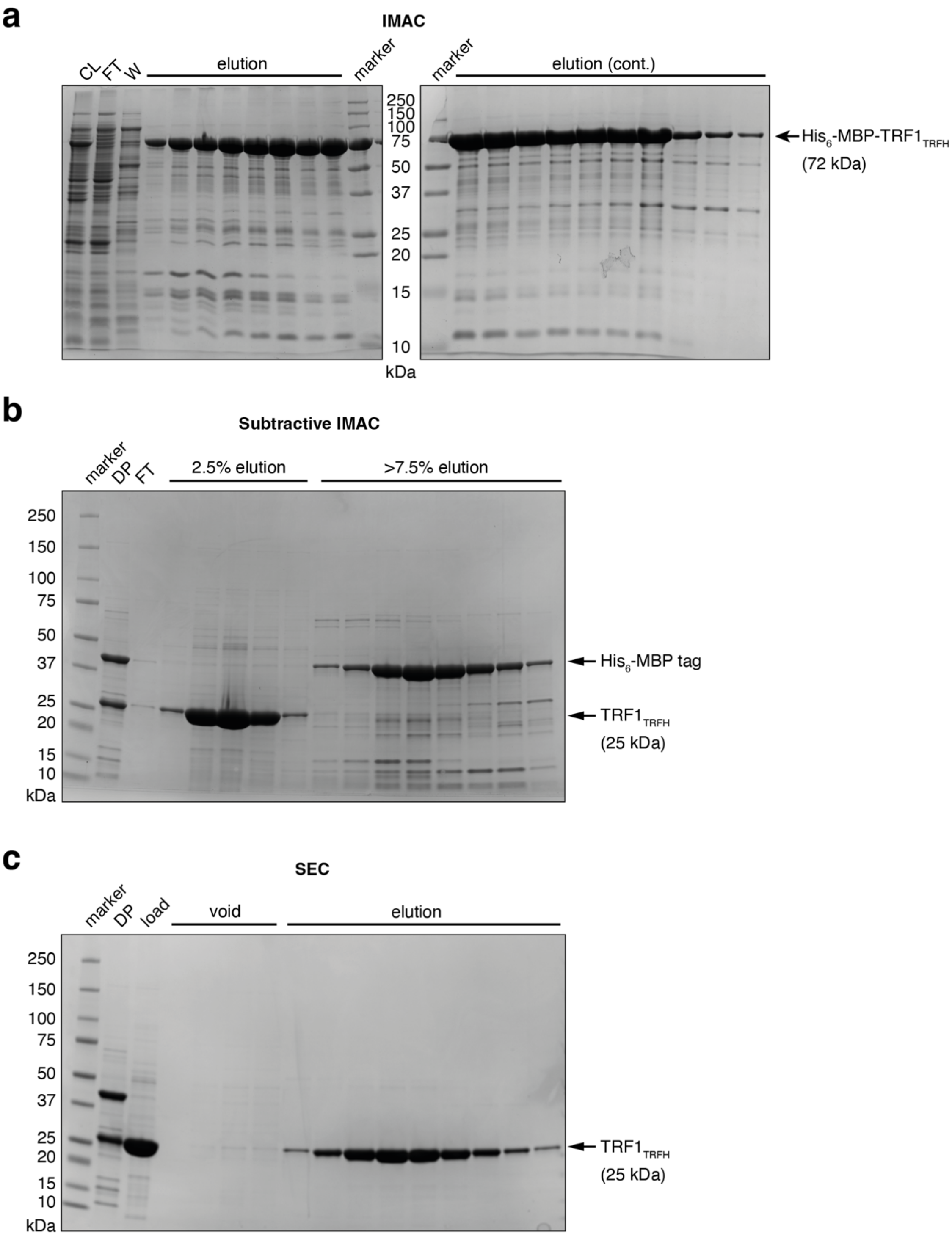

**Supplementary Figure 7: TRF1$_{TRFH}$ purification documented by SDS-PAGE and Coomassie staining. (a)** Immobilised metal ion affinity chromatography (IMAC) step: cell lysate (CL), flowthrough (FT), wash and eluted protein. **(b)** Subtractive IMAC following 6xHis-MBP tag cleavage using TEV protease: dialysed and cleaved protein (DP), flowthrough (FT) and stepwise elutions of protein with 2.5% elution buffer and >7.5% elution buffer. **(c)** Final size exclusion chromatography (SEC) purification: dialysed protein (DP), the protein loaded onto the column (Load), the void volume peak and the elution peak.

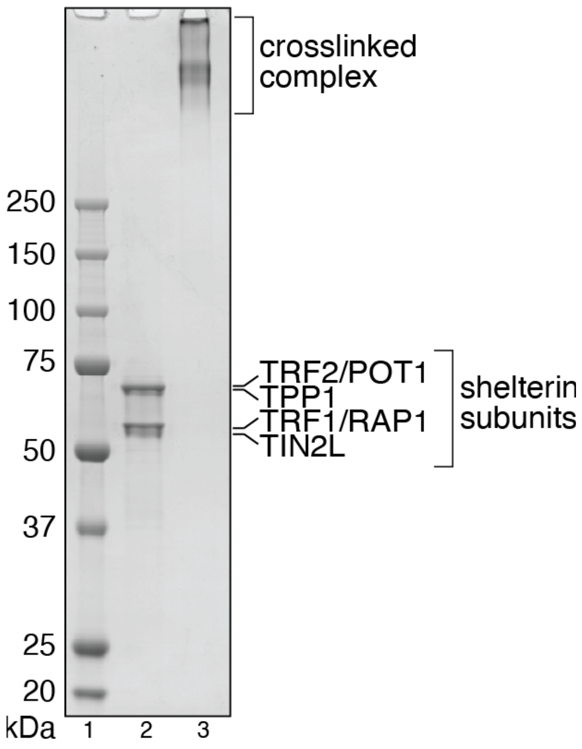

**Supplementary Figure 8: SDS-PAGE/Coomassie analysis of shelterin for mass photometry experiments.** Subunit labels for the purified shelterin complex before crosslinking correspond to lane 2. Lane 3 shows the complex upon crosslinking by addition of 0.1% glutaraldehyde for 25 min at room temperature.
