## Supplementary material for "Discovery of first-in-class inhibitors of the TRF1-TIN2 protein-protein interaction by fragment screening": Chemistry Supplementary Information

**Chemistry Supplementary Information for**

### Compound sources

All compounds that were commercially sourced were purchased as solid stocks with >90% purity and solubilised in-house in DMSO-d6 when required.
Compound **28** (4-methoxy-1-naphthalenesulfonic acid) was purchased from Merck Life Science/Sigma Aldrich (catalog number: S433853, CAS number: 84473-60-9)
Compound **29** (1-naphthol-4-sulfonic acid) was purchased from Combi Blocks Inc. (catalog number: SS-2420, CAS number: 84-87-7).
Compound **33** (4-ethoxynaphthalene-1-sulfonamide) was purchased from Enamine Ltd. (catalog number; Z385453312, CAS number: 861092-30-0
Compound **34** (4-ethoxy-1-naphthoic acid) was purchased from Fluorochem Ltd. (catalog number: F228774, CAS number: 19692-24-1)
Compound **35** was purchased from Enamine Ltd. (catalog number: Z283280258, CAS number: 299404-24-3).
Compound **36** was purchased from Enamine Ltd. (catalog number: Z815140124, CAS number: 299404-26-5)

### General synthetic experimental details

Starting materials and solvents were purchased from commercial suppliers and were used without further purification. NMR spectra were recorded on Bruker AMX500 or AV600 instruments using internal deuterium locks. Chemical shifts (δ) are reported relative to the solvent in which they were measured. Compounds were assessed for purity by tandem HPLC–MS. LC/MS analysis was performed on an Agilent 1260 Infinity II series UPLC and diode array detector coupled to a 6530 quadrupole time of flight mass spectrometer with Agilent Jet Stream ESI source using a negative ionisation method. Analytical separation was carried out at 40°C on a Phenomenex Kinetex C18 column (30 x 2.1 mm, 2.6u, 100A) using a flow rate of 0.5 mL/min in a 2 minute gradient elution with detection at 280 nm. The mobile phase was a mixture of water (solvent A) and methanol (solvent B), both containing formic acid at 0.1%. Gradient elution was as follows: 90:10 (A/B) to 10:90 (A/B) over 1.00 min, 10:90 (A/B) for 0.5 min, and then reversion back to 90:10 (A/B) over 0.2 min, finally 90:10 (A/B) for 0.3 min.

*General synthetic procedure*

4-hydroxynaphthalene-1-sulfonic acid (200 mg, 0.89 mmol, 1 eq) and sodium hydroxide (89 mg, 2.22 mmol, 2.5 eq) were dissolved in ethanol (0.6 mL, 0.75 M) and water (0.6 mL, 0.75 M) in a 2 mL microwave vial. Once the solution had cooled to room temperature the alkyl bromide (1.34 mmol, 1.5 eq) was added. The microwave vial was sealed and heated to 80 °C for 18 hrs.
The reaction mixture was cooled, and the precipitate was isolated by vacuum filtration, washing with dichloromethane, to give the desired product as the sodium salt.

**LC/MS, $^{1}$H NMR and $^{13}$C NMR spectra of compound 27**

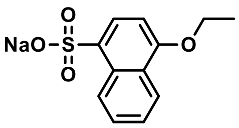


Following the general procedure 4-ethoxynaphthalene-1-sulfonic acid (57 mg, 23%, 0.225 mmol) was isolated as the sodium salt.

$^{1}$H NMR (600 MHz, DMSO) δ 8.78 (ddd, J = 8.5, 1.4, 0.7 Hz, 1H), 8.16 (ddd, J = 8.3, 1.5, 0.8 Hz, 1H), 7.85 (d, J = 8.0 Hz, 1H), 7.51 (ddd, J = 8.4, 6.7, 1.5 Hz, 1H), 7.47 (ddd, J = 8.2, 6.7, 1.4 Hz, 1H), 6.85 (d, J = 8.1 Hz, 1H), 4.22 (q, J = 7.0 Hz, 2H), 1.48 (t, J = 6.9 Hz, 3H).

$^{13}$C NMR (151 MHz, DMSO) δ 154.5, 136.5, 130.1, 127.6, 125.8, 125.1, 125.1, 124.7, 121.2, 103.0, 63.5, 14.6.

MS (m/z): [M-Na]$^{-}$ 251

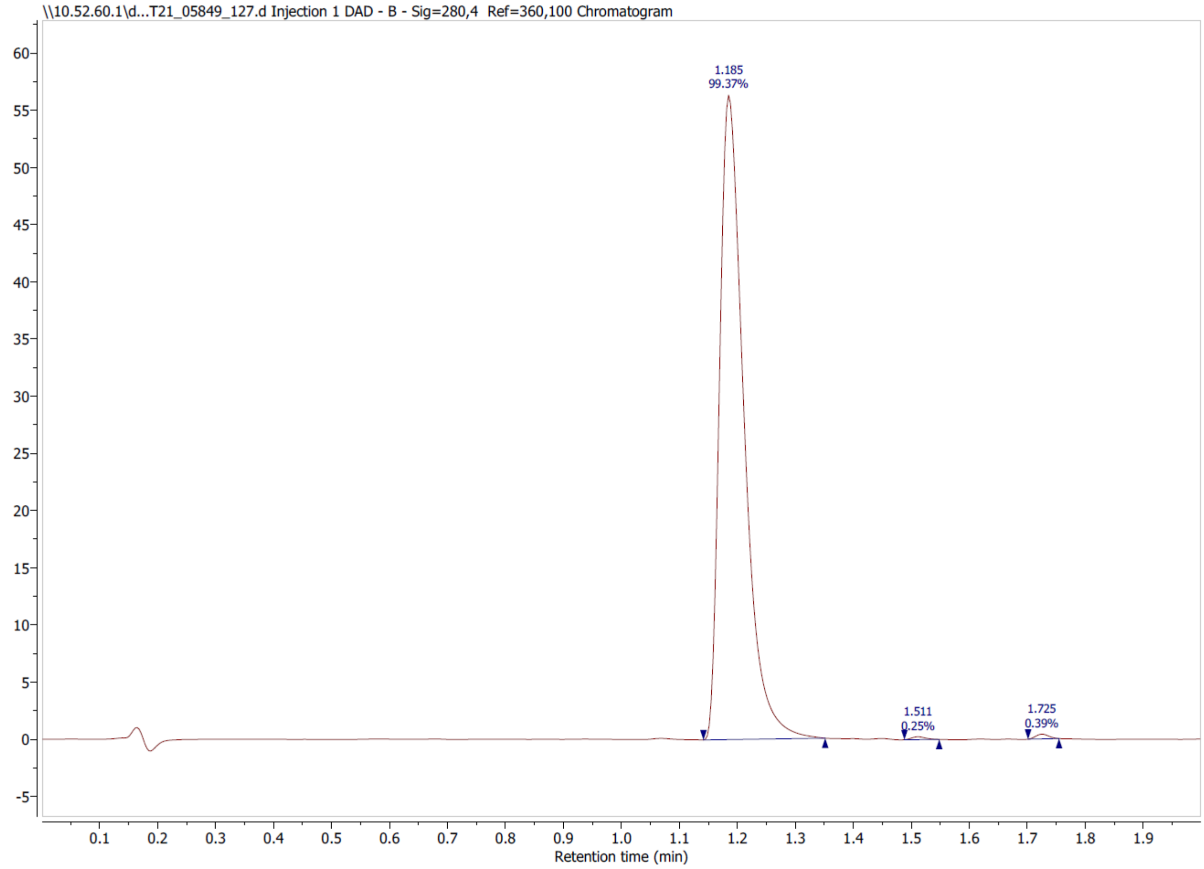

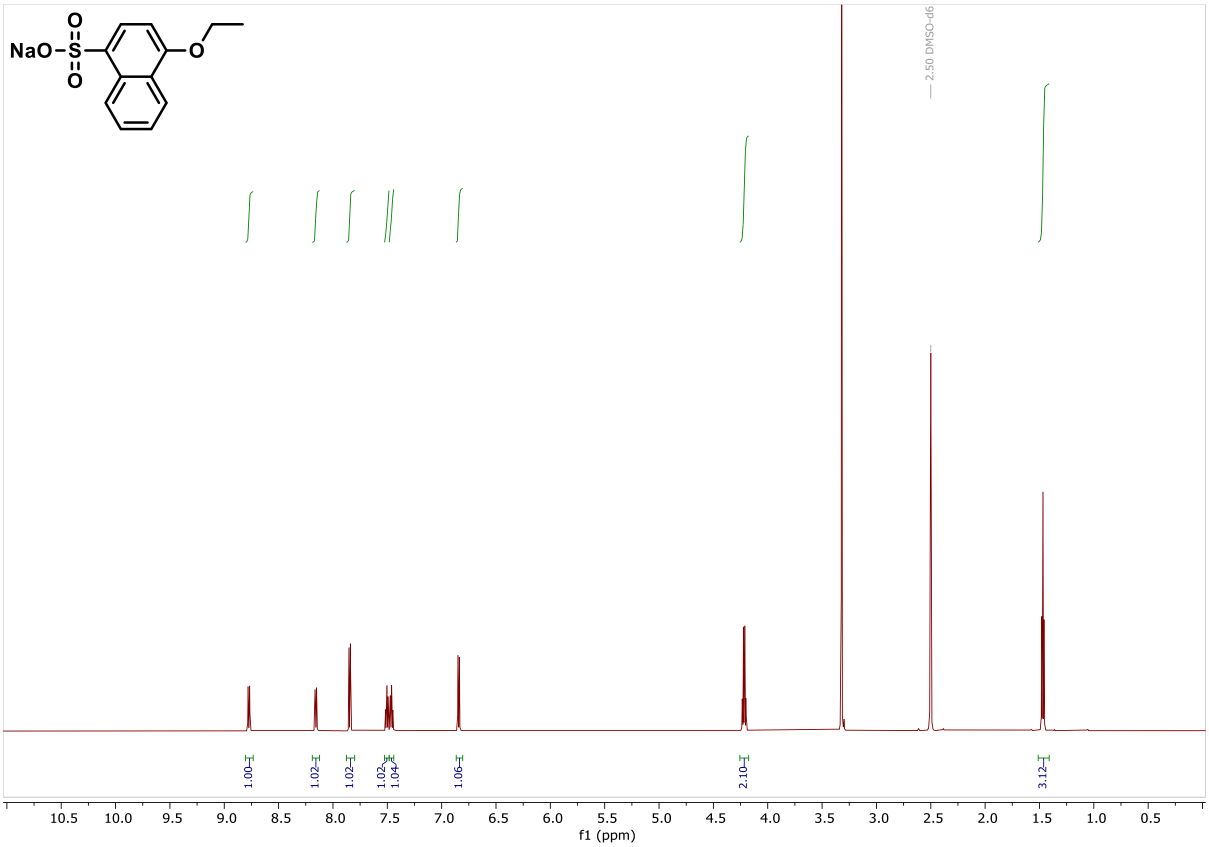
NaO
2.50 DMSO-d6
1.00
1.02
1.02
1.02
1.04
1.06
2.10
3.12
f1 (ppm)

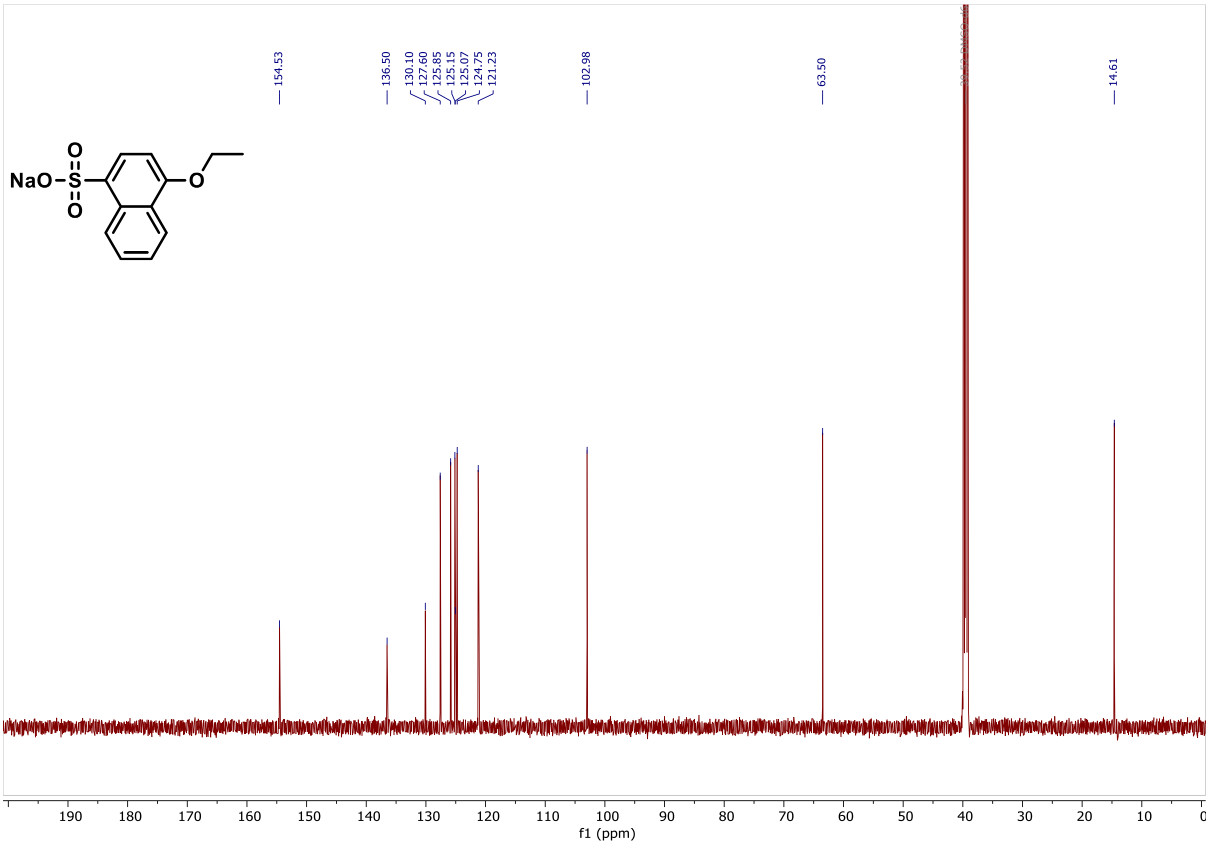
NaO
154.53
136.50
130.10
127.60
125.85
125.15
125.07
124.75
121.23
102.98
63.50
14.61
f1 (ppm)

#### LC/MS, $^1$H NMR and $^{13}$C NMR spectra of compound 30

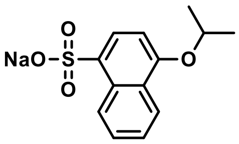


Following the general procedure 4-isopropoxynaphthalene-1-sulfonic acid (111 mg, 43%, 0.417 mmol) was isolated as the sodium salt.

$^1$H NMR (600 MHz, DMSO) δ 8.77 (ddd, J = 8.6, 1.4, 0.7 Hz, 1H), 8.14 (ddd, J = 8.3, 1.5, 0.7 Hz, 1H), 7.85 (d, J = 8.1 Hz, 1H), 7.49 (ddd, J = 8.4, 6.7, 1.5 Hz, 1H), 7.44 (ddd, J = 8.2, 6.7, 1.4 Hz, 1H), 6.89 (d, J = 8.1 Hz, 1H), 4.88 – 4.79 (m, 1H), 1.38 (d, J = 6.0 Hz, 6H).

$^{13}$C NMR (151 MHz, DMSO) δ 153.4, 136.3, 130.3, 127.5, 125.8, 125.8, 125.2, 124.7, 121.5, 104.4, 69.7, 21.8.

MS (m/z): [M-Na]$^-$ 265

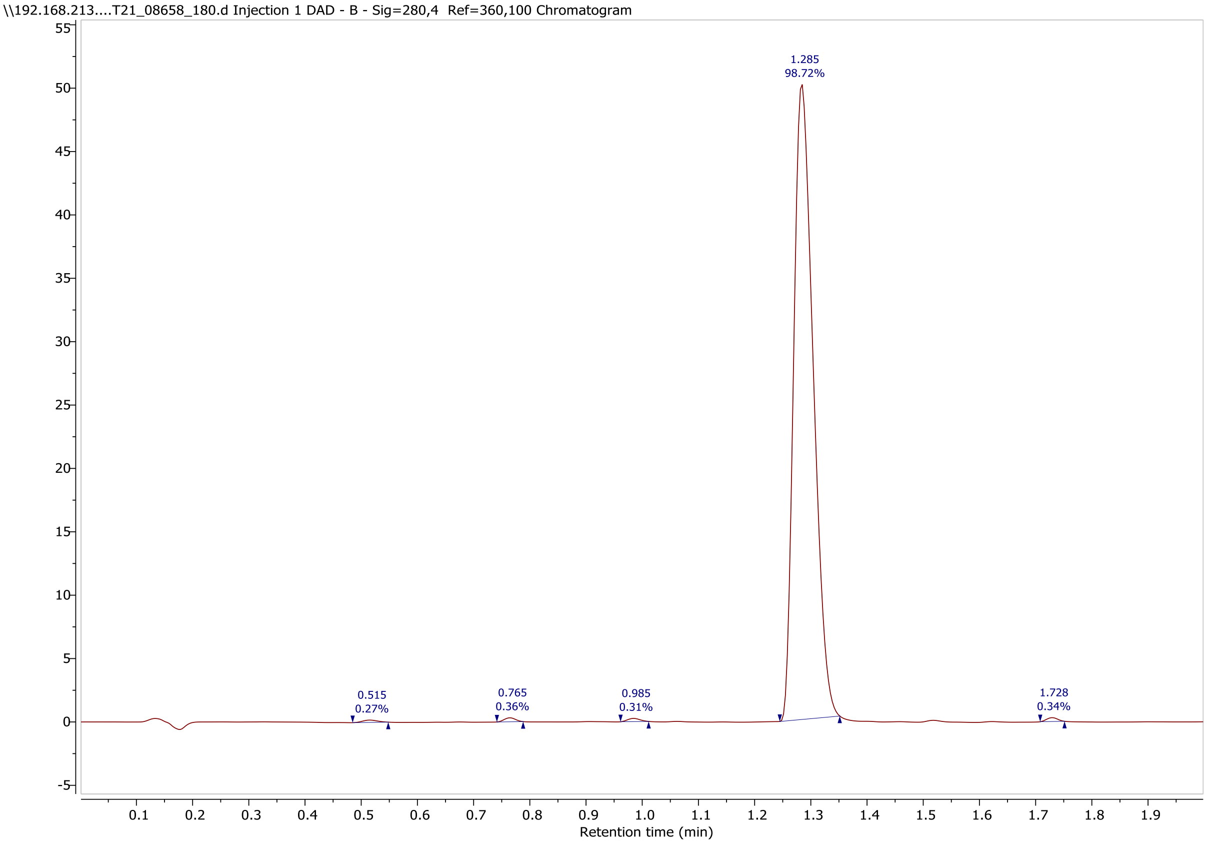

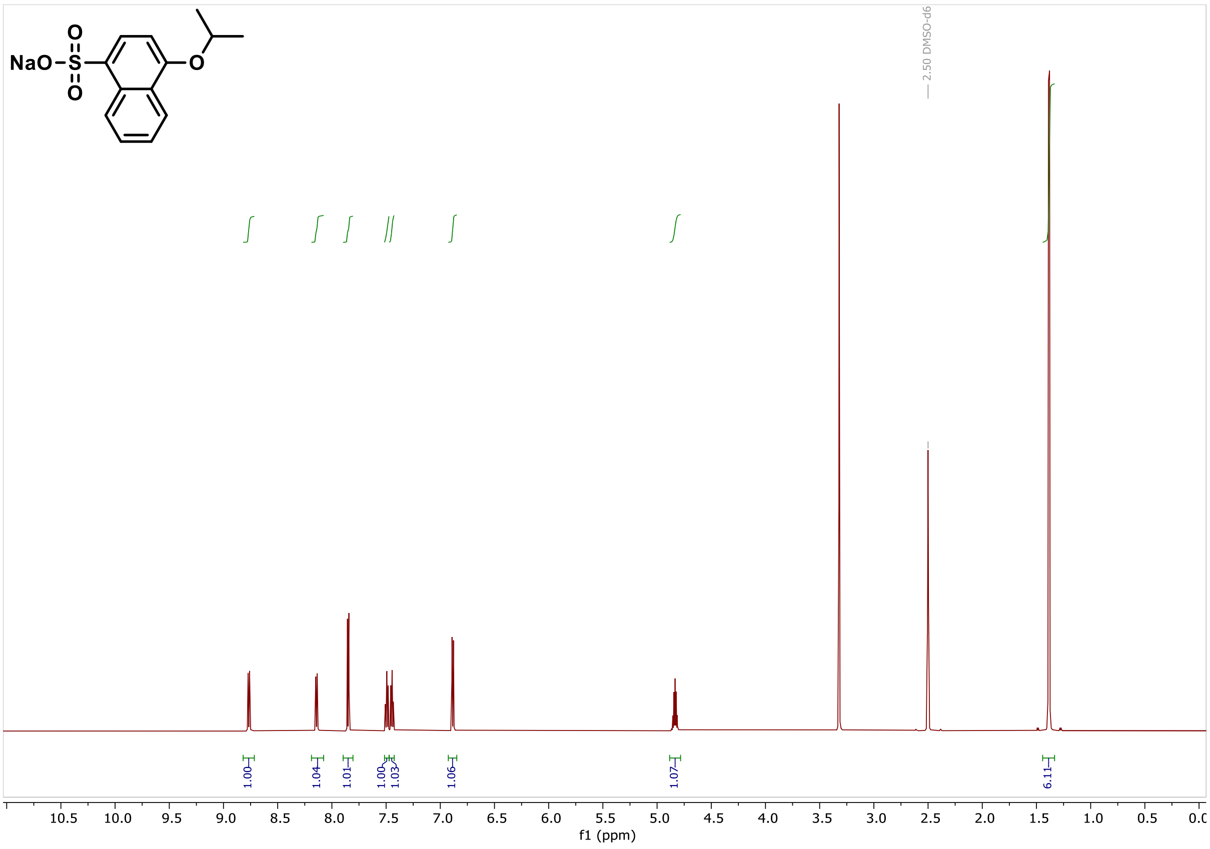
NaO-S(O)(O)- O
2.50 DMSO-d6
1.00
1.04
1.01
1.00
1.03
1.06
1.07
6.11
10.5 10.0 9.5 9.0 8.5 8.0 7.5 7.0 6.5 6.0 5.5 5.0 4.5 4.0 3.5 3.0 2.5 2.0 1.5 1.0 0.5 0.0
f1 (ppm)

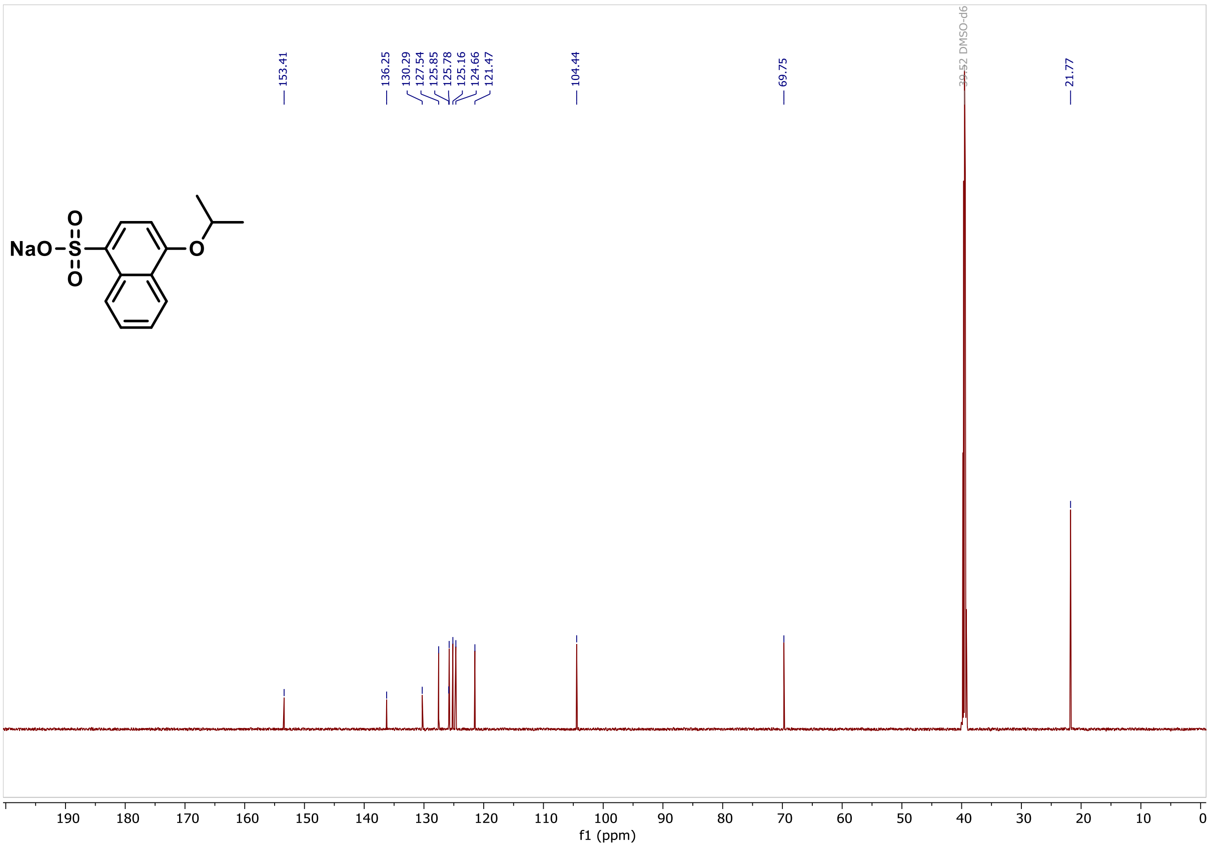
153.41
136.25
130.29
127.54
125.85
125.78
125.16
124.66
121.47
104.44
69.75
39.52 DMSO-d6
21.77
NaO-S(O)(O)- O
190 180 170 160 150 140 130 120 110 100 90 80 70 60 50 40 30 20 10 0
f1 (ppm)

**LC/MS, $^1$H NMR and $^{13}$C NMR spectra of compound 31**

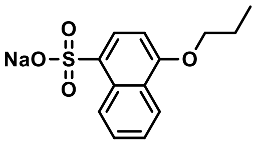

Following the general procedure 4-propoxynaphthalene-1-sulfonic acid (121 mg, 47%, 0.454 mmol) was isolated as the sodium salt.

$^1$H NMR (600 MHz, DMSO) δ 8.82 – 8.74 (m, 1H), 8.17 (ddd, J = 8.2, 1.6, 0.7 Hz, 1H), 7.85 (d, J = 8.0 Hz, 1H), 7.51 (ddd, J = 8.4, 6.7, 1.6 Hz, 1H), 7.47 (ddd, J = 8.1, 6.7, 1.4 Hz, 1H), 6.85 (d, J = 8.1 Hz, 1H), 4.12 (t, J = 6.4 Hz, 2H), 1.88 (dtd, J = 13.7, 7.4, 6.4 Hz, 2H), 1.08 (t, J = 7.4 Hz, 3H).

$^{13}$C NMR (151 MHz, DMSO) δ 154.6, 136.4, 130.1, 127.6, 125.9, 125.2, 125.1, 124.8, 121.2, 103.0, 69.2, 22.1, 10.6.

MS (m/z): $[M\text{-}Na]^-$ 251

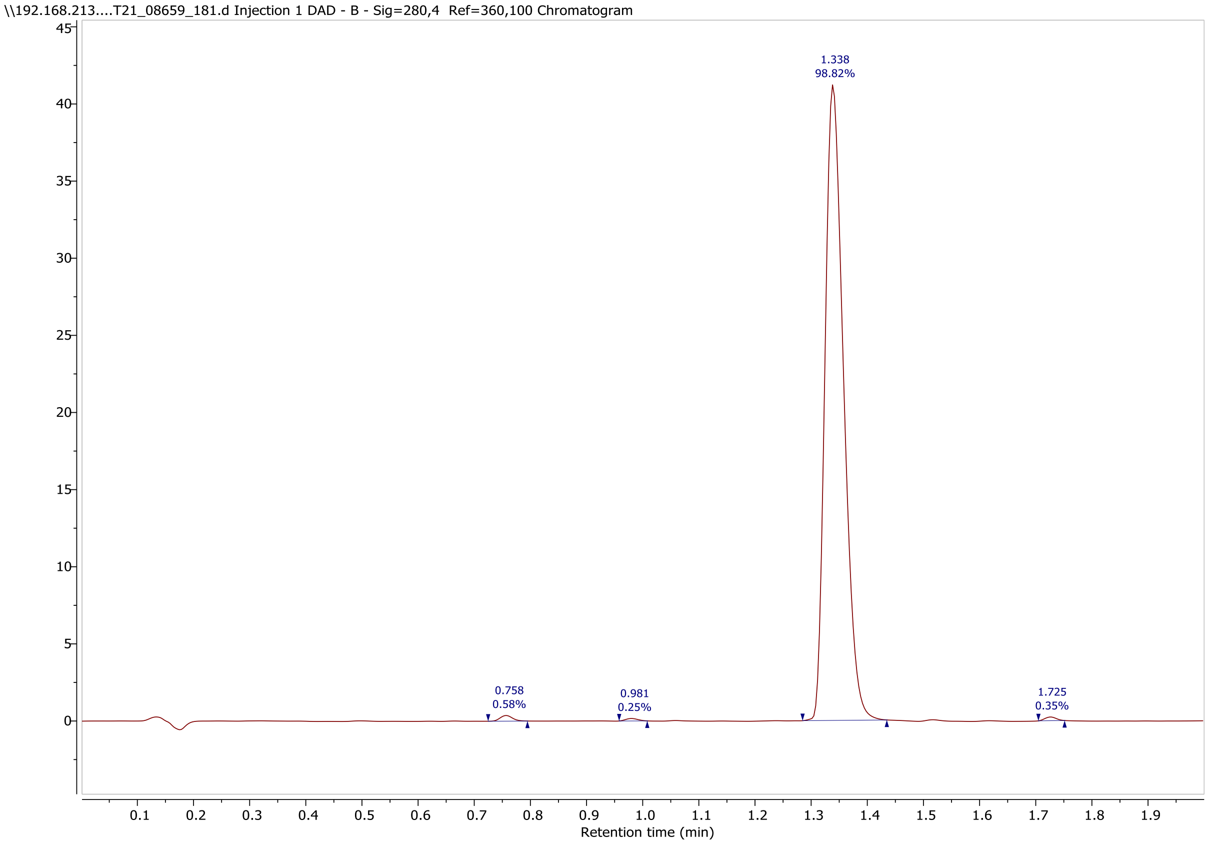

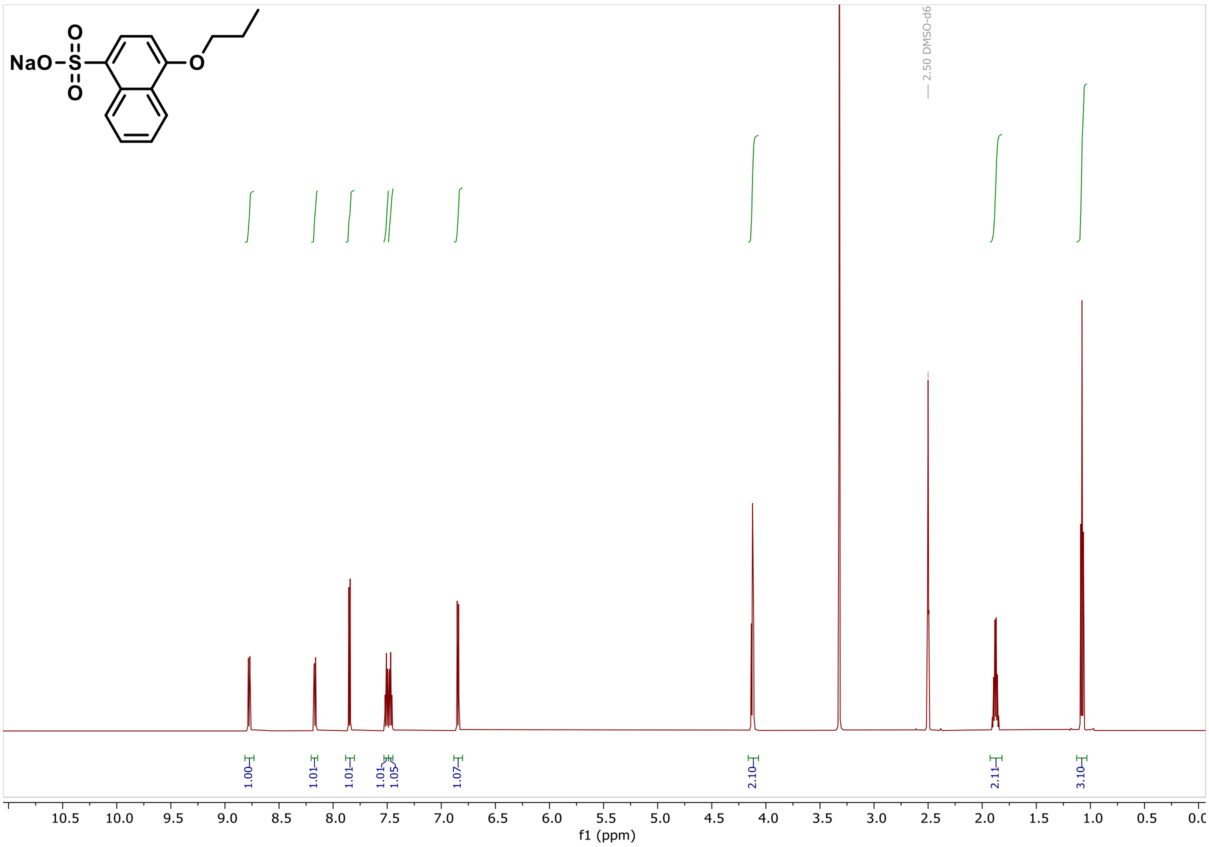
NaO-S-O
O
O
2.50 DMSO-d6
1.00
1.01
1.01
1.01
1.05
1.07
2.10
2.11
3.10
10.5 10.0 9.5 9.0 8.5 8.0 7.5 7.0 6.5 6.0 5.5 5.0 4.5 4.0 3.5 3.0 2.5 2.0 1.5 1.0 0.5 0.0
f1 (ppm)

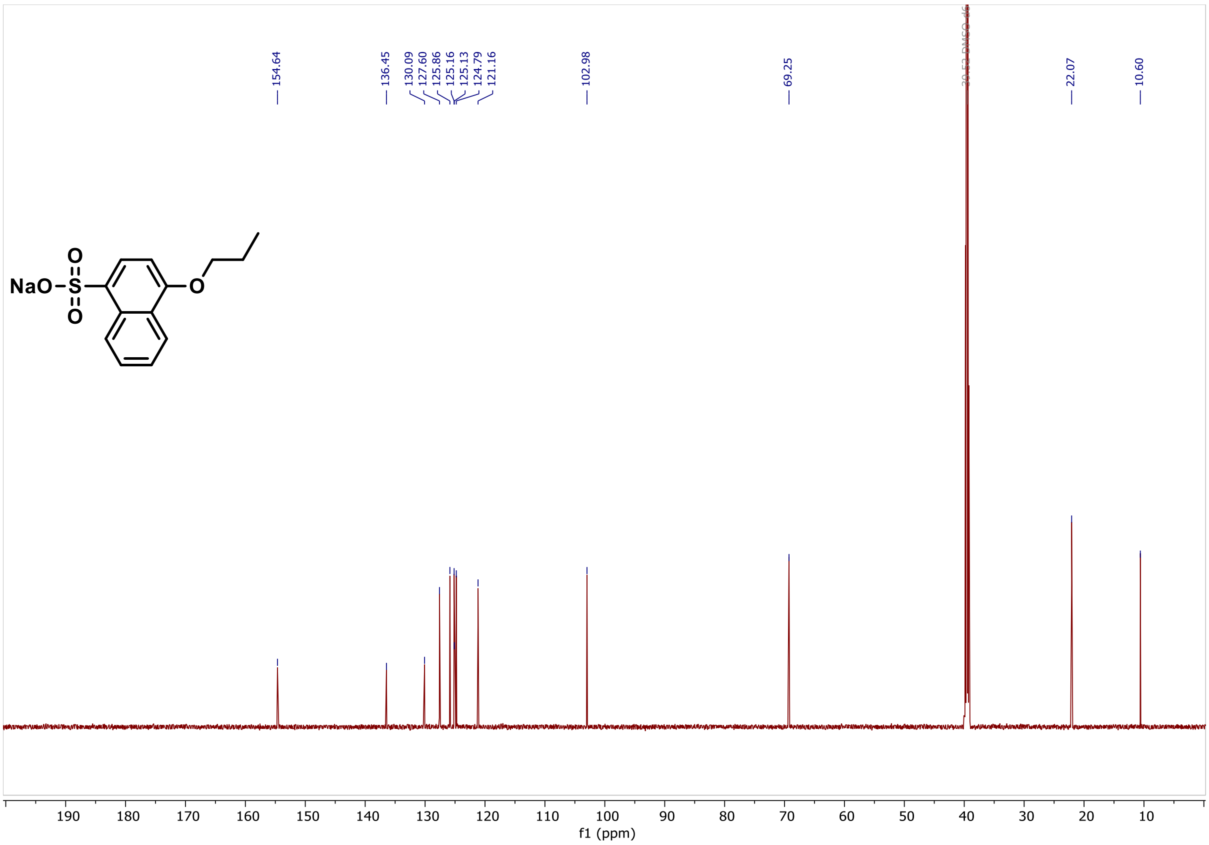
154.64
136.45
130.09
127.60
125.86
125.16
125.13
124.79
121.16
102.98
69.25
22.07
10.60
NaO-S-O
O
O
190 180 170 160 150 140 130 120 110 100 90 80 70 60 50 40 30 20 10
f1 (ppm)

**LC/MS, [1]H NMR and [13]C NMR spectra of compound 32**

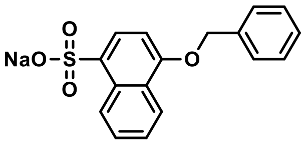

Following the general procedure 4-benzyloxynaphthalene-1-sulfonic acid (113 mg, 38%, 0.360 mmol) was isolated as the sodium salt.

$^{1}$H NMR (600 MHz, DMSO) δ 8.82 – 8.77 (m, 1H), 8.23 – 8.17 (m, 1H), 7.86 (d, J = 8.0 Hz, 1H), 7.57 – 7.54 (m, 2H), 7.52 (ddd, J = 8.4, 6.7, 1.5 Hz, 1H), 7.48 (ddd, J = 8.1, 6.7, 1.4 Hz, 1H), 7.45 – 7.41 (m, 2H), 7.39 – 7.31 (m, 1H), 6.98 (d, J = 8.1 Hz, 1H), 5.32 (s, 2H).

$^{13}$C NMR (151 MHz, DMSO) δ 154.2, 137.0, 136.9, 130.1, 128.5, 127.8, 127.6, 127.5, 125.9, 125.2, 125.0, 121.2, 103.7, 69.4.

MS (m/z): $[M\text{-}Na]^{-}$ 313

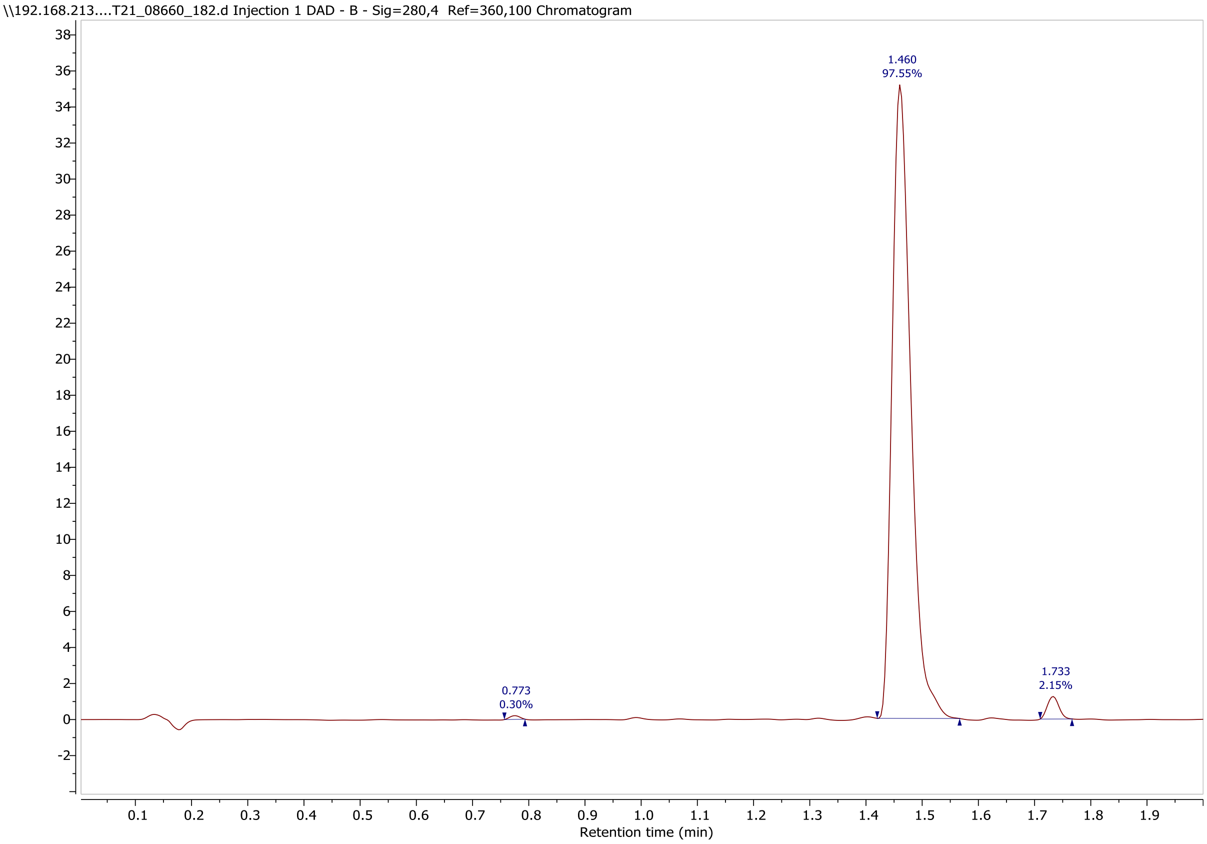

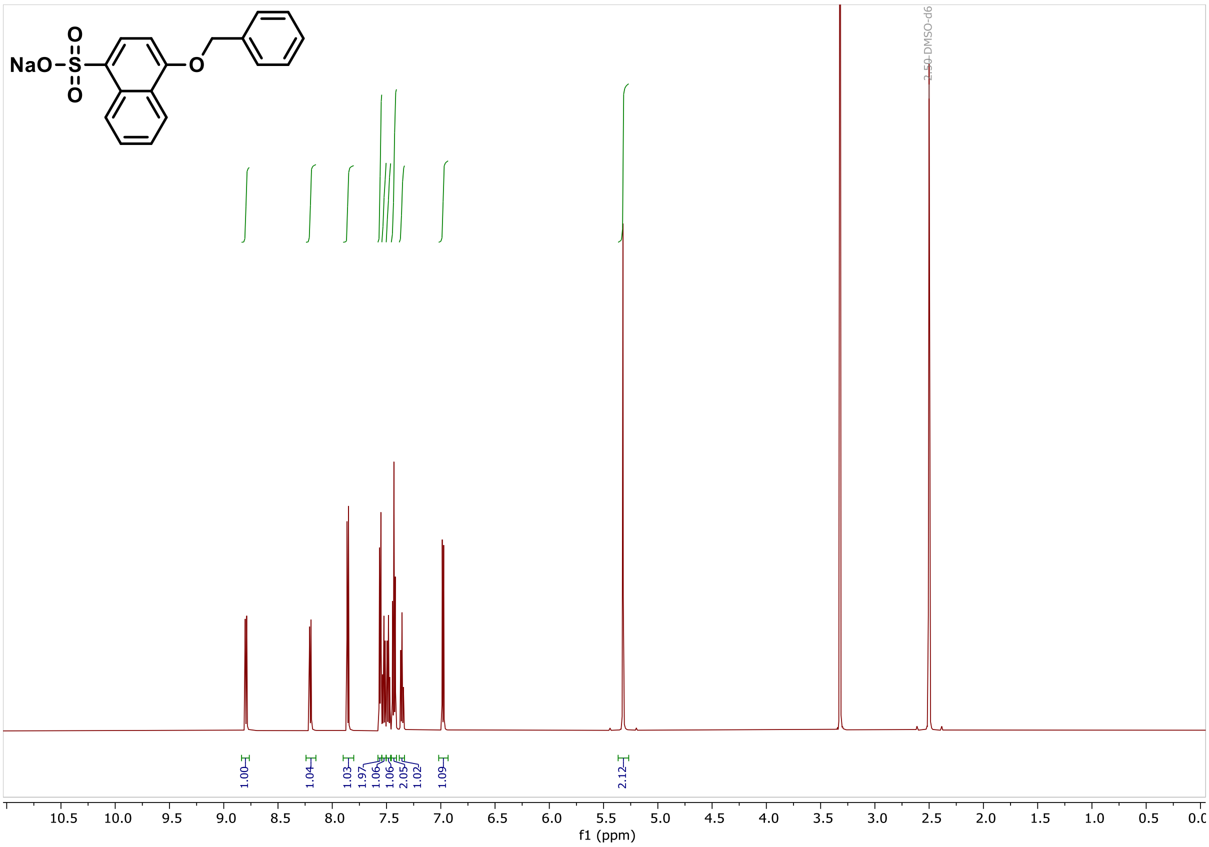
NaO
O
O
S
O
2.50 DMSO-d6
1.00
1.04
1.03
1.97
1.06
1.06
2.05
1.02
1.09
2.12
10.5
10.0
9.5
9.0
8.5
8.0
7.5
7.0
6.5
6.0
5.5
5.0
4.5
4.0
3.5
3.0
2.5
2.0
1.5
1.0
0.5
f1 (ppm)

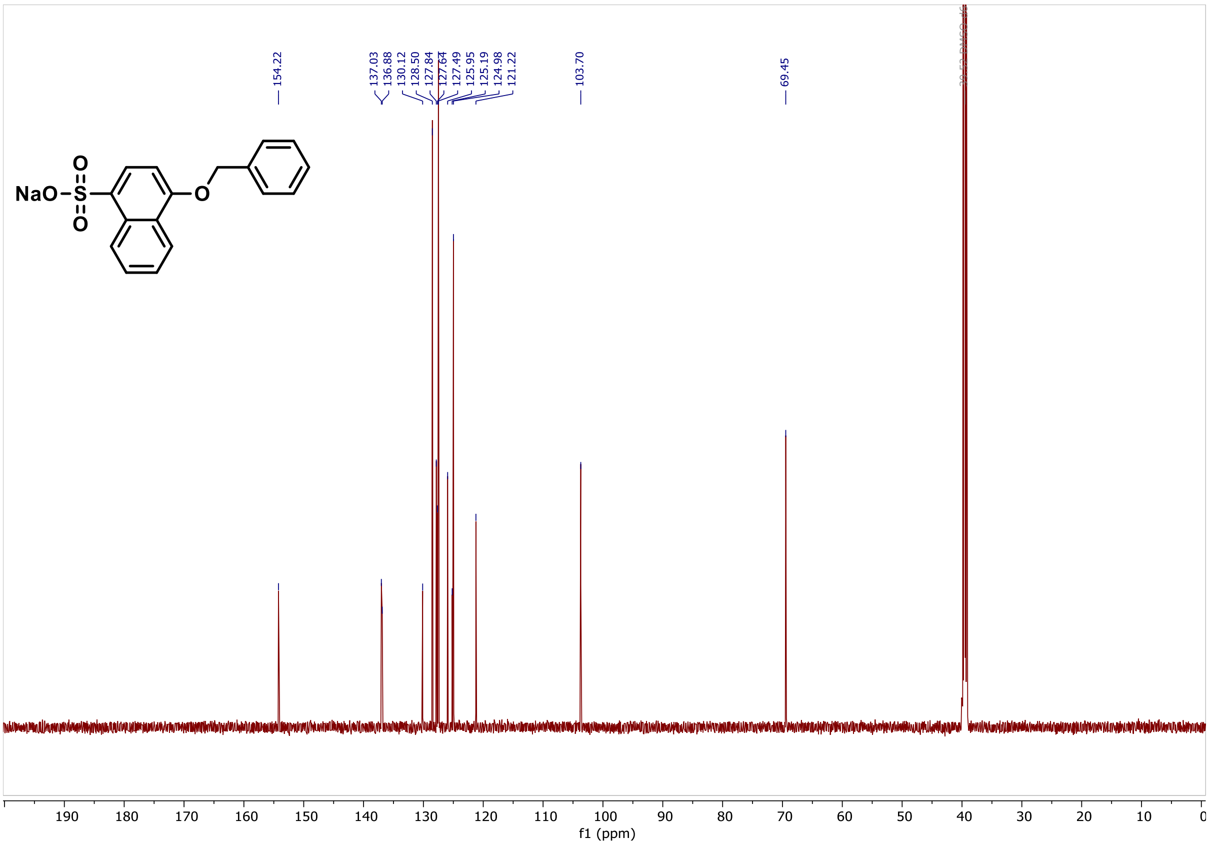
154.22
137.03
136.88
130.12
128.50
127.84
127.64
127.49
125.95
125.19
124.98
121.22
103.70
69.45
NaO
O
O
S
O
190
180
170
160
150
140
130
120
110
100
90
80
70
60
50
40
30
20
10
f1 (ppm)

**LC/MS, $^{1}$H NMR and $^{13}$C NMR spectra of compound 37**

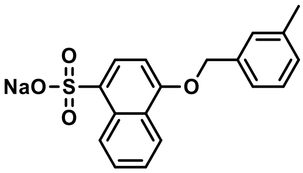

Following the general procedure 4-(m-tolylmethoxy)naphthalene-1-sulfonic acid (99.1 mg, 32%, 0.302 mmol) was isolated as the sodium salt.

$^{1}$H NMR (600 MHz, DMSO) δ 8.79 (d, *J* = 8.5 Hz, 1H), 8.20 (d, *J* = 7.6 Hz, 1H), 7.86 (d, *J* = 8.0 Hz, 1H), 7.53 (ddd, *J* = 8.5, 6.7, 1.5 Hz, 1H), 7.51 – 7.44 (m, 1H), 7.39 – 7.26 (m, 3H), 7.17 (d, *J* = 7.3 Hz, 1H), 6.97 (d, *J* = 8.1 Hz, 1H), 5.28 (s, 2H), 2.34 (s, 3H).

$^{13}$C NMR (151 MHz, DMSO) δ 154.3, 137.7, 136.9, 136.7, 130.1, 128.5, 128.4, 128.1, 127.6, 126.0, 125.2, 125.1, 125.0, 124.6, 121.3, 103.7, 69.5, 21.1.

MS (m/z): $[M\text{-}Na]^-$ 327.1

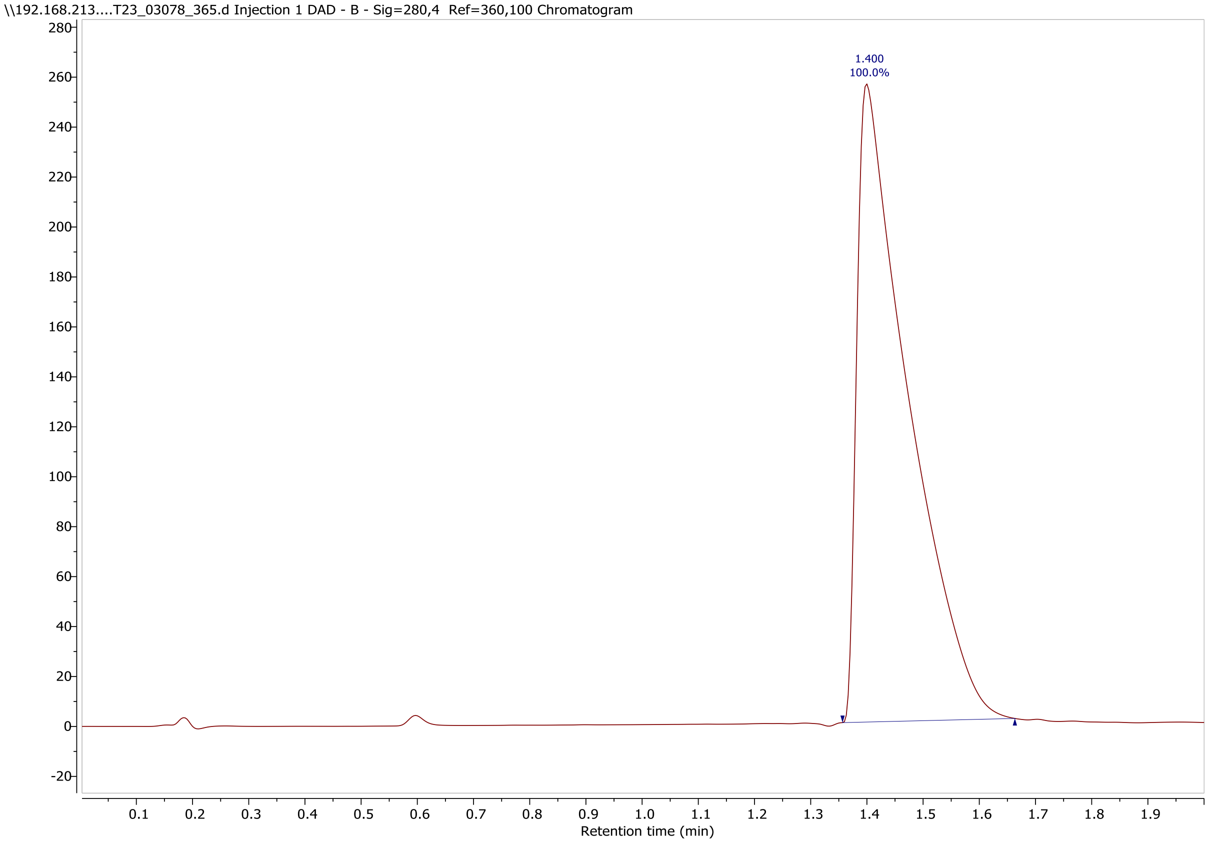

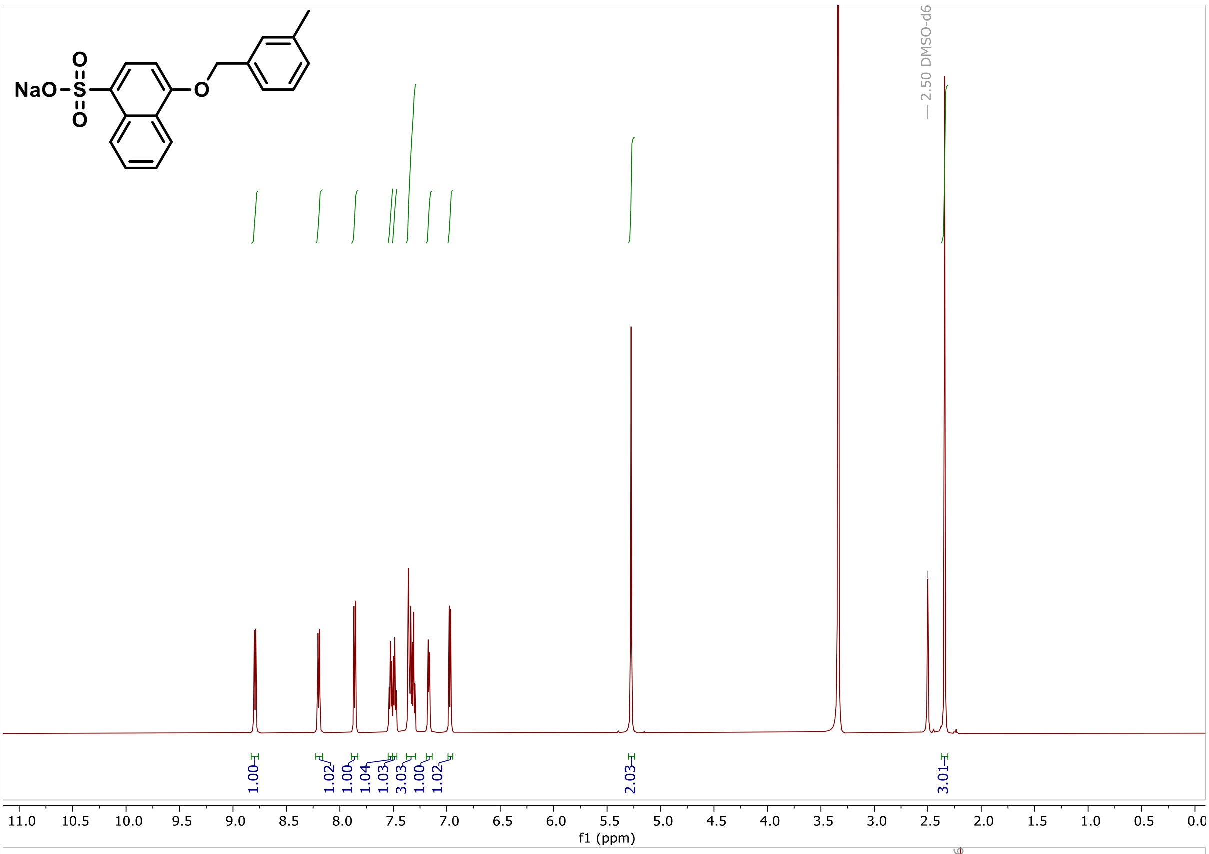
NaO
S
O
O
O
2.50 DMSO-d6
1.00
1.02
1.00
1.04
1.03
3.03
1.00
1.02
2.03
3.01
f1 (ppm)

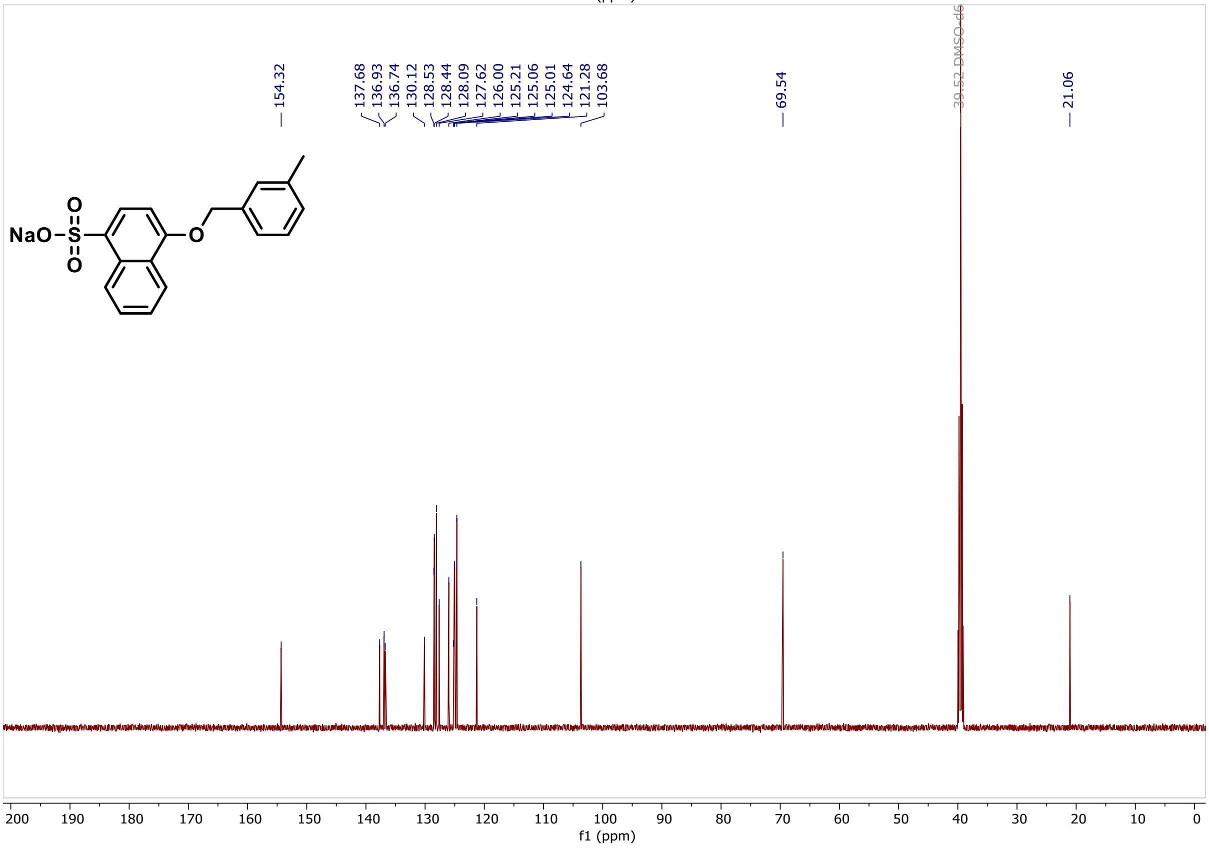
154.32
137.68
136.93
136.74
130.12
128.53
128.44
128.09
127.62
126.00
125.21
125.06
125.01
124.64
121.28
103.68
69.54
39.52 DMSO-d6
21.06
NaO
S
O
O
O
f1 (ppm)

**LC/MS, [1]H NMR and [13]C NMR spectra of compound 38**

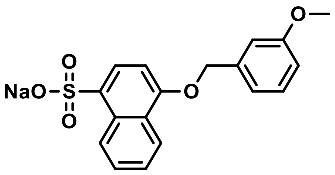


Following the general procedure 4-[(3-methoxyphenyl)methoxy]naphthalene-1-sulfonic acid (72 mg, 22%, 0.209 mmol) was isolated as the sodium salt.

$^{1}$H NMR (500 MHz, DMSO) δ 8.83 – 8.76 (m, 1H), 8.24 – 8.18 (m, 1H), 7.86 (d, $J$ = 8.0 Hz, 1H), 7.51 (dddd, $J$ = 18.4, 8.1, 6.8, 1.5 Hz, 2H), 7.37 – 7.30 (m, 1H), 7.14 – 7.08 (m, 2H), 6.97 (d, $J$ = 8.2 Hz, 1H), 6.92 (ddd, $J$ = 8.3, 2.4, 1.2 Hz, 1H), 5.29 (s, 2H), 3.77 (s, 3H).

$^{13}$C NMR (151 MHz, DMSO) δ 159.9, 154.7, 139.1, 137.3, 130.6, 130.1, 128.1, 126.5, 125.7, 125.5, 125.5, 121.7, 120.0, 113.7, 113.5, 104.2, 69.8, 55.5.

MS (m/z): [M-Na]$^{-}$ 343.0

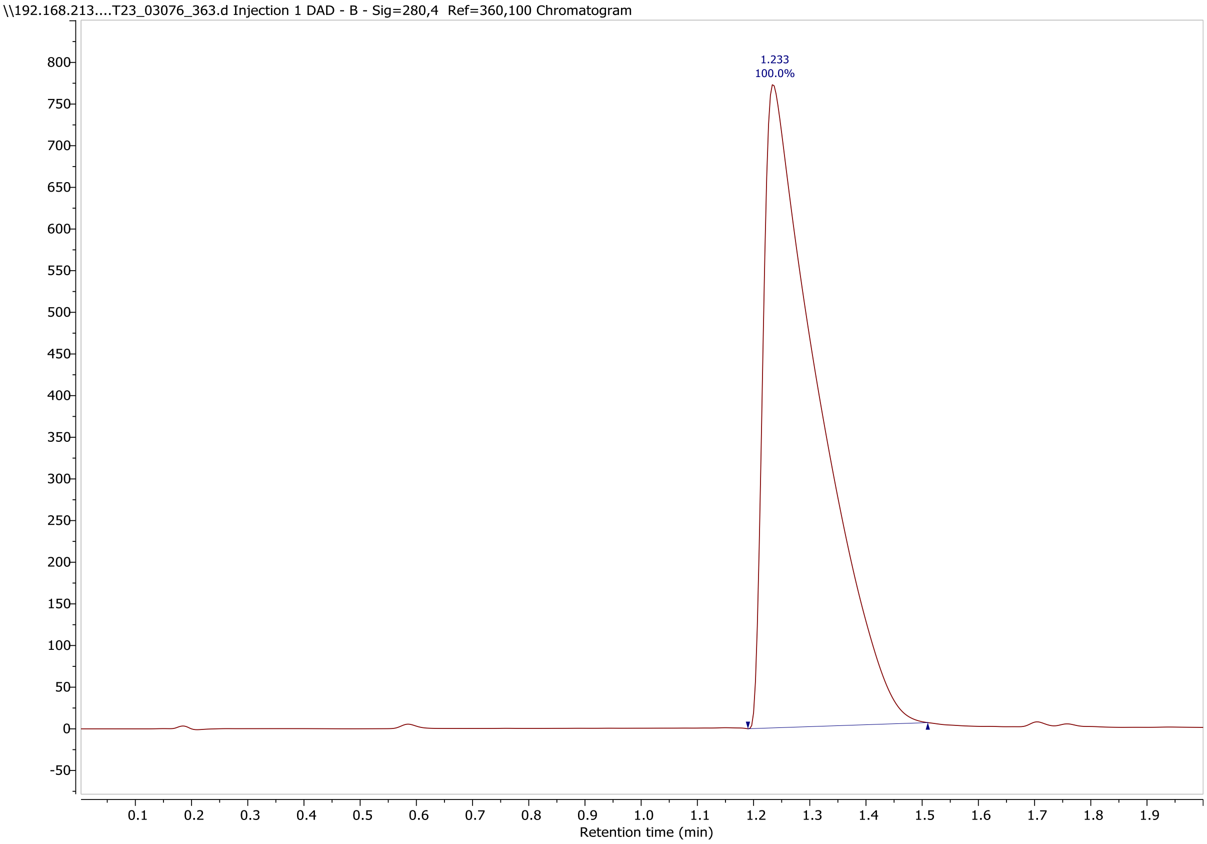

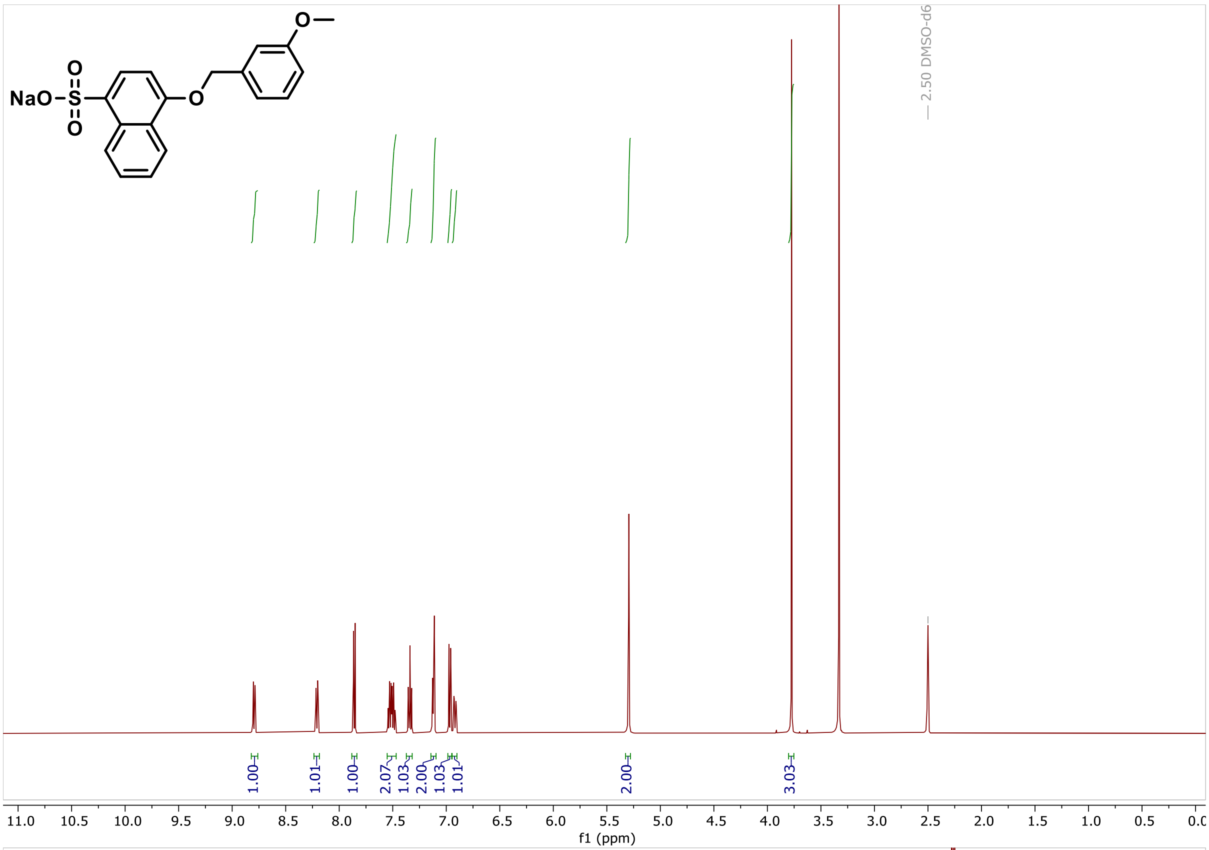
NaO
O
S
O
O
O
2.50 DMSO-d6
1.00
1.01
1.00
2.07
1.03
2.00
1.03
1.01
2.00
3.03
f1 (ppm)

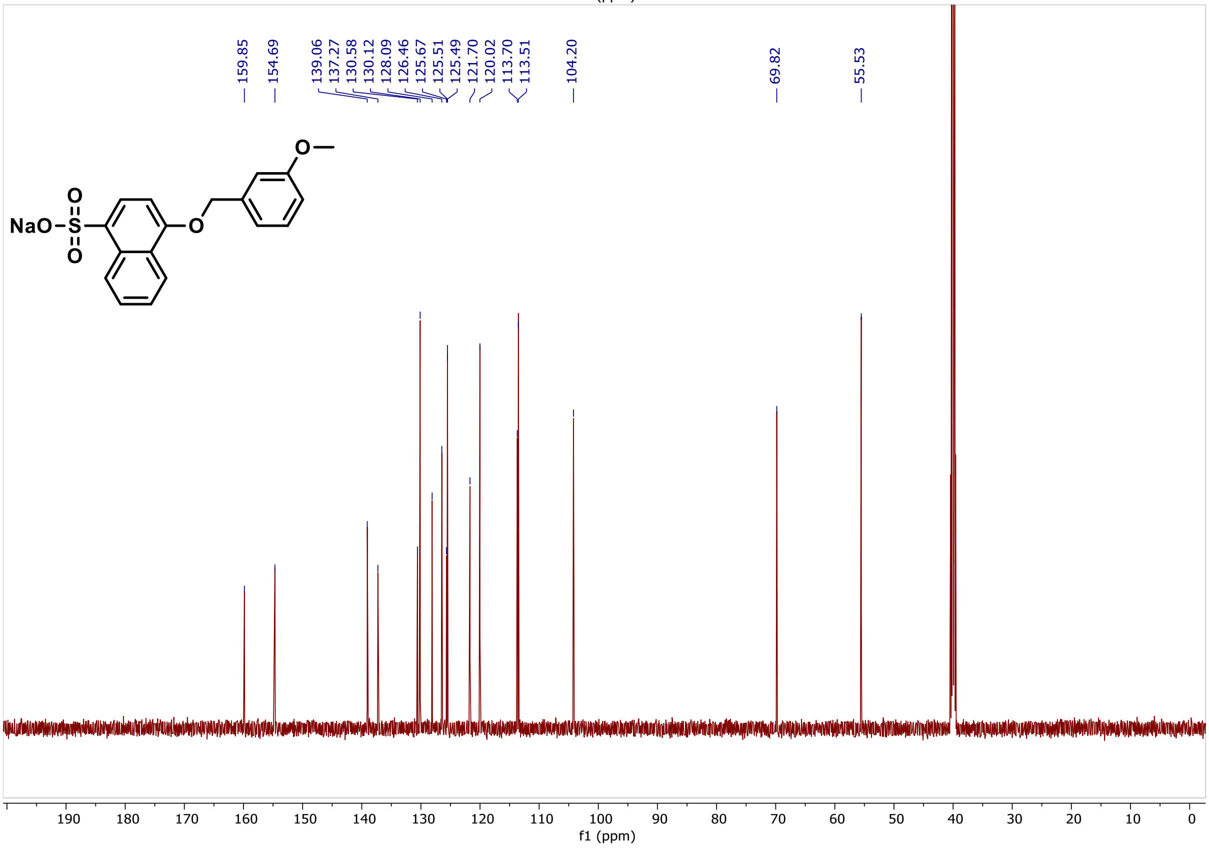
159.85
154.69
139.06
137.27
130.58
130.12
128.09
126.46
125.67
125.51
125.49
121.70
120.02
113.70
113.51
104.20
69.82
55.53
NaO
O
S
O
O
O
f1 (ppm)

**LC/MS, $^{1}$H NMR and $^{13}$C NMR spectra of compound 39**

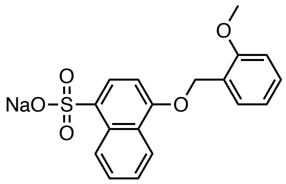


Following the general procedure 4-[(2-methoxyphenyl)methoxy]naphthalene-1-sulfonic acid (5.1 mg, 2%, 0.015 mmol) was isolated as the sodium salt.

$^{1}$H NMR (600 MHz, DMSO) δ 8.79 (d, *J* = 8.4 Hz, 1H), 8.18 (d, *J* = 7.8 Hz, 1H), 7.86 (d, *J* = 8.0 Hz, 1H), 7.56 – 7.49 (m, 2H), 7.49 – 7.44 (m, 1H), 7.38 – 7.33 (m, 1H), 7.09 (d, *J* = 8.2 Hz, 1H), 7.00 (t, *J* = 7.4 Hz, 1H), 6.96 (d, *J* = 8.1 Hz, 1H), 5.27 (s, 2H), 3.85 (s, 3H).

$^{13}$C NMR (151 MHz, DMSO) δ 157.4, 154.9, 137.3, 130.6, 129.8, 129.2, 128.1, 126.4, 125.7, 125.6, 125.4, 125.1, 121.8, 120.9, 111.4, 104.0, 65.4, 56.0.

MS (m/z): $[M\text{-}Na]^-$ 343.0

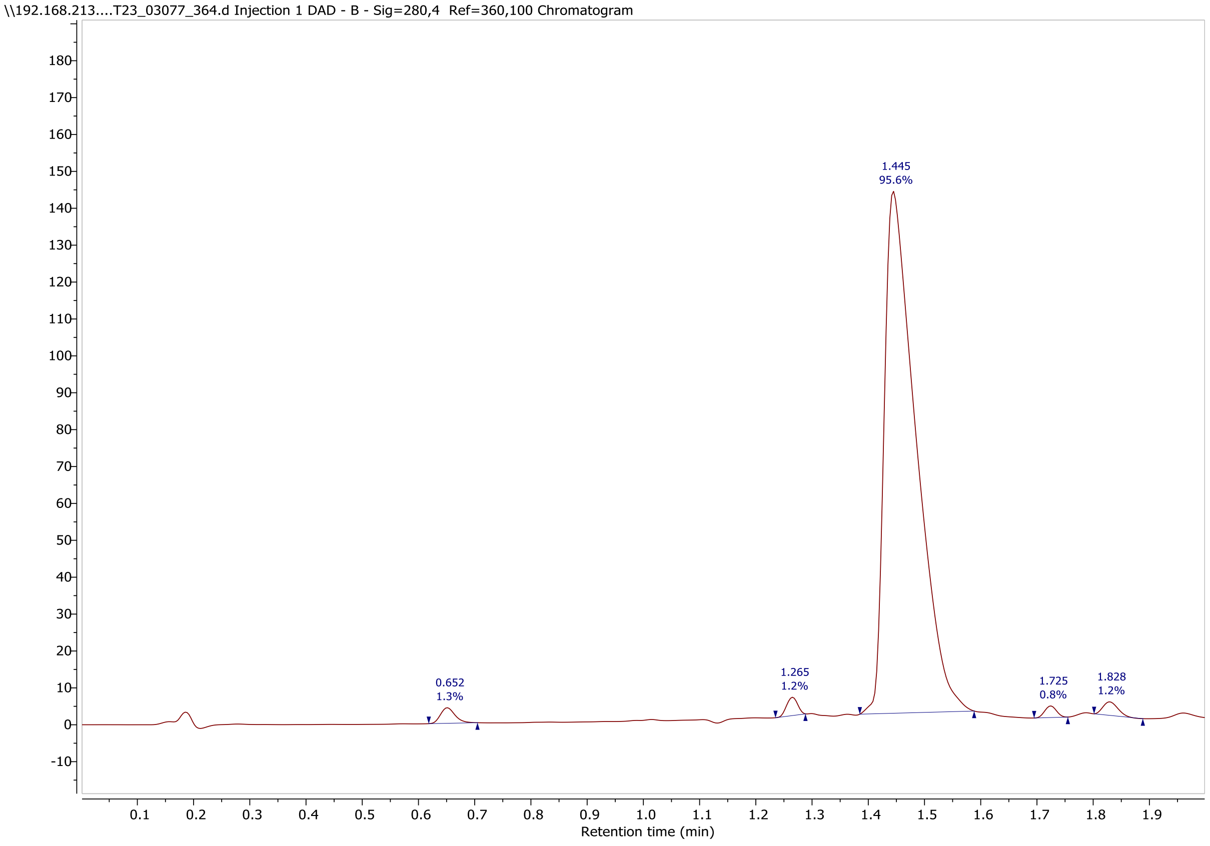

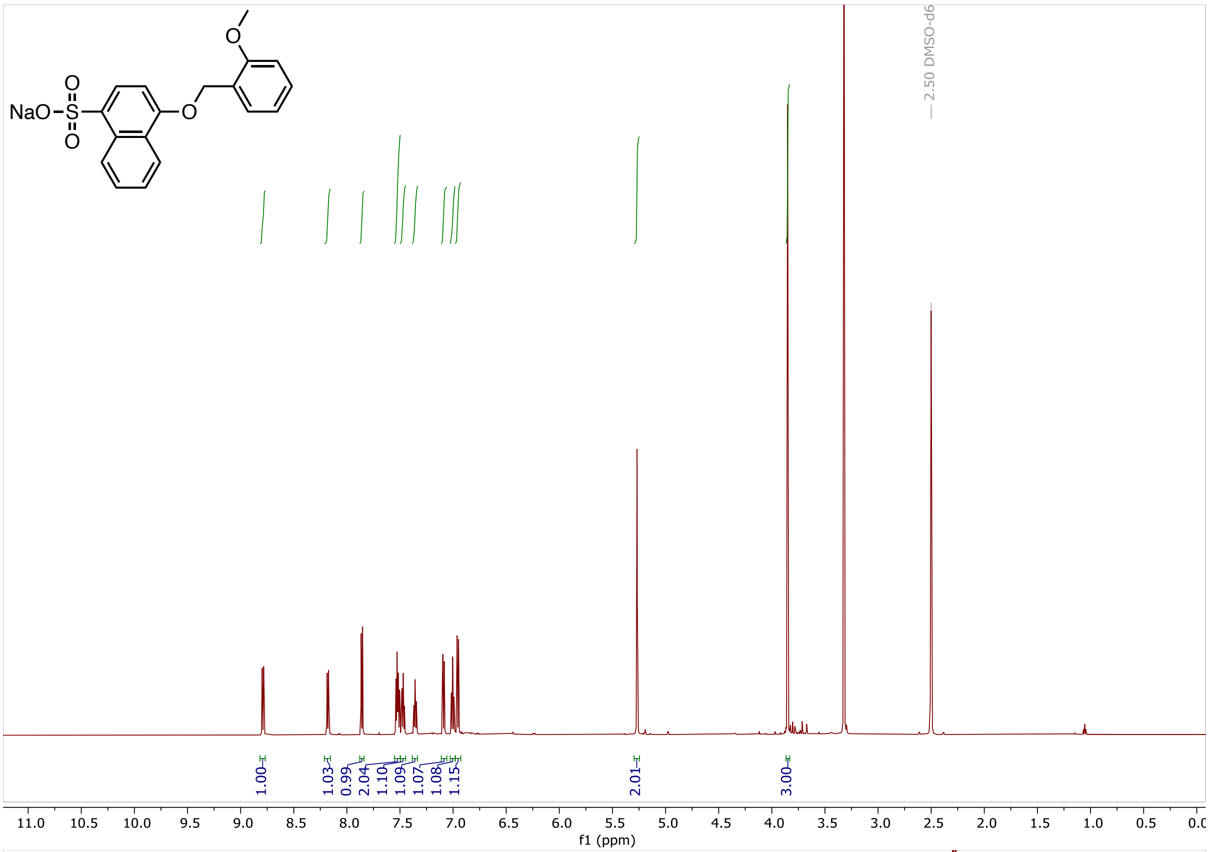
NaO
— 2.50 DMSO-d6
1.00
1.03
0.99
2.04
1.10
1.09
1.07
1.08
1.15
2.01
3.00
f1 (ppm)

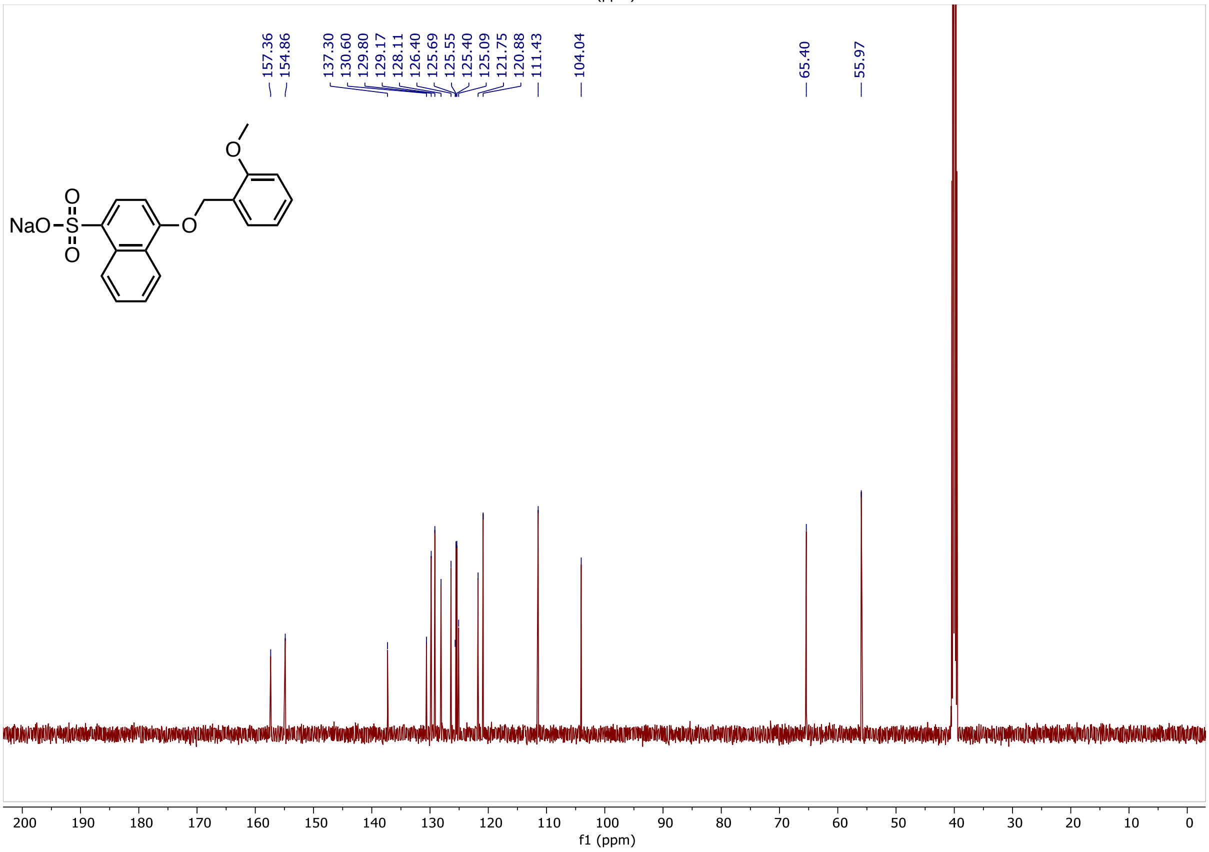
157.36
154.86
137.30
130.60
129.80
129.17
128.11
126.40
125.69
125.55
125.40
125.09
121.75
120.88
111.43
104.04
65.40
55.97
NaO
f1 (ppm)

**LC/MS, [1]H NMR and [13]C NMR spectra of compound 40**

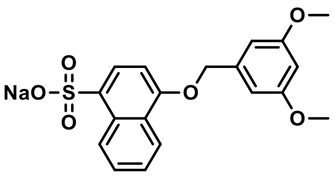


Following the general procedure 4-[(3,5-dimethoxyphenyl)methoxy]naphthalene-1-sulfonic acid (60.8 mg, 17%, 0.162 mmol) was isolated as the sodium salt.

$^{1}$H NMR (600 MHz, DMSO) δ 8.82 – 8.77 (m, 1H), 8.22 – 8.18 (m, 1H), 7.86 (d, $J$ = 8.0 Hz, 1H), 7.52 (dddd, $J$ = 18.9, 8.2, 6.7, 1.5 Hz, 2H), 6.95 (d, $J$ = 8.1 Hz, 1H), 6.71 (d, $J$ = 2.3 Hz, 2H), 6.47 (d, $J$ = 2.3 Hz, 1H), 5.25 (s, 2H), 3.75 (s, 6H).

$^{13}$C NMR (151 MHz, DMSO) δ 160.6, 154.2, 139.4, 136.8, 130.1, 127.6, 126.0, 125.2, 125.1, 125.0, 121.2, 105.3, 103.7, 99.4, 69.4, 55.2.

MS (m/z): [M-Na]$^{-}$ 373.0

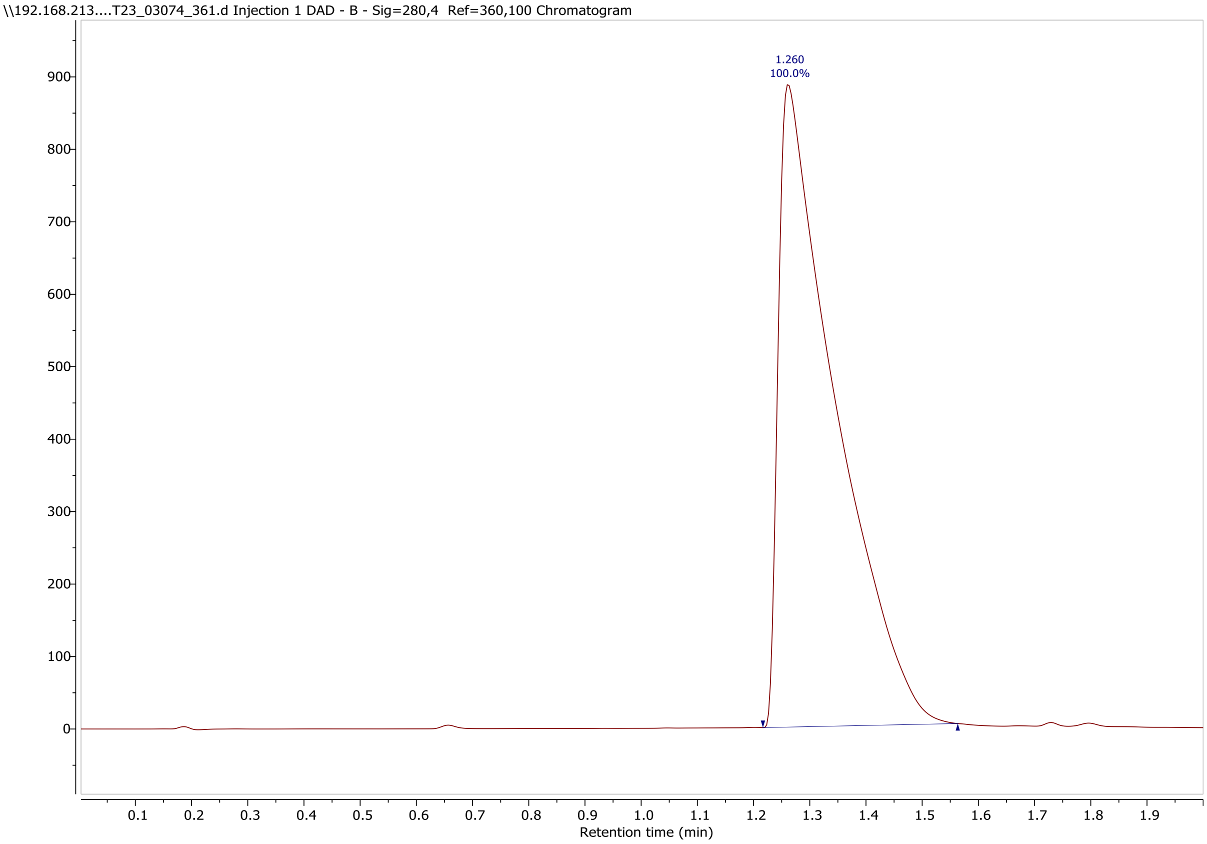

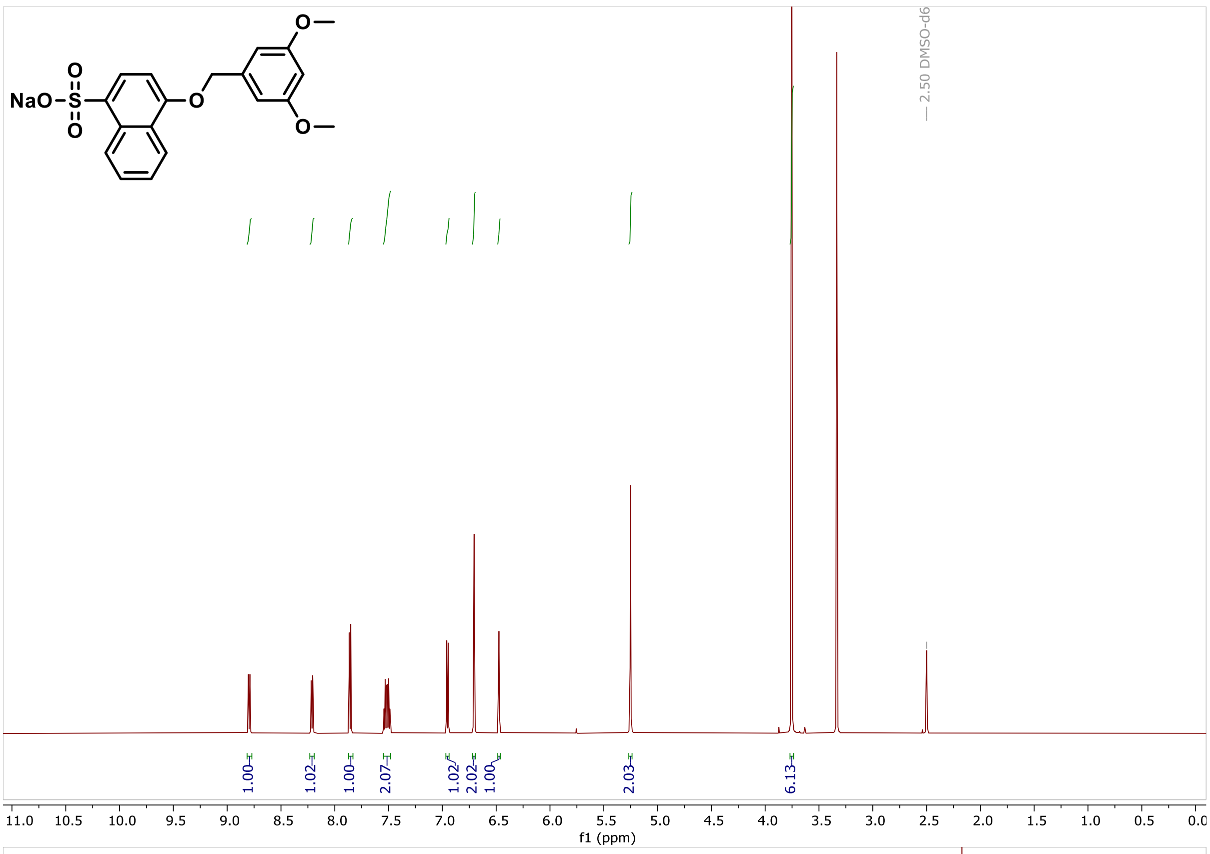
NaO
O
S
O
O
O
O
2.50 DMSO-d6
1.00
1.02
1.00
2.07
1.02
2.02
1.00
2.03
6.13
11.0 10.5 10.0 9.5 9.0 8.5 8.0 7.5 7.0 6.5 6.0 5.5 5.0 4.5 4.0 3.5 3.0 2.5 2.0 1.5 1.0 0.5 0.0
f1 (ppm)

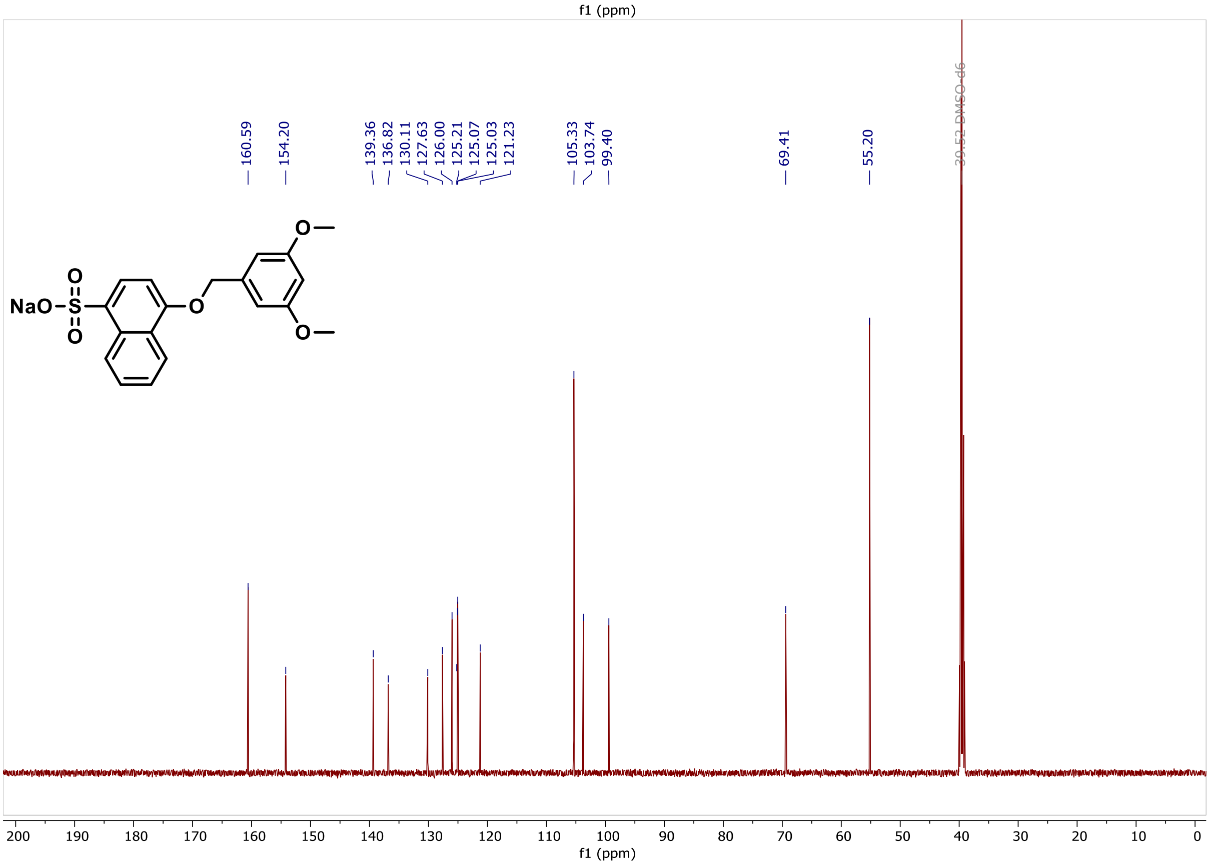
160.59
154.20
139.36
136.82
130.11
127.63
126.00
125.21
125.07
125.03
121.23
105.33
103.74
99.40
69.41
55.20
39.52 DMSO-d6
NaO
O
S
O
O
O
O
200 190 180 170 160 150 140 130 120 110 100 90 80 70 60 50 40 30 20 10 0
f1 (ppm)

**LC/MS, $^{1}$H NMR and $^{13}$C NMR spectra of compound 41**

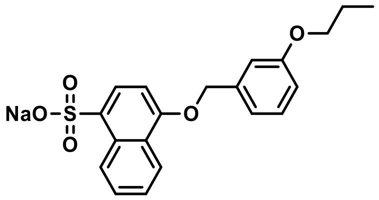


Following the general procedure 4-[(3-propoxyphenyl)methoxy]naphthalene-1-sulfonic acid (4 mg, 1%, 0.011 mmol) was isolated as the sodium salt.

$^{1}$H NMR (600 MHz, DMSO) δ 8.82 – 8.77 (m, 1H), 8.22 – 8.18 (m, 1H), 7.85 (d, *J* = 8.0 Hz, 1H), 7.51 (dddd, *J* = 21.9, 8.1, 6.7, 1.4 Hz, 2H), 7.32 (t, *J* = 8.0 Hz, 1H), 7.14 – 7.08 (m, 2H), 6.96 (d, *J* = 8.1 Hz, 1H), 6.92 – 6.88 (m, 1H), 5.29 (s, 2H), 3.95 (t, *J* = 6.5 Hz, 2H), 1.74 (h, *J* = 7.1 Hz, 2H), 0.98 (t, *J* = 7.4 Hz, 3H).

$^{13}$C NMR (151 MHz, DMSO) δ 159.3, 154.6, 139.1, 137.4, 130.6, 130.1, 128.2, 126.4, 125.7, 125.5, 125.4, 121.7, 119.9, 114.2, 114.0, 104.2, 69.8, 69.4, 22.5, 10.9.

MS (m/z): $[M\text{-}Na]^{-}$ 371.0

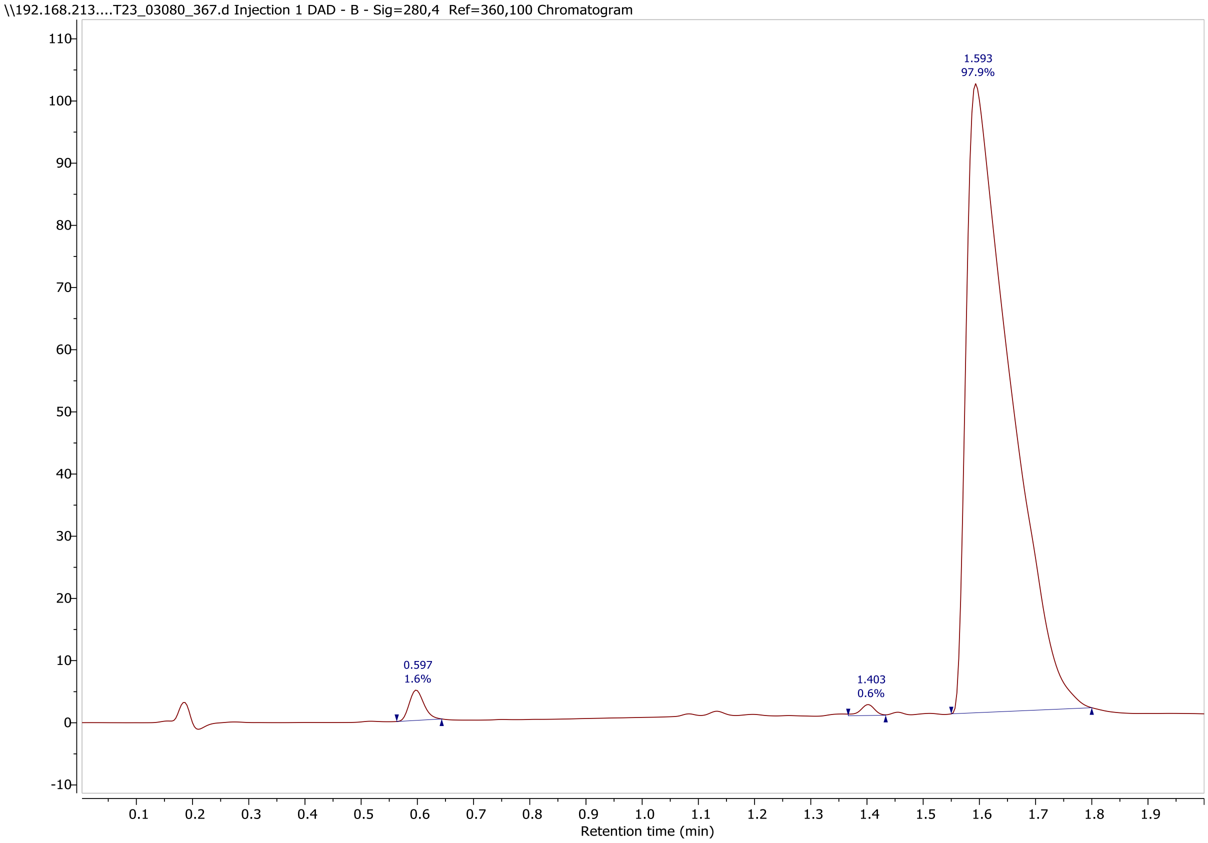

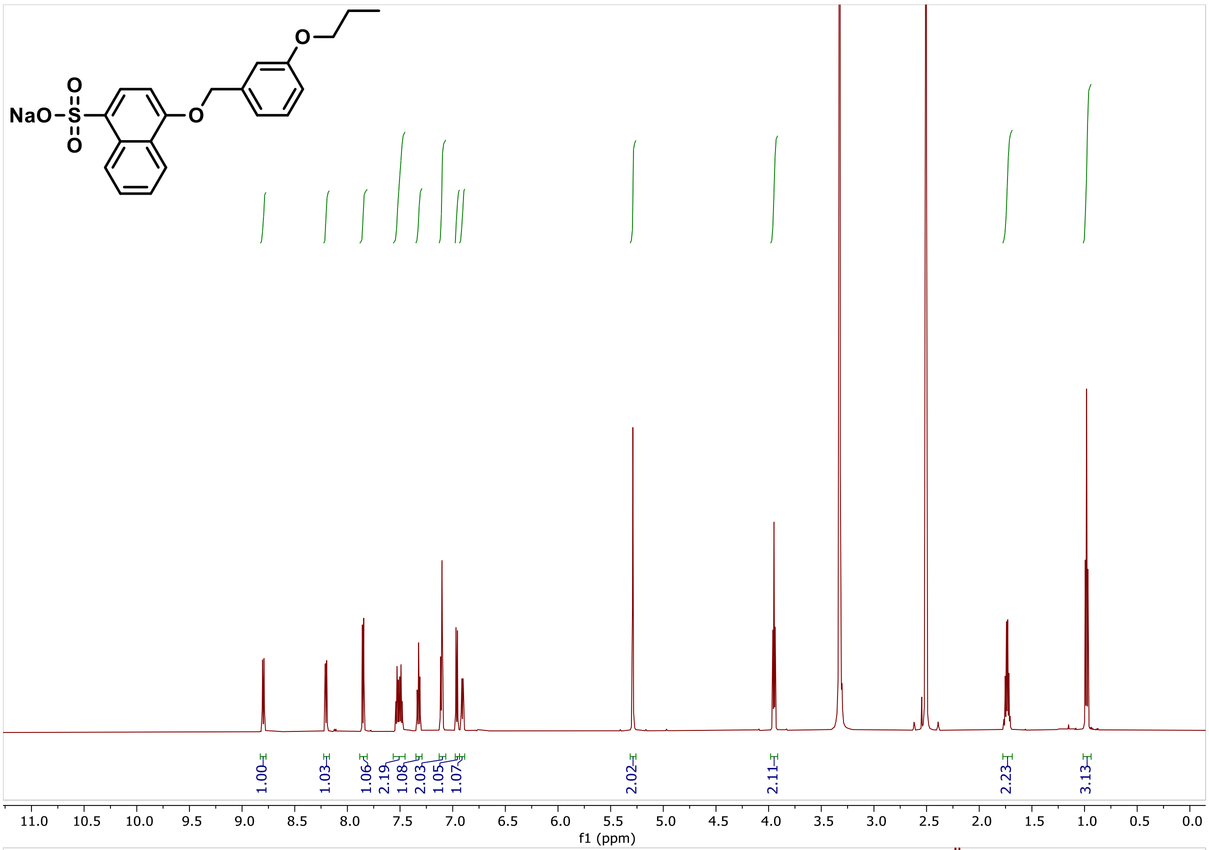
NaO
S
O
O
O
O
1.00
1.03
1.06
2.19
1.08
2.03
1.05
1.07
2.02
2.11
2.23
3.13
11.0
10.5
10.0
9.5
9.0
8.5
8.0
7.5
7.0
6.5
6.0
5.5
5.0
4.5
4.0
3.5
3.0
2.5
2.0
1.5
1.0
0.5
0.0
f1 (ppm)

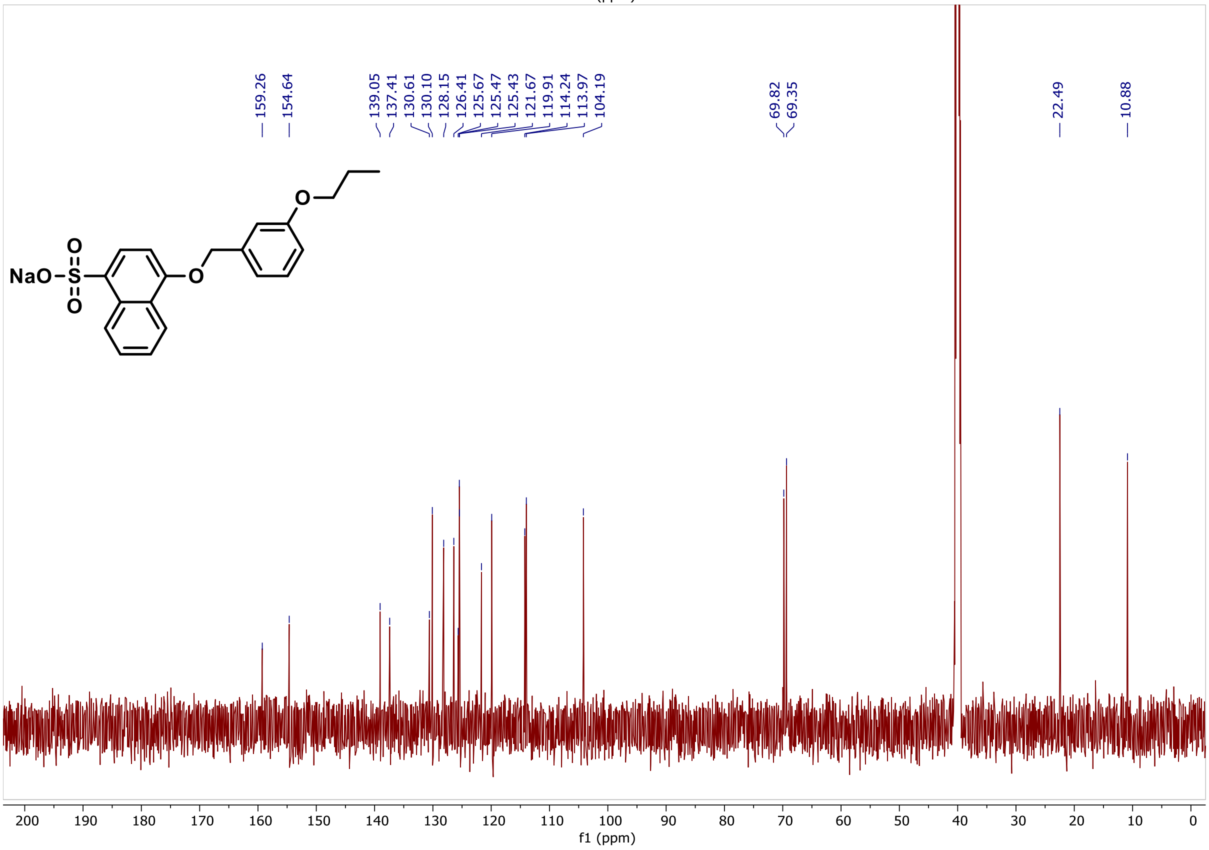
159.26
154.64
139.05
137.41
130.61
130.10
128.15
126.41
125.67
125.47
125.43
121.67
119.91
114.24
113.97
104.19
69.82
69.35
22.49
10.88
NaO
S
O
O
O
O
200
190
180
170
160
150
140
130
120
110
100
90
80
70
60
50
40
30
20
10
0
f1 (ppm)

**LC/MS, $^{1}$H NMR and $^{13}$C NMR spectra of compound 42**

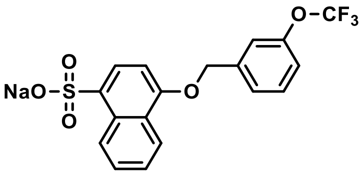


Following the general procedure 4-[[3-(trifluoromethoxy)phenyl]methoxy]naphthalene-1-sulfonic acid (93.1 mg, 25%, 0.234 mmol) was isolated as the sodium salt.

$^{1}$H NMR (600 MHz, DMSO) δ 8.82 – 8.76 (m, 1H), 8.21 (dd, *J* = 8.2, 1.6 Hz, 1H), 7.87 (d, *J* = 8.0 Hz, 1H), 7.63 – 7.46 (m, 5H), 7.38 – 7.32 (m, 1H), 6.98 (d, *J* = 8.1 Hz, 1H), 5.39 (s, 2H).

$^{13}$C NMR (151 MHz, DMSO) δ 154.5, 149.0, 140.4, 137.5, 131.1, 130.6, 128.1, 126.9, 126.5, 125.6, 125.4, 121.6, 120.8, 120.3, 104.3, 69.0.

$^{19}$F NMR (471 MHz, DMSO) δ -56.66.

MS (m/z): [M-Na]$^{-}$ 397.0

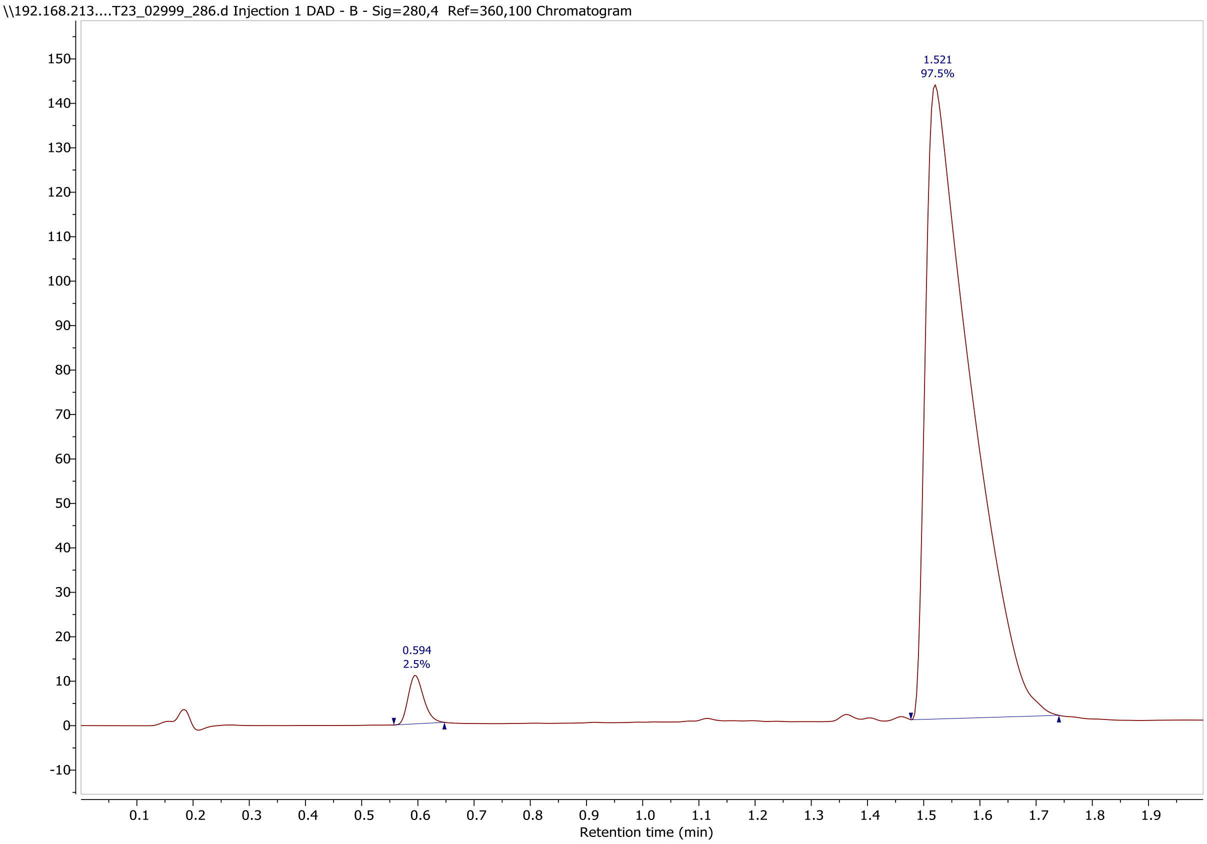

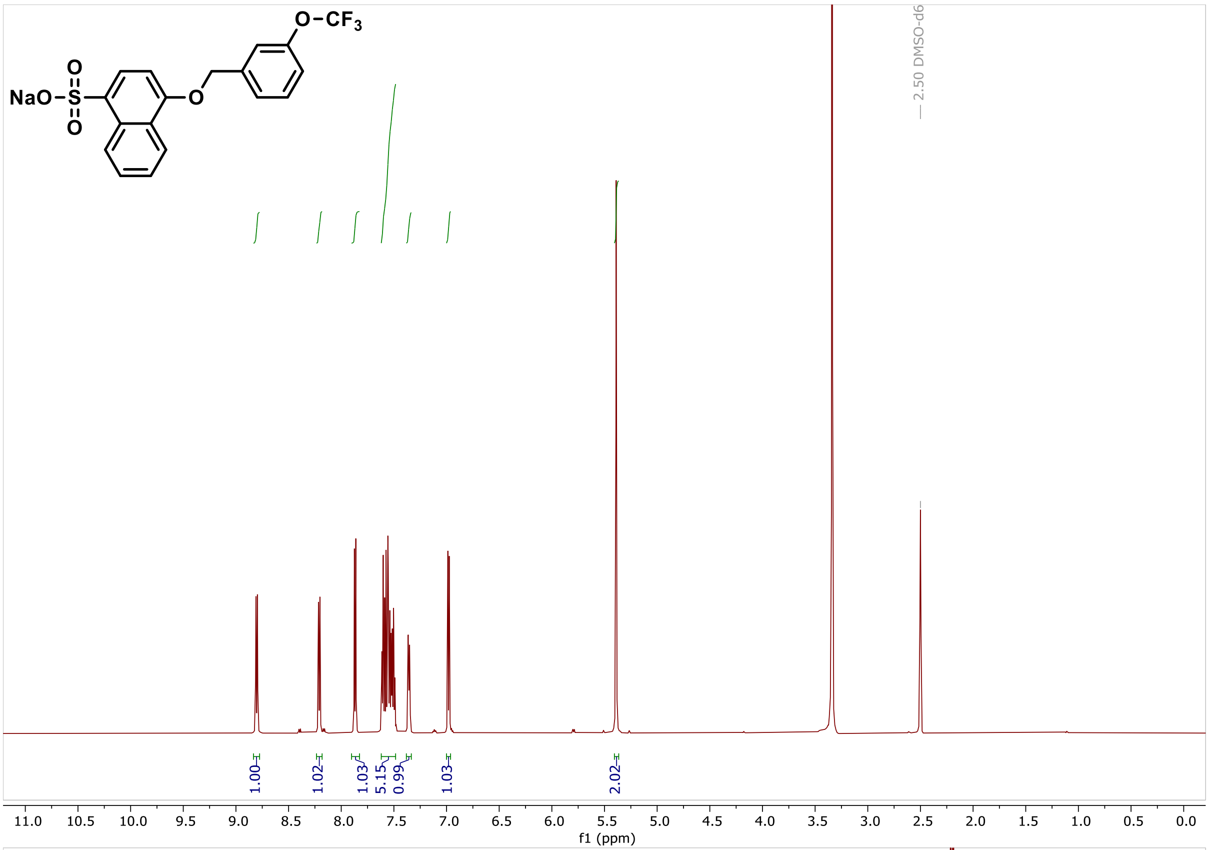

O−CF3
NaO−S−O
O
O
2.50 DMSO-d6
1.00
1.02
1.03
5.15
0.99
1.03
2.02
11.0 10.5 10.0 9.5 9.0 8.5 8.0 7.5 7.0 6.5 6.0 5.5 5.0 4.5 4.0 3.5 3.0 2.5 2.0 1.5 1.0 0.5 0.0
f1 (ppm)


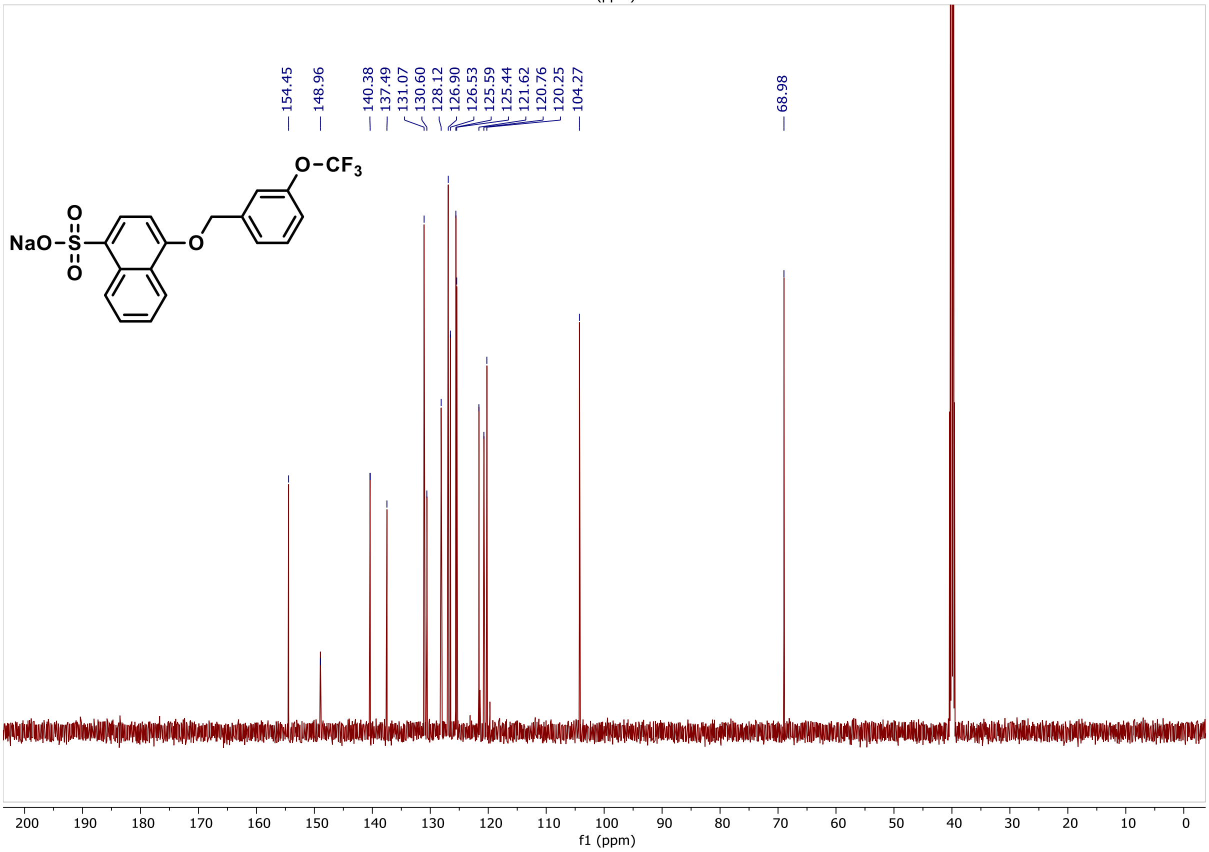

154.45
148.96
140.38
137.49
131.07
130.60
128.12
126.90
126.53
125.59
125.44
121.62
120.76
120.25
104.27
68.98
O−CF3
NaO−S−O
O
O
200 190 180 170 160 150 140 130 120 110 100 90 80 70 60 50 40 30 20 10 0
f1 (ppm)

**LC/MS, $^{1}$H NMR and $^{13}$C NMR spectra of compound 43**

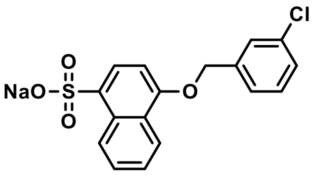


Following the general procedure 4-[(3-chlorophenyl)methoxy]naphthalene-1-sulfonic acid (98.3 mg, 30%, 0.282 mmol) was isolated as the sodium salt.

$^{1}$H NMR (600 MHz, DMSO) δ 8.83 – 8.77 (m, 1H), 8.23 – 8.17 (m, 1H), 7.86 (d, *J* = 8.0 Hz, 1H), 7.62 (t, *J* = 1.9 Hz, 1H), 7.56 – 7.48 (m, 3H), 7.46 (t, *J* = 7.8 Hz, 1H), 7.42 (dt, *J* = 8.2, 1.5 Hz, 1H), 6.97 (d, *J* = 8.1 Hz, 1H), 5.34 (s, 2H).

$^{13}$C NMR (151 MHz, DMSO) δ 154.5, 140.1, 137.5, 133.6, 131.0, 130.6, 128.3, 128.1, 127.7, 126.5, 126.5, 125.6, 125.6, 125.4, 121.7, 104.2, 69.0.

MS (m/z): $[M\text{-}Na]^{-}$ 347.0

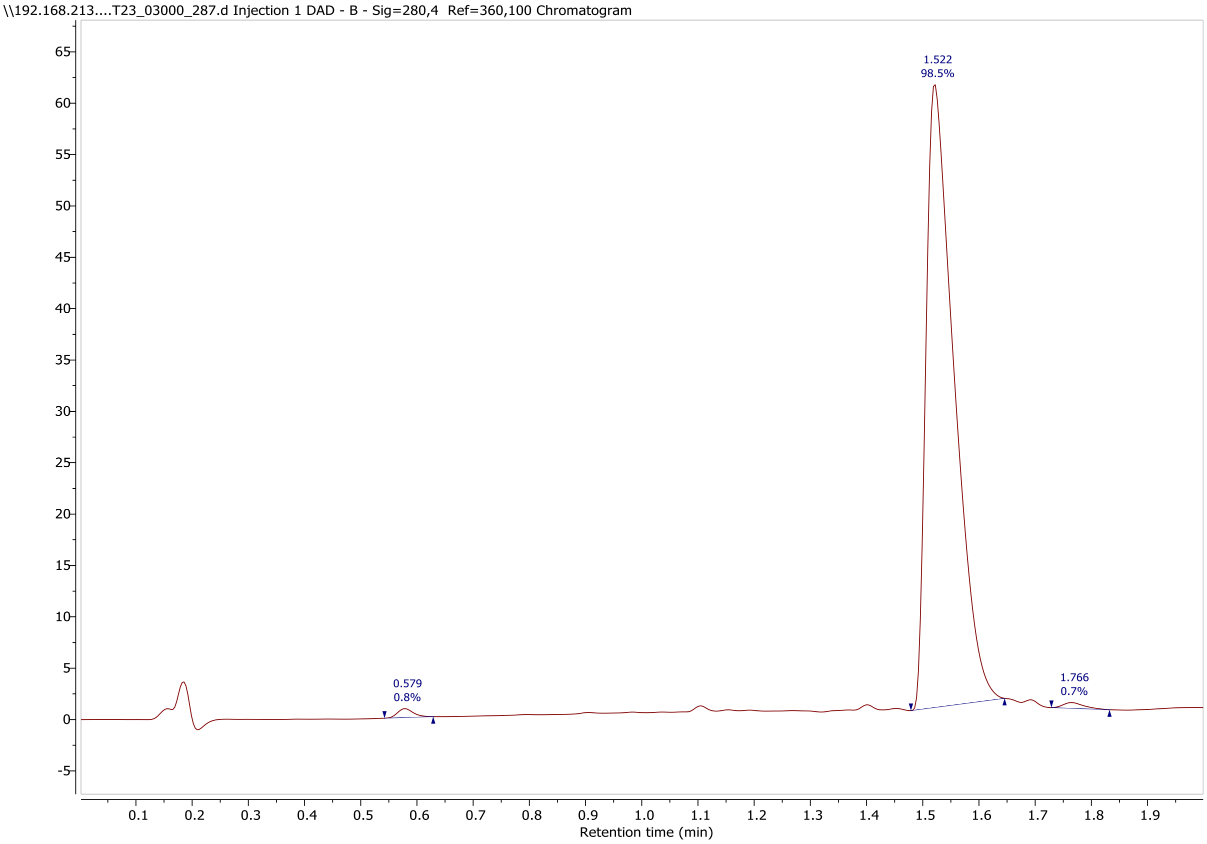

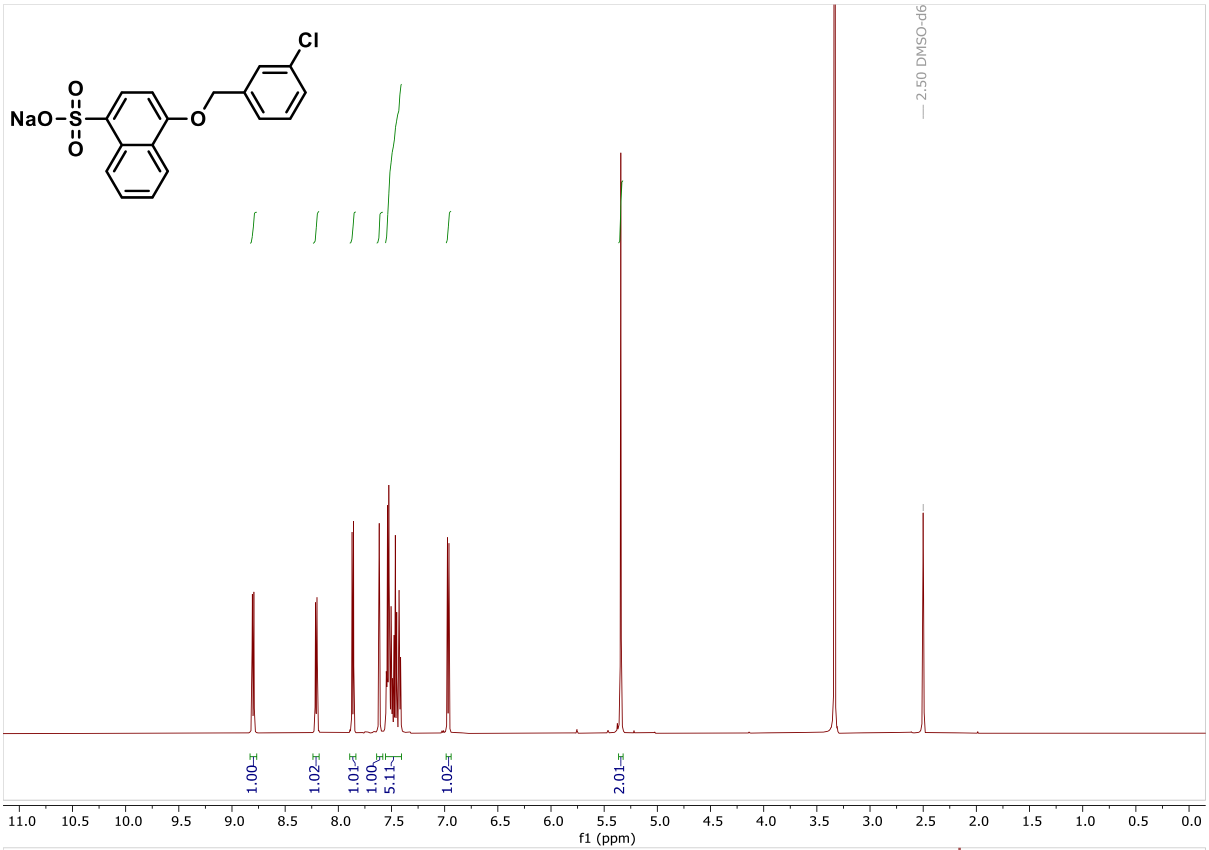

Cl
NaO
S
O
O
O
2.50 DMSO-d6
1.00
1.02
1.01
1.00
5.11
1.02
2.01
11.0 10.5 10.0 9.5 9.0 8.5 8.0 7.5 7.0 6.5 6.0 5.5 5.0 4.5 4.0 3.5 3.0 2.5 2.0 1.5 1.0 0.5 0.0
f1 (ppm)


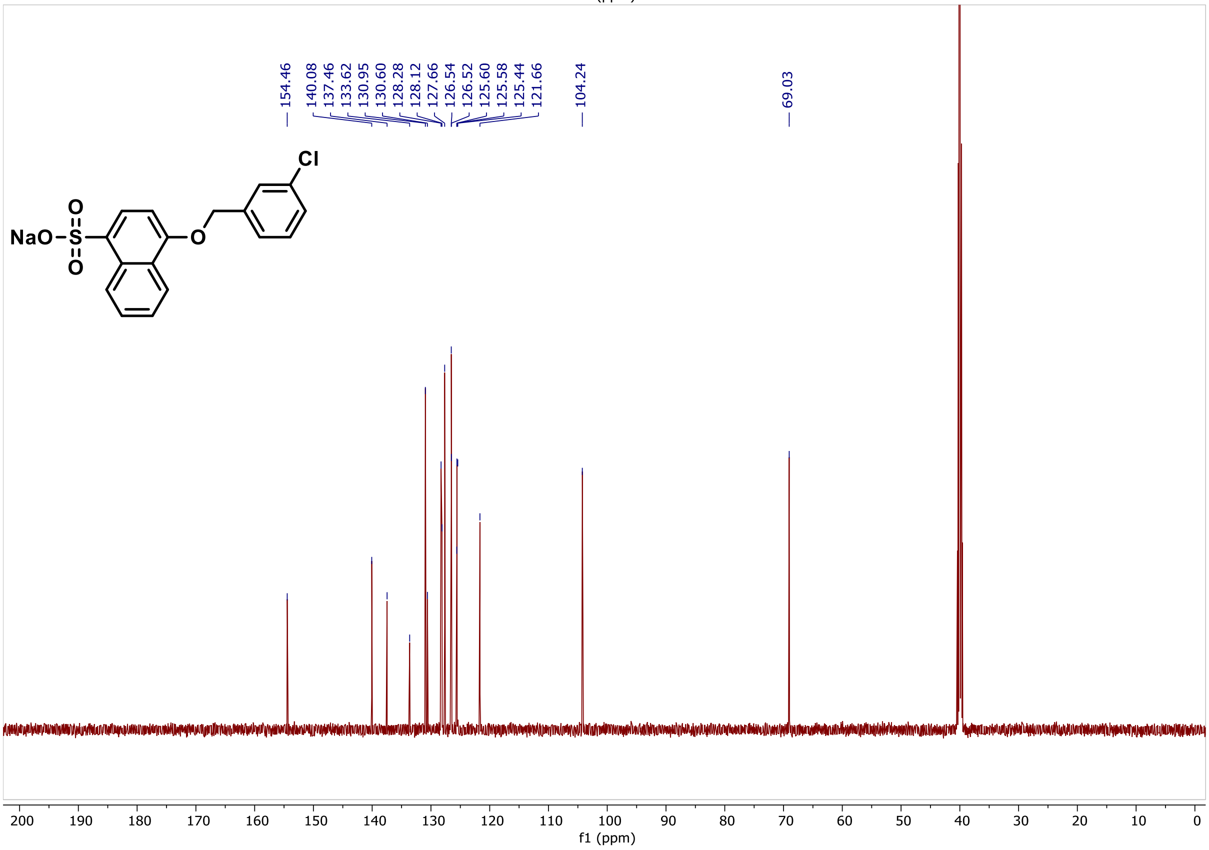

154.46
140.08
137.46
133.62
130.95
130.60
128.28
128.12
127.66
126.54
126.52
125.60
125.58
125.44
121.66
104.24
69.03
Cl
NaO
S
O
O
O
200 190 180 170 160 150 140 130 120 110 100 90 80 70 60 50 40 30 20 10 0
f1 (ppm)

**LC/MS, $^{1}$H NMR and $^{13}$C NMR spectra of compound 44**

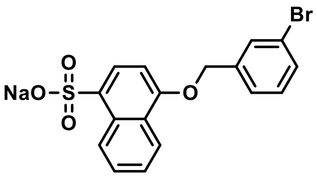


Following the general procedure 4-[(3-bromophenyl)methoxy]naphthalene-1-sulfonic acid (136.6 mg, 37%, 0.347 mmol) was isolated as the sodium salt.

$^{1}$H NMR (600 MHz, DMSO) δ 8.84 – 8.79 (m, 1H), 8.24 – 8.18 (m, 1H), 7.88 (d, *J* = 8.0 Hz, 1H), 7.76 (t, *J* = 1.9 Hz, 1H), 7.59 – 7.49 (m, 4H), 7.40 (t, *J* = 7.8 Hz, 1H), 6.97 (d, *J* = 8.1 Hz, 1H), 5.34 (s, 2H).

$^{13}$C NMR (151 MHz, DMSO) δ 154.5, 140.3, 137.4, 131.3, 131.2, 130.6, 130.6, 128.1, 126.9, 126.5, 125.6, 125.5, 122.2, 121.7, 104.3, 69.0.

MS (m/z): $[M\text{-}Na]^{-}$ 390.9

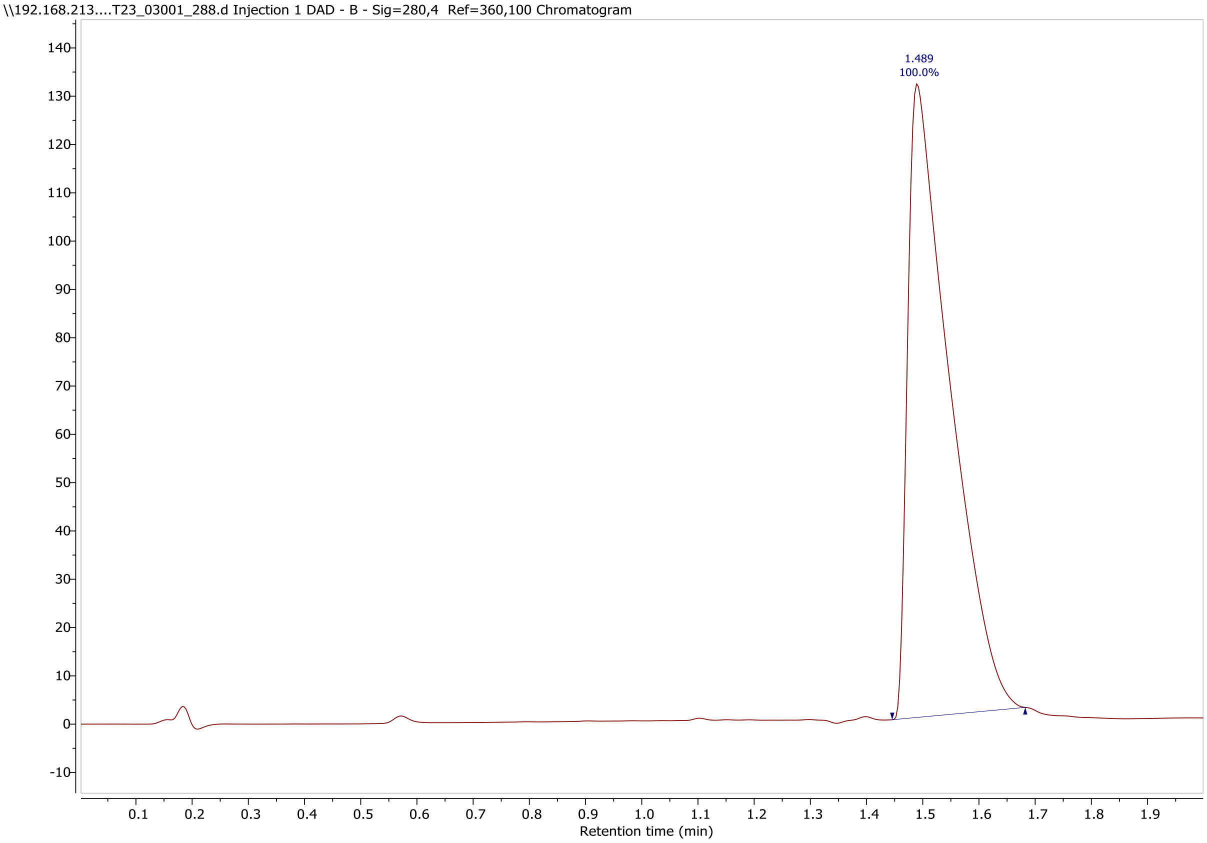

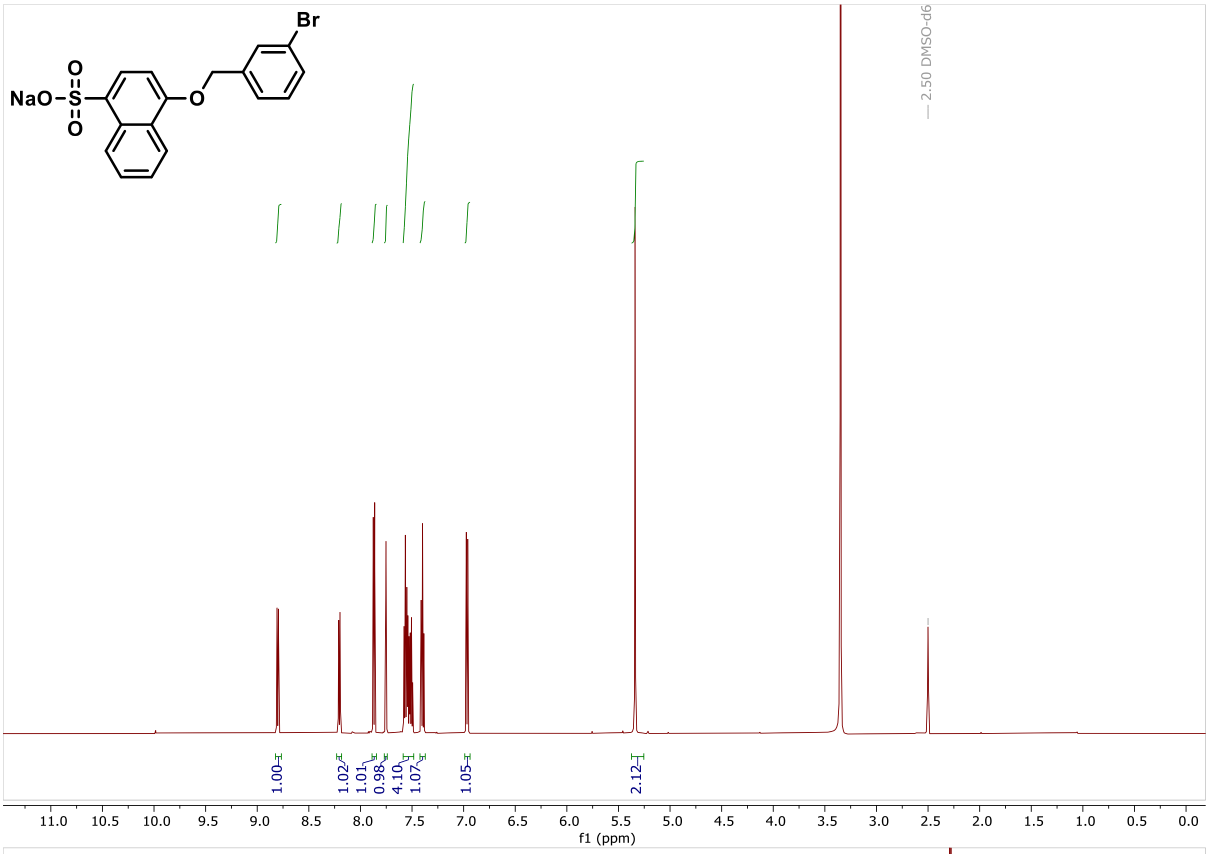
Br
NaO
S
O
O
O
2.50 DMSO-d6
1.00
1.02
1.01
0.98
4.10
1.07
1.05
2.12
f1 (ppm)

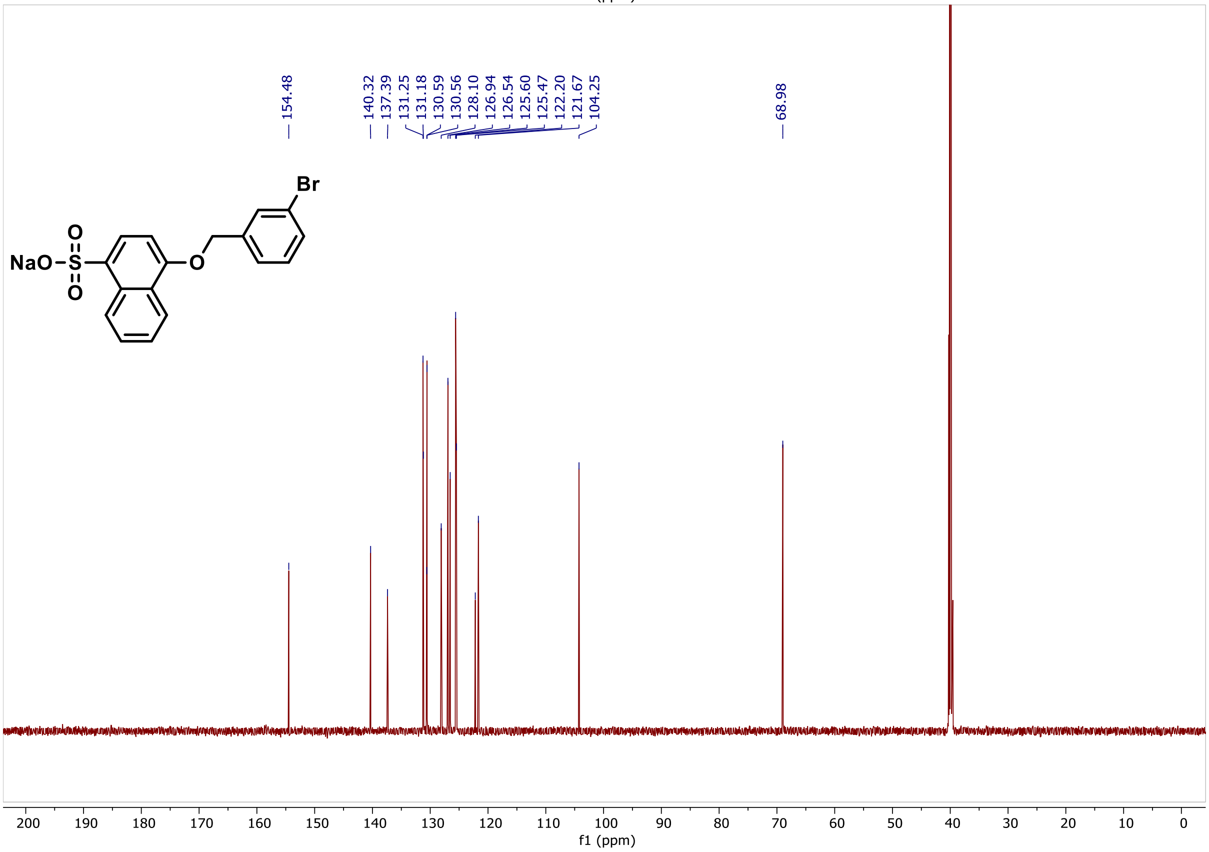
154.48
140.32
137.39
131.25
131.18
130.59
130.56
128.10
126.94
126.54
125.60
125.47
122.20
121.67
104.25
68.98
Br
NaO
S
O
O
O
f1 (ppm)
